# Physics-driven self-supervised learning for quantitative high-fidelity structured illumination microscopy

**DOI:** 10.64898/2026.06.12.731778

**Authors:** Yujun Tang, Zewei Luo, Xi Zhu, Wenyi Wang, Xichuan Ge, Meiqi Li, Cong Chen, Tongsheng Chen, Ceshi Chen, Peng Xi, Gang Wen

## Abstract

Structured illumination microscopy (SIM) enables rapid, long-term super-resolution (SR) imaging of live-cell dynamics. However, current SIM reconstruction methods, although achieving high structural fidelity, generally lack reliable intensity quantification. Here we present qHiFi-SIM, a physics-driven self-supervised learning framework for quantitative high-fidelity SIM imaging. By leveraging the wide-field (WF) image as a physical intensity reference and conventional SIM reconstruction as a structural continuity prior, qHiFi-SIM enables self-supervised training without ground-truth intensity data. It achieves structural fidelity comparable to HiFi-SIM while maintaining excellent intensity linearity at the SIM point-spread function scale (R^2^ > 0.986, mean intensity ratio near 1). The framework transfers across diverse SIM setups and typical samples, and directly improves the intensity linearity of other state-of-the-art algorithms without retraining. We demonstrate its utility for live-cell quantitative visualization of mitochondrial intensity dynamics and high-fidelity SIM-FRET functional imaging, offering a practical tool for SR structural visualization and quantitative functional imaging in live cells.

## Introduction

Owing to the inherent transparency of subcellular organelles, fluorescence labeling has emerged as the dominant approach for probing their localization and abundance. Quantitative fluorescence microscopy enables accurate detection of signal concentration, which underlies its widespread use across numerous methodologies^1,2^, with representative examples including fluorescence correlation spectroscopy (FCS)^3^, Förster resonance energy transfer (FRET)^4^, fluorescence recovery after photobleaching (FRAP)^5^, fluorescence lifetime imaging microscopy (FLIM)^6^, and spectral unmixing^7^. Super-resolution (SR) fluorescence microscopy provides nanoscale insights into biological processes, serving as an essential tool for biomedical research, particularly in live-cell imaging^8,9^. Yet current SR methods focus predominantly on structural imaging, while their capability for quantitative intensity imaging remains substantially limited. Among various SR techniques, structured illumination microscopy (SIM) stands out as the preferred tool for long-term live-cell dynamic imaging due to its rapid acquisition speed, high photon efficiency, low phototoxicity, and excellent compatibility with fluorescent labels^10,11^. SIM typically acquires multiple images under varying illumination patterns and reconstructs them to achieve up to a twofold resolution enhancement over conventional wide-field (WF) microscopy. As a computational imaging method, the reliability and practicality of SIM critically depend on the structural fidelity and intensity quantification of the reconstructed SR images. Nowadays, numerous advanced SIM reconstruction algorithms, such as Hessian-SIM^12^, HiFi-SIM^13^, noise-controlled SIM^14^, Sparse-SIM^15^, direct-SIM^16^, PCA-SIM^17^, BF-SIM^18^, FlexSIM^19^, Lock-in-SIM^20^, and deep-learning-based approaches^21-28^, have been developed to effectively suppress typical reconstruction artifacts, enabling high-fidelity resolution of fine specimen structures. Nevertheless, the intensity quantification of SIM images remains underexplored, which substantially restricts the development and application of quantitative SR imaging in live cells.

Standard SIM reconstruction follows the generalized Wiener deconvolution procedure (hereinafter referred to as Wiener-SIM), which theoretically guarantees structural interpretability and intensity quantification in SR-SIM images^29-31^ (**Supplementary Note 1**). In practice, however, various factors such as out-of-focus background, noise, mismatched point spread function (PSF), errors in estimated pattern parameters, and suboptimal user-defined parameters frequently introduce artifacts^12-20^. Consequently, advanced SIM algorithms incorporating various strategies, including background removal^18,20,32^, spectrum optimization^13,14,16^, and regularized deconvolution^12,15,33-35^, have been developed to suppress typical artifacts. Yet most of these operations nonlinearly distort the intensity linearity of SR-SIM reconstructions (**Supplementary Note 1**), and such distortions are difficult to characterize explicitly. Moreover, improper operations during reconstruction, such as frequency-domain zero-padding and improper normalization (**Supplementary Note 2; Supplementary Figs. 1 and 2**), can introduce deviations between the reconstructed intensity and the underlying sample distribution. Although recent computational advances have sought to improve intensity quantification in SIM^18,36,37^, this long-standing challenge remains unresolved.

Deep learning-based SR (DLSR) approaches have achieved substantial success by learning end-to-end image transformations from large-scale datasets, independent of explicit analytical models^21-23,26,27^. Yet such approaches, particularly within supervised learning frameworks, typically require large sets of paired low-quality (or low-resolution) inputs and corresponding high-quality ground truth (GT) images. Moreover, they frequently exhibit limited transferability across imaging systems and generalize poorly to different sample structures. Consequently, when applied to intensity-quantitative SIM reconstruction, existing supervised DLSR strategies encounter a fundamental obstacle: the absence of experimentally accessible GT intensity information from biological specimens at the SR scale. To date, investigations into the reconstruction intensity linearity of DLSR methods remain scarce. These bottlenecks have long hindered the development of quantitative SR-SIM technology, leaving a critical gap in the field and limiting its application in cutting-edge biomedical research.

Here we present qHiFi-SIM, a physics-driven self-supervised DL framework that simultaneously achieves high-fidelity reconstruction of SR structures and quantitative preservation of intensity linearity, a combination that current SOTA SIM methods fail to deliver. The model is trained using the equivalent WF image generated from raw SIM data as a physical intensity reference, together with the SR image reconstructed by our previously established HiFi-SIM algorithm^13^ as a structural prior. This strategy effectively suppresses common reconstruction artifacts while eliminating the need for inaccessible GT intensity labels. Through comprehensive simulations and rigorous experimental validation, we demonstrate that qHiFi-SIM achieves structural fidelity and spatial resolution gain comparable to those of the HiFi-SIM algorithm, while maintaining excellent intensity linearity, with R^2^ > 0.986 between degraded and experimental WF intensities, and mean intensity ratio near 1 at the SIM PSF scale. The framework also exhibits good transferability across different SIM setups and sample types, and can be directly applied to improve the intensity linearity of mainstream advanced SIM algorithms (e.g., fairSIM^32^, BF-SIM^18^, FlexSIM^19^, and TV-based^34^ and Hessian-based^12^ denoising) without retraining. We further demonstrate its advantages for quantitative visualization of mitochondrial nanoscale structures and fluorescence intensity dynamics in live cells, as well as high-fidelity SIM-FRET functional imaging that demands heightened intensity quantification. To facilitate broad accessibility by both the SIM research community and biological researchers, we provide an easy-to-use software package aimed at accelerating the integration of quantitative SR-SIM into routine imaging practices.

## Results

### Development of qHiFi-SIM

An ideal quantitative SIM technique should achieve artifact-free SR reconstruction of fine structures while preserving fluorescence intensity linearity at the SIM PSF scale across the entire field-of-view (FoV). Yet, despite the structural fidelity provided by advanced SIM algorithms, maintaining reliable intensity linearity at this scale remains a challenge. To this end, we developed qHiFi-SIM, a physics-driven self-supervised DL framework for intensity-quantitative SIM imaging (**Fig. 1a,b; Supplementary Fig. 3**). Our approach is grounded in two complementary physical constraints: photon-count conservation at the WF PSF scale, which governs the total photon budget, and biological structural continuity at the SIM PSF scale, which constrains the spatial allocation of photon counts. During SIM acquisition, averaging raw images across all phase shifts and orientations yields an experimental WF image that provides quantitatively accurate intensity information within the diffraction-limited cutoff frequency (*k*_c_), whose intensity is related to the fluorophore distribution of the sample by convolution with the system PSF^13^

**Fig. 1.**
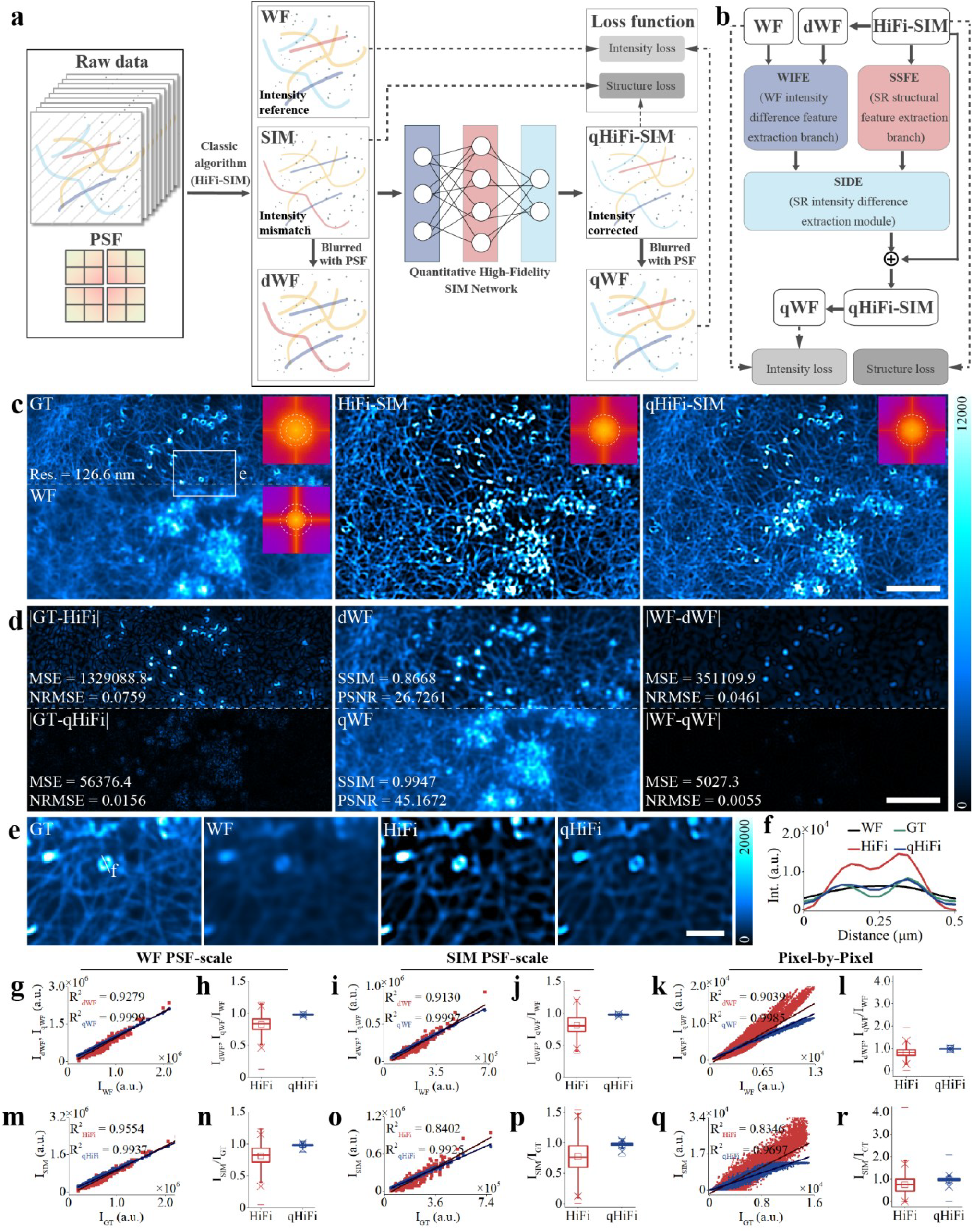
Principle of qHiFi-SIM and simulation validation. **a**, Reconstruction workflow. **b**, Network architecture comprising dual-branch feature extraction, SIDE fusion module, and identity skip connection. **c**, Representative simulation results. The GT was obtained by applying HiFi-SIM reconstruction to F-actin data from the BioSR dataset; SIM imaging was simulated using parameters matching those of the dataset. Shown are the GT (upper left), WF image (lower left), HiFi-SIM reconstruction (middle), and qHiFi-SIM reconstruction (right). **d**, Left: error maps of the HiFi-SIM and qHiFi-SIM reconstructions relative to the GT. Middle: degraded WF images dWF (top) and qWF (bottom), generated by convolving the corresponding SR images in **c** with the theoretical PSF. Right: error maps of the degraded WF images relative to the experimental WF. **e**, Magnified views of the white-boxed region in **c. f**, Fluorescence intensity profiles along the white line in **e. g–l**, Quantitative comparison of degraded WF images (dWF and qWF) against the experimental WF across three spatial scales: WF PSF scale (**g**,**h**), SIM PSF scale (**i,j**), and single-pixel scale (**k,l**). R^2^ values (**g,i,k**) and box plots of intensity ratios (**h,j,l**) are derived from 500 randomly selected ROIs at each scale. **m–r**, Quantitative comparison of SIM reconstructions (HiFi-SIM and qHiFi-SIM) against GT across the same three spatial scales: WF PSF scale (**m,n**), SIM PSF scale (**o,p**), and single-pixel scale (**q,r**). R^2^ values (**m,o,q**) and box plots of intensity ratios (**n,p,r**) are derived from 500 randomly selected ROIs at each scale. In the box plots, the center line and open square represent the median and mean, respectively. The box edges denote the 25th and 75th percentiles, while the whiskers extend to the 5th and 95th percentiles. Maximum and minimum values are marked by crosses, and the 99% and 1% percentiles are indicated by dashes. Scale bars: **c, d** 4 μm; **e** 1 μm.

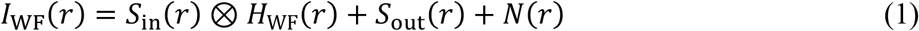

where *I*_WF_ is the experimental WF image; *S*_in_ is the in-focus sample signal, ⊗ indicates the convolution operation, *H*_WF_ is the system PSF, *S*_out_ denotes the out-of-focus background, and *N* accounts for noise signal. Since *H*_WF_ is normalized (∬ *H*_WF_ = 1), the convolution in Eq. (**1**) conserves the total photon count, rendering *I*_WF_ a physically accurate intensity reference within the diffraction limit. However, *I*_WF_ specifies only the photon budget within each WF PSF-scale region, not how these photons are distributed among individual pixels; therefore, recovering the SIM PSF-scale intensity from *I*_WF_ alone is ill-posed. We resolve this ambiguity with a second physical prior, biological structural continuity, provided by the HiFi-SIM reconstruction (**Fig. 1a,b**): although its absolute intensities may deviate, its structural distribution is positionally reliable^13^, indicating where, rather than how many, photons reside. The two priors are complementary by construction: the WF image fixes the photon budget of each region, while the HiFi-SIM image constrains the photon allocation within it. Building on these two priors, qHiFi-SIM reconstruction can be formulated as a constrained optimization problem. Among all candidate intensity distributions *I*_SIM_ satisfying the WF photon-count constraint, the optimal solution is obtained by minimizing a regularization functional *R*(*I*_SIM_) that encodes the structural continuity prior^38,39^

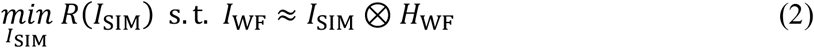

where the symbol ≈ denotes that the residual *I*_SIM_ ⊗ *H*_WF_−*I*_WF_ is measured in the L1 sense, tolerating the background and noise terms in Eq. (**1**). For a strictly convex *R*(·), as is standard in classical inverse problems^38,39^, Eq. (**2**) admits a unique global minimizer. Essentially, Eq. (**2**) states that the intensity at the SIM PSF scale is not a free fitting variable; it is jointly constrained by photon conservation and structural continuity.

Solving Eq. (**2**) explicitly for every dataset is computationally challenging. qHiFi-SIM therefore trains a neural network to approximate its solution^39-41^ (**Fig. 1a,b**). The fidelity of this approximation is established experimentally against known GT (**Fig. 1c–r**). The network adopts a dual-branch architecture for feature extraction (**Methods; Supplementary Fig. 3**). The first branch, termed SR structural feature extraction (SSFE), extracts high-fidelity structure details from the HiFi-SIM input, preventing resolution degradation (**Supplementary Fig. 4; Supplementary Table 1**). The second branch, termed WF intensity difference feature extraction (WIFE), processes the experimental WF reference (denoted WF) alongside a degraded WF image (denoted dWF), which is obtained by convolving the HiFi-SIM input with the system PSF, to extract the discrepancy between measured and reconstructed intensities, thereby preserving the intensity linearity and signal-to-noise ratio (SNR) (**Supplementary Fig. 5; Supplementary Tables 2–4**). Features from both branches are fused through the SR intensity difference extraction (SIDE) module, which comprises alternating residual and channel attention blocks and incorporates spatial-attention and frequency-domain channel-attention mechanisms to prioritize regions requiring correction while improving training stability and convergence quality (**Fig. 1b; Supplementary Figs. 3b-d, 6 and 7; Supplementary Table 5**). Finally, an identity skip connection integrates the input HiFi-SIM image with the learned intensity correction map to produce the final quantitative output (denoted qHiFi-SIM). By decoupling intensity correction learning from structural prior, this architecture allows the network to leverage the HiFi-SIM structural reference without relearning structural features across imaging systems or sample types, thereby reducing sensitivity to system parameters and specimen types. Ablation studies confirmed that the skip connection contributes to training stability and noise robustness (**Supplementary Table 6**). Systematic transferability experiments are discussed in the next section.

qHiFi-SIM is trained in a self-supervised manner by minimizing a composite loss function

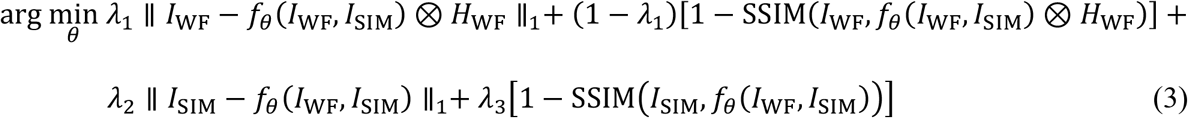

where ∥·∥_1_ denotes the L1-norm for distance calculation; *I*_WF_ is the experimental WF reference, *I*_SIM_ is the HiFi-SIM reconstruction, and *f*_*θ*_ represents the network-predicted SR reconstruction. The first two terms constitute the intensity fidelity loss, computed at the WF PSF scale between the synthetic WF (denoted qWF), generated by convolving the network output with the system PSF, and the experimental WF reference (**Fig. 1a,b**): the L1 term imposes the WF constraint of Eq. (**2**), and the SSIM term enforces perceptual consistency. The last two terms form the structural fidelity loss, preserving the SR structural details of the HiFi-SIM reconstruction. This composite loss is a differentiable realization of the two priors in Eqs. (**1**) and (**2**) and a practical relaxation of the idealized strictly convex setting, with the structural fidelity terms instantiating the regularizer in Eq. (**2**). The weighting factors λ_1_, λ_2_, and λ_3_ balance the intensity and structural terms and were optimized on the validation set (**Supplementary Note 3; Supplementary Fig. 8; Supplementary Table 7**). Because the WF image serves as the physical intensity reference, this self-supervised training requires no GT SR intensity labels. Training was conducted on high-SNR F-actin samples (labeled as RawSIMData_gt) from the BioSR dataset (**Methods**). See the **Methods** section for details on the principle and implementation of qHiFi-SIM.

### Simulation and experimental validation of qHiFi-SIM

To enable quantitative evaluation independent from the WF reference used for training, we performed a simulation with known GT distribution. Using the F-actin SR image from the test dataset reconstructed by HiFi-SIM as the GT, we simulated 2D-SIM imaging under matched acquisition parameters (emission 525 nm, numerical aperture 1.41, pixel size 62.6 nm) and compared reconstructions from HiFi-SIM and qHiFi-SIM (**Fig. 1c**). The HiFi-SIM reconstruction was normalized using the PSF-convolution least-squares method (**Supplementary Note 2**). Although both algorithms achieved comparable SR structural resolution, qHiFi-SIM exhibited substantially higher visual similarity to the GT (**Fig. 1c,e,f**), reducing MSE by ∼23.6-fold and NRMSE by ∼4.9-fold (**Fig. 1d**, left). Consistently, when the reconstructed SR images were degraded with the system PSF, qWF (SSIM = 0.9947, PSNR = 45.17 dB) matched the experimental WF far more closely than dWF (SSIM = 0.8668, PSNR = 26.73 dB; **Fig. 1d**, middle), with error analysis confirming a ∼69.8-fold and ∼8.4-fold reduction in MSE and NRMSE, respectively (**Fig. 1d**, right).

To assess the intensity linearity, we performed region-of-interest (ROI)-based analysis at the WF PSF and SIM PSF scales using 500 randomly selected ROIs per scale (**Fig. 1g–j,m–p; Supplementary Fig. 9**). At the WF PSF scale, qWF achieved near-perfect correlation with the experimental WF (R^2^ = 0.9999; mean intensity ratio of 0.981 ± 0.002), whereas dWF exhibited lower linearity (R^2^ = 0.9279; ratio = 0.770 ± 0.045) (**Fig. 1g,h**). At the SIM PSF scale, the smallest physically reliable unit for intensity quantification in SIM images, qWF maintained excellent intensity linearity (R^2^ = 0.9997 and ratio = 0.983 ± 0.003), substantially outperforming dWF (R^2^ = 0.9130; ratio = 0.865 ± 0.028) (**Fig. 1i,j**). Similarly, qHiFi-SIM consistently outperformed HiFi-SIM against GT at both PSF scales (**Fig. 1m–p**). At the SIM PSF scale, qHiFi-SIM achieved R^2^ = 0.9925 and ratio = 0.963 ± 0.009, versus HiFi-SIM (R^2^ = 0.8402; ratio = 0.794 ± 0.007). Pixel-wise analysis also showed that qHiFi-SIM outperformed HiFi-SIM (**Fig. 1k,l,q,r**). We then applied the trained network directly to the experimental F-actin raw data used to generate the simulation GT (**Fig. 1c**). As expected, qHiFi-SIM exhibited excellent intensity linearity, consistent with the simulation results (**Supplementary Fig. 10**). Across the WF PSF, SIM PSF, and single-pixel scales, qHiFi-SIM achieved R^2^ of 0.9999, 0.9999, and 0.9988, with ratios of 0.981 ± 0.008, 0.982 ± 0.008, and 0.980 ± 0.018, substantially outperforming HiFi-SIM (R^2^: 0.9460, 0.9275, 0.9334; ratios: 0.843 ± 0.126, 0.853 ± 0.161, 0.865 ± 0.169).

Collectively, these results confirm that qHiFi-SIM achieves reliable intensity linearity at the SIM PSF scale (R^2^ > 0.992 and mean intensity ratio ≥ 0.963).

We further evaluated the transferability through simulations on structurally distinct samples. A synthetic microtubule image^42^ (**Supplementary Fig. 11**) and a natural scene image^43^ (**Supplementary Fig. 12**), the latter markedly different in morphology from the F-actin training data, were used as GT under identical imaging parameters. At both WF PSF and SIM PSF scales, qHiFi-SIM achieved R^2^ > 0.994 and mean intensity ratios ≥ 0.958 against both WF and GT, whereas HiFi-SIM yielded R^2^ values of only 0.848–0.956 with mean ratios of 0.847–0.922. These results confirm that qHiFi-SIM maintains stable quantitative performance across diverse structures without retraining. We next assessed its transferability across systematic, structural, and algorithmic variations. Validation experiments included simulation on mitochondrial cristae (**Supplementary Fig. 13**), PSF mismatch tests (**Supplementary Figs. 14** and **15**), cross-training validation using F-actin and mitochondrial cristae datasets (**Supplementary Fig. 16**), and application to diverse biological samples acquired on different SIM systems, including endoplasmic reticulum (**Supplementary Fig. 17**), synaptonemal complexes (**Supplementary Fig. 18**), and external lipid droplets (**Supplementary Fig. 19**). In addition, we imaged an Argo-SIM standard slide, including a paperclip-like pattern under varying exposure times (**Supplementary Fig. 20**) and a line-pair pattern at a single exposure (**Supplementary Fig. 21**). Performance was also evaluated under varying SNR (**Supplementary Fig. 22; Supplementary Table 8**) and varying out-of-focus background conditions (**Supplementary Fig. 23; Supplementary Table 9**). Across all tested conditions (excluding the severely artifacted extreme low-SNR case; **Supplementary Fig. 22a**), qHiFi-SIM achieved R^2^ > 0.986 and mean intensity ratios ≥ 0.945 at the SIM PSF scale, whereas HiFi-SIM yielded R^2^ values of only 0.629–0.966 with mean ratios of 0.638– 0.927, demonstrating robust transferability across systems, samples, and imaging conditions.

We also assessed whether the intensity correction compromises spatial resolution using a line-pair pattern on an Argo-SIM slide (**Supplementary Fig. 21**). Both algorithms resolved 120-nm spaced lines, with decorrelation analysis^44^ yielding nearly identical resolution estimates (∼113.3 nm for HiFi-SIM vs. ∼112.3 nm for qHiFi-SIM; **Supplementary Fig. 21a**). Across nine independent datasets, qHiFi-SIM consistently maintained resolution comparable to HiFi-SIM (**Supplementary Fig. 21i**), confirming that quantitative intensity correction preserves spatial resolution. Collectively, across all tested scenarios in this study, with the exception of strong out-of-focus backgrounds where moderate resolution degradation may occur (**Supplementary Fig. 23**), qHiFi-SIM reconstructions robustly preserve the spatial resolution of the initial input SIM image.

### qHiFi-SIM simultaneously achieves high structural fidelity and optimal intensity linearity, outperforming existing SOTA approaches

To systematically benchmark qHiFi-SIM against SOTA algorithms, including HiFi-SIM^13^, fairSIM^32^, BF-SIM^18^, and FlexSIM^19^, we performed comparative reconstructions on identical F-actin data (**Fig. 2a**). For fair comparison, OTF attenuation was disabled for HiFi-SIM, fairSIM, and FlexSIM. Under these settings, fairSIM and FlexSIM achieved R^2^ values exceeding 0.990, comparable to qHiFi-SIM (**Fig. 2d–g**), but exhibited lower mean intensity ratios than qHiFi-SIM (**Fig. 2e,g**) and pronounced sidelobe artifacts^45,46^ (**Fig. 2a,b**). FlexSIM yielded low SNR, leaving many structures unresolved that were clearly visible in HiFi-SIM and qHiFi-SIM (**Fig. 2c; Supplementary Fig. 24e,f**). Further processing of fairSIM reconstructions with TV regularization^34^, Hessian regularization^12^, or sparse deconvolution^15^ [five Richardson–Lucy (RL) iterations] failed to eliminate sidelobe artifacts, and TV regularization additionally introduced staircase artifacts^12^ (**Fig. 2b; Supplementary Fig. 24a**). BF-SIM, with its inherent background-subtraction strategy, not only failed to eliminate structural artifacts but also caused structural loss (**Fig. 2a–c**), while severely degrading intensity linearity (**Fig. 2d–g**). In contrast, qHiFi-SIM effectively retained the high structural fidelity of HiFi-SIM, consistently delivering artifact-free reconstructions (**Fig. 2a–c**).

**Fig. 2.**
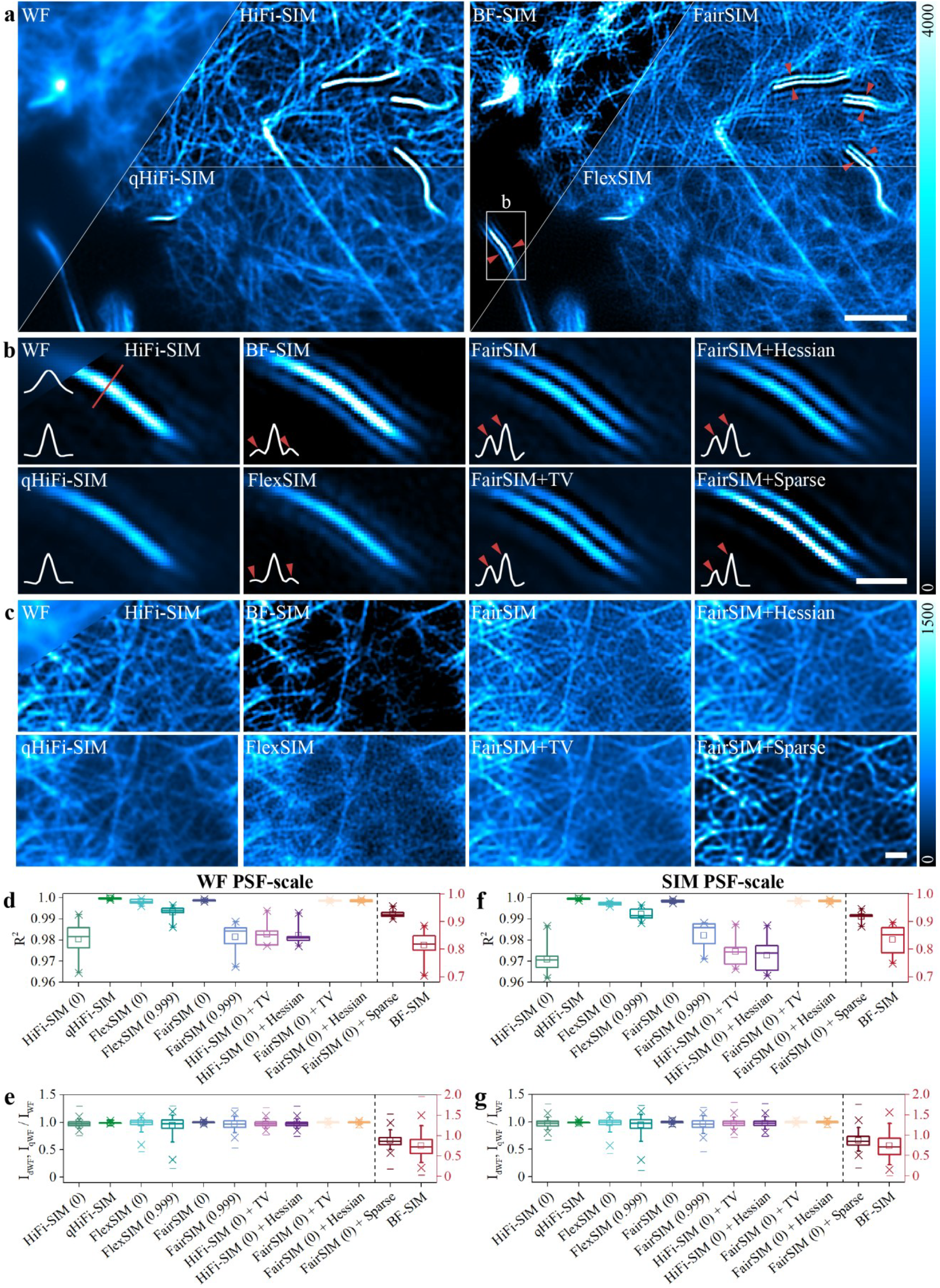
qHiFi-SIM simultaneously achieves high structural fidelity and optimal intensity linearity in SIM reconstruction. All baseline SIM reconstructions in this figure were normalized using the PSF-convolution least-squares method described in **Supplementary Note 2.a**, Comparative reconstructions of qHiFi-SIM and SOTA algorithms (HiFi-SIM, fairSIM, BF-SIM, and FlexSIM) on F-actin data. For fair comparison, OTF attenuation was disabled for HiFi-SIM, fairSIM, and FlexSIM. Red arrows highlight sidelobe artifacts in all methods except HiFi-SIM and qHiFi-SIM. **b**, Magnified views of the white-boxed region in **a**. The fairSIM reconstruction was further processed with TV regularization, Hessian regularization, and sparse deconvolution (five Richardson-Lucy iterations). Red arrows and white dashed profiles show that qHiFi-SIM inherits the structural fidelity of HiFi-SIM, whereas other methods exhibit evident artifacts. **c**, Reconstructions on an additional F-actin dataset. BF-SIM loses substantial structural information; FlexSIM yields the lowest SNR, leaving many structures unresolved; and TV regularization introduces staircase artifacts. **d–g**, Quantitative comparisons of R^2^ values (**d,f**) and intensity ratios (**e,g**) across nine test datasets at both WF and SIM PSF scales. The right red ordinate corresponds to sparse deconvolution and BF-SIM results; labels “0” and “0.999” denote OTF attenuation disabled and enabled, respectively. Intensity ratios were computed from 4,500 ROIs in total (500 randomly selected ROIs per dataset). At the WF PSF scale, FlexSIM and fairSIM with OTF attenuation disabled achieve intensity linearity comparable to qHiFi-SIM, but at the cost of structural artifacts (**a,b**); enabling OTF attenuation or applying sparse deconvolution degrades linearity, whereas BF-SIM shows the most severe deviations. At the SIM PSF scale, all methods show reduced linearity, yet qHiFi-SIM consistently achieves the highest R^2^ (>0.998) and the smallest intensity deviations (intensity ratio approximately 0.989, approaching the ideal value of 1.0). Collectively, among the methods compared, qHiFi-SIM simultaneously achieves high structural fidelity and optimal quantitative intensity linearity at both the WF PSF and SIM PSF scales. Box plots follow the same conventions as defined in the caption of **Fig. 1**. Scale bars: **a** 2 μm; **b, c** 0.5 μm.

We further quantified intensity linearity using nine independent F-actin test datasets at both WF and SIM PSF scales, with 4,500 ROIs in total (500 per dataset) at each scale, and two complementary metrics: R^2^ and mean intensity ratio (**Fig. 2d–g**). With OTF attenuation disabled, fairSIM and FlexSIM achieved R^2^ values comparable to qHiFi-SIM (**Fig. 2d,f**), but showed greater intensity ratio deviations. Specifically, qHiFi-SIM maintained mean intensity ratios of ∼0.995 at both PSF scales, whereas fairSIM dropped to ∼0.988, and FlexSIM to ∼0.914 and ∼0.909 (**Fig. 2e,g**). Enabling OTF attenuation (parameter 0.999) degraded their intensity linearity, underscoring the inherent trade-off between artifact suppression and intensity fidelity (**Fig. 2d–g; Supplementary Figs. 25** and **26**). TV and Hessian regularized denoising had relatively modest effects on intensity linearity, depending largely on the input reconstruction quality (**Fig. 2d–g; Supplementary Fig. 24**). Sparse deconvolution and BF-SIM severely compromised intensity linearity, with BF-SIM performing worst among all methods tested (R^2^ ranging from 0.7443 to 0.8829; mean intensity ratios of ∼0.617 and ∼0.608 at the WF and SIM PSF scales, respectively). At the SIM PSF scale, all methods showed reduced linearity relative to the WF PSF scale, yet qHiFi-SIM remained optimal (**Fig. 2d–g**). We further validated the intensity linearity of qHiFi-SIM using an external lipid droplet SIM dataset originally employed for BF-SIM benchmarking; qHiFi-SIM consistently outperformed BF-SIM across two intensity fidelity metrics (based on random 500 ROIs and lipid-droplet-specific regions; **Supplementary Fig. 19**). Collectively, qHiFi-SIM simultaneously achieves the best performance in structural fidelity and intensity linearity.

### qHiFi-SIM acts as a broadly applicable tool to enhance intensity linearity of other advanced algorithms

OTF attenuation and background subtraction are widely adopted strategies in many advanced SIM algorithms to suppress typical artifacts and improve optical sectioning (OS) capability, yet they inevitably impair the intensity linearity of SIM images (**Fig. 2d–f; Supplementary Figs. 25** and **26**). To demonstrate that qHiFi-SIM can eliminate the nonlinear intensity distortions caused by these operations, we applied it to SR images reconstructed by HiFi-SIM (OTF attenuation strength 0.9), fairSIM (0.999), and FlexSIM (0.999), as well as to BF-SIM. After qHiFi-SIM processing, all tested algorithms achieved R^2^ > 0.990 at both WF and SIM PSF scales (**Fig. 3a**). In addition, for FlexSIM, qHiFi-SIM not only improved intensity linearity but also moderately enhanced the SNR (**Fig. 3b**); for BF-SIM, qHiFi-SIM further enhanced the visibility of weak signal structures (**Fig. 3b**, red arrowheads). Notably, reconstructions of the same dataset by different SIM algorithms have exhibited pronounced intensity discrepancies, yet qHiFi-SIM effectively reduced these fluctuations, bringing their intensity distributions into close alignment (**Fig. 3c**).

**Fig. 3.**
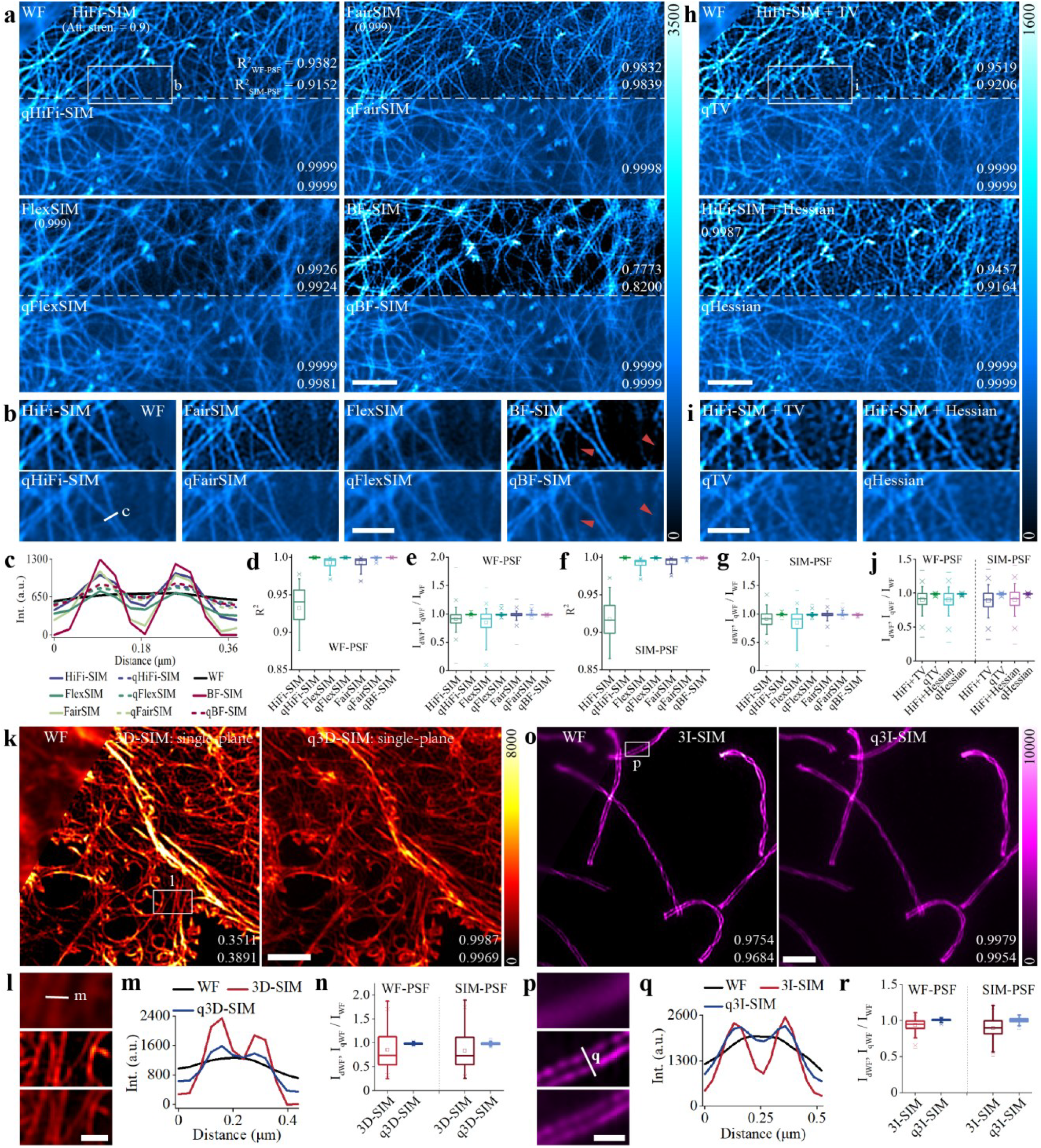
qHiFi-SIM serves as a broadly applicable tool for improving intensity fidelity across advanced SIM methods. All baseline SIM reconstructions in this figure were normalized using the PSF-convolution least-squares method described in **Supplementary Note 2.a**, Application of qHiFi-SIM to correct intensity nonlinearity from OTF attenuation (HiFi-SIM: attenuation strength 0.9; FlexSIM, fairSIM: attenuation strength 0.999) and background subtraction (BF-SIM). Top: initial SIM images; bottom: qHiFi-SIM outputs. Lower-right numbers denote R^2^ metrics between degraded (dWF, qWF) and experimental WF images at two PSF scales. **b**, Magnified views of the region indicated by the white box in **a**. Red arrowheads indicate the loss of valid weak structures in the BF-SIM reconstruction. **c**, Intensity profiles along the white line in **b. d,e**, Box plots of R^2^ (**d**) and intensity ratio (**e**) for 9 F-actin datasets before and after qHiFi-SIM processing at the WF PSF scale. BF-SIM uses the same 4,500 random ROIs (500 per dataset) as in **Fig. 2d,e**, while other algorithms adopt four attenuation strengths (HiFi-SIM: 0.8, 0.85, 0.9, 0.95; FlexSIM: 0.99, 0.995, 0.999, 0.9995; fairSIM: 0.9, 0.99, 0.995, 0.999), totaling 18,000 random ROIs per algorithm. **f,g**, Corresponding box plots of R^2^ (**f**) and intensity ratio (**g**) at the SIM PSF scale. BF-SIM uses the same 4,500 random ROIs as in **Fig. 2f,g. h**, Low-SNR F-actin data reconstructed by HiFi-SIM and subsequently denoised by TV and Hessian regularization (top row), and further processed by qHiFi-SIM (bottom row). **i**, Magnified views of the region indicated by the white box in **h. j**, Box plot of intensity ratios across three low-SNR F-actin datasets before and after qHiFi-SIM processing at WF and SIM PSF scales (1,500 random ROIs per algorithm; 500 per dataset). **k**, Comparison of open-source single-plane 3D-SIM data of actin filaments in liver sinusoidal endothelial cells (LSECs)^32^ reconstructed by HiFi-SIM and subsequently processed by qHiFi-SIM. **l**, Magnified views of the region indicated by the white box in **k. m**, Intensity profiles along the white line in **l. n**, Box plot of intensity ratios, validating the 3D performance of qHiFi-SIM. **o**, Quantitative reconstruction of synaptonemal complex SIM data from open-access 3I-SIM imaging using qHiFi-SIM. The SR image labeled “3I-SIM” was reconstructed using the open-source 3I-SIM algorithm described in ref. 49. **p**, Magnified views of the region indicated by the white box in **o. q**, Intensity profiles along the white line in **p. r**, Box plot of intensity ratios, verifying effectiveness across illumination modes. Box plots follow the same conventions as defined in the caption of **Fig. 1**. Scale bars: **a, h, k, o** 2 μm; **b, i** 1 μm; **l, p** 0.5 μm.

To quantitatively evaluate the correction efficacy of qHiFi-SIM across different algorithms, we performed a systematic analysis on nine F-actin datasets using R^2^ and mean intensity ratio metrics at both WF-and SIM-PSF scales. For HiFi-SIM, FlexSIM, and fairSIM, we systematically tested each algorithm with four distinct OTF attenuation strengths on the same dataset, yielding 18,000 random ROIs in total per algorithm. BF-SIM employed the same 4,500 random ROIs as in **Fig. 2d–g**. At both PSF scales, qHiFi-SIM significantly improved the intensity linearity of all tested algorithms across all attenuation settings and for the BF-SIM reconstruction, with post-processing R^2^ exceeding 0.997 and mean intensity ratios greater than 0.982 (**Fig. 3d–g**). For low-SNR data, TV and Hessian regularization are often applied to suppress noise artifacts in conventional SIM algorithms; however, the inherent intensity deviations of these algorithms often persist in the denoised outputs (**Fig. 2d–g; Supplementary Fig. 24**). Taking the low-SNR F-actin data (labeled RawSIMData_level_07) reconstructed by HiFi-SIM as a representative example, we applied TV and Hessian denoising, followed by further processing with qHiFi-SIM (**Fig. 3h,i**). The results demonstrate that qHiFi-SIM effectively improves the intensity linearity of the denoised reconstructions (**Fig. 3j**).

We further evaluated qHiFi-SIM on single-plane 3D-SIM data^32,47^ (**Fig. 3k–n**). Notably, while effectively correcting intensity distortions, qHiFi-SIM moderately enhanced out-of-focus signals, which reduced the OS performance of full 3D-SIM^48^ (**Supplementary Fig. 27**); thus, its direct use for full 3D-SIM is not recommended. We also tested qHiFi-SIM on the recently released SOTA 3I-SIM system, which reconstructs an SR image from only seven raw frames^49^. As expected, qHiFi-SIM effectively corrected the intensity distortions inherent to the 3I-SIM algorithm (**Fig. 3o–r**). In addition, we compared the intensity linearity of qHiFi-SIM with SOTA DL-based SR reconstruction or denoising algorithms, including DFCAN and DFGAN^22^, rDL-SIM^23^, SN2N^24^, and PRS^25^ (**Supplementary Fig. 28**). Although these methods inherit the favorable intensity linearity of the Wiener-SIM reconstructions used as the training datasets or the inputs to be denoised, subsequent qHiFi-SIM post-processing can still moderately improve their intensity linearity, achieving an R^2^ > 0.995 and mean intensity ratios ≥ 0.987 at the SIM PSF scale (**Supplementary Fig. 28c,d,g,h**). Collectively, these results establish qHiFi-SIM as a robust post-processing tool for correcting nonlinear intensity deviations in advanced SIM algorithms.

### qHiFi-SIM enables spatiotemporally accurate intensity quantification and photobleaching correction for live-cell dynamic imaging

Systematic validation on static datasets establishes the quantitative advantage of qHiFi-SIM in intensity linearity for single-frame imaging (**Figs. 1–3**). In time-lapse live-cell SR-SIM imaging, however, a fundamental challenge has long persisted: the photobleaching effect remains a key bottleneck for reliable dynamic intensity quantification^50^, beyond the intensity distortions introduced by algorithmic reconstruction. According to imaging principles, SR-SIM reconstructions and their paired WF images originate from identical raw acquisition data. As such, fluorescent samples undergo identical intrinsic photobleaching decay kinetics under both imaging modes, theoretically ensuring nearly parallel temporal intensity decay trajectories. Yet mainstream SIM reconstruction algorithms (e.g., BF-SIM, fairSIM, HiFi-SIM) inherently exhibit spatial intensity nonlinearity, which, compounded by photobleaching during time-lapse imaging, ultimately causes their temporal intensity decay trajectories to deviate nonlinearly and significantly from the WF physical baseline (**Fig. 4a,b**). This deep coupling compromises the dynamic intensity reliability of mainstream SIM algorithms and precludes the application of standard WF photobleaching correction (BC) models to time-lapse SIM imaging. For this reason, despite broad usage of SR-SIM in live-cell imaging, its practical utility remains largely confined to visualizing subcellular structural dynamics (e.g., mitochondrial cristae^12^ or outer membranes, **Fig. 4c**), rather than reliable intensity quantification over time-lapse sequences.

**Fig. 4.**
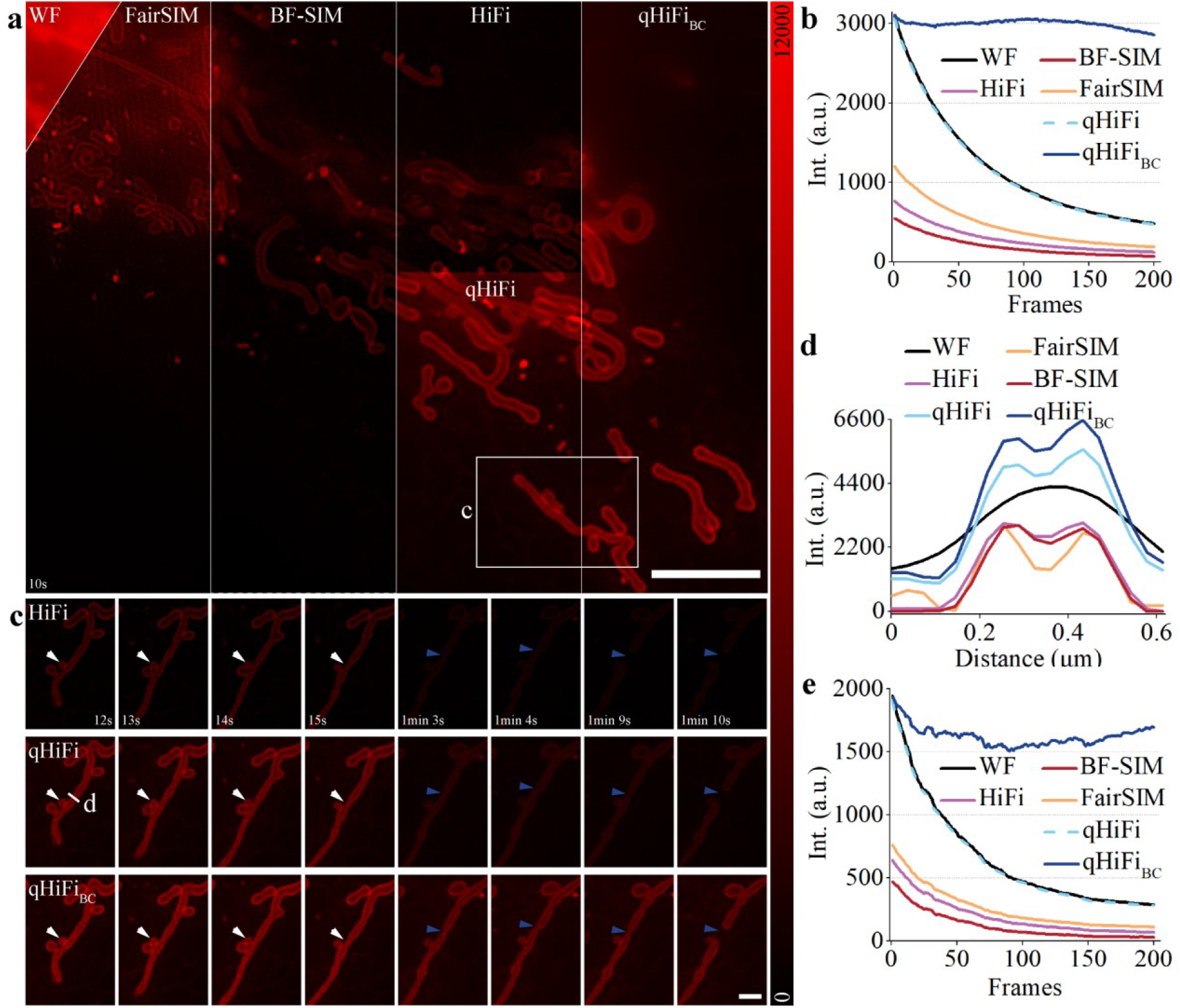
qHiFi-SIM enables quantitative visualization of mitochondrial structure and intensity dynamics decoupling algorithmic intensity bias from photobleaching. All baseline SIM reconstructions (fairSIM, BF-SIM, HiFi-SIM) are presented without normalization, as per-frame normalization will change the relative intensities across the time series. **a**, Live-cell imaging of the mitochondrial outer membrane (MOM) in U2OS cells. From left to right: WF, fairSIM, BF-SIM, HiFi-SIM, and qHiFi-SIM SR images, followed by the bleach-corrected qHiFi-SIM result (qHiFi-SIM_BC_). **b**, Mean intensity profiles across a 200-time-point series for imaging modalities indicated in **a. c**, Time-lapse images of the white-boxed region in **a**, showing the HiFi-SIM, qHiFi-SIM, and qHiFi-SIM_BC_ reconstructions. White and blue arrows mark mitochondrial fusion and fission events, respectively. **d**, Intensity profiles along the white line in **c. e**, Mean intensity profiles within the white-boxed region in **a**. Experiments were repeated independently five times, all showing similar results. Scale bars: **a** 4 μm; **c** 1 μm.

We next demonstrated the dynamic intensity quantification advantages of qHiFi-SIM in live-cell imaging. The qHiFi-SIM network was deployed on a custom-built SIM-FRET (fluorescence resonance energy transfer) platform (**Methods**), where 200-frame time-lapse imaging of the mitochondrial outer membrane (MOM) was acquired. Reconstructions with fairSIM, BF-SIM, and HiFi-SIM exhibit a non-physical decay, whereas the qHiFi-SIM output closely follows the WF physical baseline (**Figs. 4b, e**), confirming that it captures intrinsic photobleaching dynamics. This property stems from the globally unified intensity linear mapping established between the qHiFi-SIM reconstruction and the WF reference, which temporally separates the algorithm-induced spatial intensity fluctuations from the physical photobleaching effects. This capability provides a quantitative foundation for downstream intensity-based analyses of live-cell dynamic SIM data at the SIM PSF scale and beyond, such as quantitative FRET functional imaging^36,37^, and other fluorescence-intensity-dependent assays. More critically, such intensity decoupling allows, for the first time, standard WF BC models to be directly applicable to qHiFi-SIM time-lapse sequences, demonstrating a capability that, to our knowledge, has not been previously reported in conventional SIM reconstructions. As an illustration, we applied a single-exponential bleach-correction model (available in Fiji)^51^ to remove photobleaching effects, yielding corrected intensity traces (**Figs. 4b,e**). In practice, the specific BC model should be chosen based on the observed photobleaching kinetics to ensure accurate intensity recovery.

### qHiFi-SIM enables SR-FRET imaging with high-fidelity structure resolution and quantitative functional readout

Fluorescence resonance energy transfer (FRET) microscopy enables quantitative visualization of dynamic protein-protein interactions and determination of binding affinity and stoichiometry in live cells^4^. This capability stems from precise measurements of donor and acceptor fluorescence intensities, from which FRET efficiencies (*E*_D_ and *E*_A_) and the acceptor-to-donor concentration ratio (*R*_C_) are calculated to report biomolecular binding status and local microenvironments^36^. However, conventional FRET imaging is diffraction-limited, preventing the resolution of spatially adjacent molecular events at subcellular scales. Although the integration of FRET with SR-SIM has enabled molecular interaction analysis within sub-diffraction regions^36,37,52^, its quantitative reliability remains constrained by intensity inaccuracies inherent to SIM reconstruction.

To address this, we employed our custom-built SIM-FRET platform incorporating the qHiFi-SIM network for quantitative SR FRET functional imaging (**Methods**). Live-cell FRET experiments were performed using the FRET reference construct ActA-G17M targeted to the mitochondrial outer membrane (MOM). The FRET reference values for ActA-G17M (*E*_D_ ≈ 0.2 and *R*_C_ ≈ 1.0) were previously reported in our earlier study^37^. Independent FLIM-FRET measurements of G17M further supported the corresponding FRET efficiency level (**Supplementary Note 5; Supplementary Fig. 30**). All spectral-crosstalk and *G*/*k* calibration parameters were kept fixed for WF-FRET and SIM-FRET calculations (**Supplementary Note 4; Supplementary Fig. 29**); inter-channel registration accuracy (**Supplementary Note 6; Supplementary Fig. 31**) and the effect of channel-wise qHiFi-SIM processing (**Supplementary Note 7; Supplementary Fig. 32**) were evaluated separately. To assess the influence of different algorithms on inter-channel intensity distributions, we quantified the mean intensity ratios of the SIM reconstructions to the corresponding WF images (**Fig. 5d**). BF-SIM and HiFi-SIM exhibited channel-dependent intensity deviations across the donor-donor (DD), donor-acceptor (DA), and acceptor-acceptor (AA) channels, yielding ratios of 0.219, 0.103, and 0.124 for BF-SIM and 0.413, 0.475, and 0.484 for HiFi-SIM, respectively (**Fig. 5a–c**). While fairSIM maintained nearly consistent intensity ratios across the channels (DD: 0.567, DA: 0.568, AA: 0.568), its mean intensities were globally reduced relative to WF measurements. In contrast, qHiFi-SIM not only maintained nearly consistent channel ratios but also exhibited mean intensity distributions closely matching those of WF (DD: 0.993, DA: 0.997, AA: 0.993), ensuring reliable FRET quantification.

**Fig. 5.**
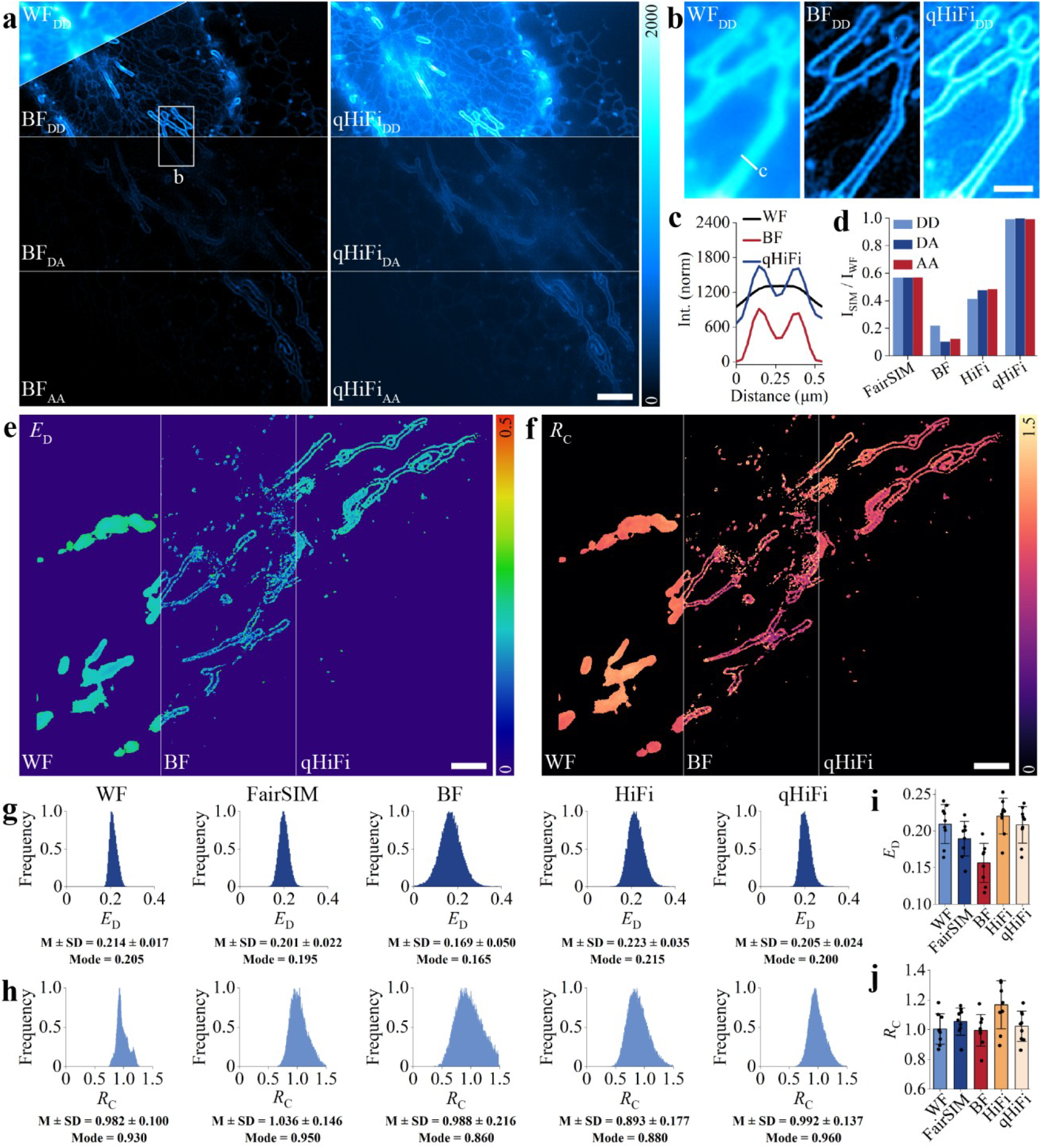
Quantitative super-resolution FRET imaging in live cells enabled by qHiFi-SIM. Baseline SIM reconstructions (fairSIM, BF-SIM, HiFi-SIM) are presented without normalization to avoid altering the relative intensities among the three FRET channels. **a**, DD, DA, and AA channel images of the mitochondrial outer membrane (MOM) reconstructed using BF-SIM and qHiFi-SIM. **b**, Magnified views of the boxed region in **a. c**, Intensity profiles along the white line in **b. d**, Mean intensity ratios of SIM reconstructions relative to the corresponding WF images for fairSIM (OTF attenuation disabled), BF-SIM, HiFi-SIM (OTF attenuation strength of 0.9), and qHiFi-SIM. **e,f**, Distributions of *E*_D_ (**e**) and *R*_C_ (**f**) obtained from WF, BF-SIM, and qHiFi-SIM images. **g,h**, Histograms of *E*_D_ (**g**) and *R*_C_ (**h**) for WF and the four SIM algorithms. Relative deviations from the nominal reference values (*E*_D_=0.20, *R*_C_=1.0) for *E*_D_ and *R*_C_, respectively, were 15.5% and 1.2% for BF-SIM, 11.5% and 10.7% for HiFi-SIM, 0.5% and 3.6% for fairSIM, and 2.5% and 0.8% for qHiFi-SIM. **i,j**, Statistical quantification of *E*_D_ (**i**) and *R*_C_ (**j**) distributions across 9 independent FoVs for WF (*E*_D_: 0.209 ± 0.025; *R*_C_: 1.004 ± 0.096) and the four SIM algorithms (fairSIM: *E*_D_: 0.190 ± 0.022; *R*_C_: 1.054 ± 0.086; BF-SIM: *E*_D_: 0.157 ± 0.025; *R*_C_: 0.996 ± 0.100; HiFi-SIM: *E*_D_: 0.220 ± 0.023; *R*_C_: 1.167 ± 0.152; qHiFi-SIM: *E*_D_: 0.208 ± 0.023; *R*_C_: 1.023 ± 0.096). Data are presented as mean ± standard deviation (SD). Scale bars: **a, e, f** 3 μm; **b** 1 μm.

We next evaluated the impact of these intensity changes on the FRET metrics. Compared with diffraction-limited WF imaging, qHiFi-SIM resolved finer structures while maintaining spatially continuous *E*_D_ and *R*_C_ distributions (**Fig. 5e,f**). Quantitative analysis in a single FoV (**Fig. 5g,h**) and across 9 independent FoVs (**Fig. 5i,j**) showed that, for *E*_D_, the relative deviations from the nominal reference value of 0.20 were 15.5% and 21.5% (for the single FoV and the 9-FoV average, respectively) for BF-SIM, 11.5% and 10.0% for HiFi-SIM, 0.5% and 5.0% for fairSIM, and 2.5% and 4.0% for qHiFi-SIM. Similarly, for *R*_C_, the relative deviations from the nominal reference value of 1.0 were 1.2% and 0.4% for BF-SIM, 10.7% and 16.7% for HiFi-SIM, 3.6% and 5.4% for fairSIM, and 0.8% and 2.3% for qHiFi-SIM. Both fairSIM and qHiFi-SIM yielded *E*_D_ values close to the reference; however, fairSIM is prone to structural artifacts (**Fig. 2a,b; Supplementary Fig. 24a**). In contrast, BF-SIM showed small deviations in *R*_C_, but large *E*_D_ deviations and a risk of losing weak structural signals (**Fig. 2c** and **Fig. 5e,f**). Collectively, qHiFi-SIM maintained both high quantitative accuracy in *E*_D_ and *R*_C_, and high structural fidelity, representing a robust tool for quantitative SR structural and functional imaging in live cells.

### High-sensitivity mapping of distinct FRET efficiencies in the cytosol and MOM via qHiFi-SIM

To evaluate the quantitative accuracy and detection sensitivity of qHiFi-SIM in heterogeneous cellular environments, we co-expressed two FRET reference constructs in live U2OS cells: mitochondria-targeted ActA-G17M (reference values, *E*_D_ ≈ 0.2 and *R*_C_ ≈ 1.0) and cytosolic G32M (reference values, *E*_D_ ≈ 0.17 and *R*_C_ ≈ 1.0) (**Fig. 6a, left panel; Supplementary Fig. 33**). Independent FLIM-FRET measurements of G17M and G32M further supported the corresponding FRET efficiency levels (**Supplementary Note 5; Supplementary Fig. 30**). FRET analysis demonstrated that qHiFi-SIM retained the structural SR advantage while yielding *E*_D_, *E*_A_, and *R*_C_ value distributions that closely aligned with WF imaging results (**Fig. 6a–d; Supplementary Fig. 34**). Quantitatively, qHiFi-SIM achieved an *E*_D_ error of only 0.001 relative to WF imaging, whereas the error for HiFi-SIM was 0.034 (**Fig. 6b**). Histograms of *E*_D_ further revealed that qHiFi-SIM resolved two distinct FRET efficiency peaks corresponding to the cytosol and MOM. In contrast, HiFi-SIM failed to clearly separate these bimodal populations due to intensity distortion and noise, and diffraction-limited WF imaging tended to merge the two peaks into a single broad peak. Interval-wise error analysis (**Methods**) confirmed that errors for qHiFi-SIM remained tightly centered around zero, in sharp contrast to the substantial fluctuations observed with HiFi-SIM (**Fig. 6f**). Statistical evaluation across nine FoVs further verified that qHiFi-SIM yielded FRET metrics consistent with the nominal reference values while preserving the SR structural advantage (**Fig. 6g–i**). In summary, these results demonstrate that qHiFi-SIM enables reliable high-sensitivity FRET readout in live cells while preserving SR structural fidelity, providing a robust tool for quantitative SR functional imaging.

**Fig. 6.**
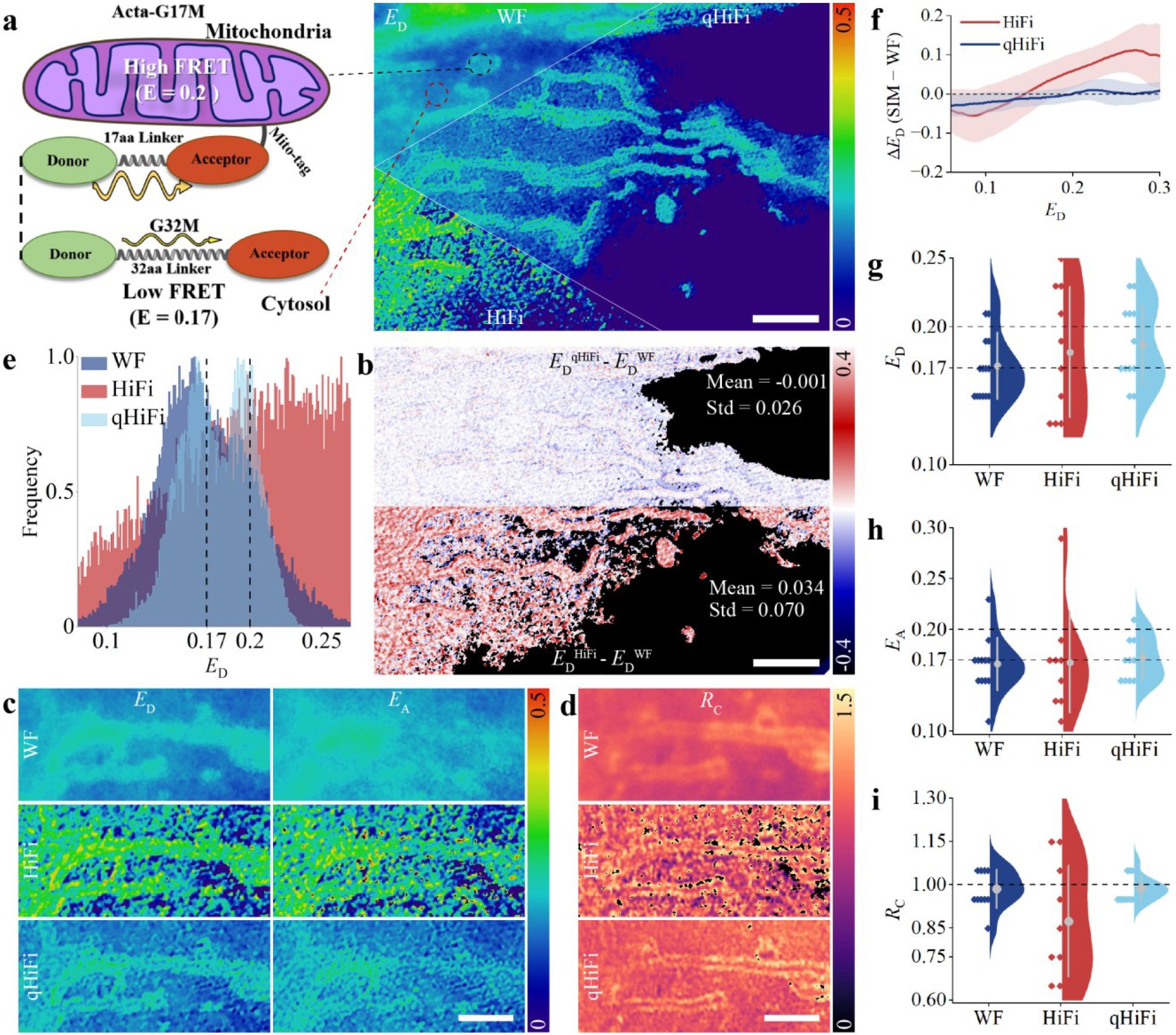
Highly sensitive detection of distinct FRET efficiencies in the cytosol and mitochondrial outer membrane via qHiFi-SIM. All HiFi-SIM reconstructions were presented without normalization to avoid altering the relative intensities among the three FRET channels. **a**, Live U2OS cells co-expressing two calibration plasmids, mitochondria-targeted ActA-G17M (*E*_D_ ≈ 0.2; *R*_C_ ≈ 1.0) and cytosolic G32M (*E*_D_ ≈ 0.17; *R*_C_ ≈ 1.0), were imaged using the SIM-FRET platform (**Supplementary Fig. 33; Methods**). The reference *E*_D_ values were independently validated by FLIM-FRET (**Supplementary Fig. 30**). Representative *E*_D_ distributions were obtained from WF, HiFi-SIM, and qHiFi-SIM images. **b**, Difference maps comparing the *E*_D_ distributions obtained from SIM reconstructions with those from WF imaging. **c,d**, Pseudo-color maps of *E*_D_, *E*_A_, and *R*_C_ in the local area. **e**, Histograms of *E*_D_ distribution in **a**, showing that qHiFi-SIM resolves two distinct peaks corresponding to the cytosol and MOM. **f**, Difference in *E*_D_ between SIM reconstructions and WF imaging over the full FoV (**Supplementary Fig. 34**). Solid lines represent the bias, and shaded regions represent SD. **g–i**, Statistical quantification of *E*_D_, *E*_A_, and *R*_C_ obtained from nine independent FoVs. Internal vertical lines indicate SD, and solid dots indicate the mean. Scale bars: **a, b** 4 μm; **c, d** 2 μm.

## Discussion

In this study, we introduce qHiFi-SIM, a physics-driven self-supervised learning framework that overcomes the long-standing trade-off between structural fidelity and intensity linearity in SR-SIM. Current SOTA SIM algorithms achieve high structural fidelity but often compromise intensity linearity through nonlinear processing steps, whereas supervised DL approaches are fundamentally limited by the lack of experimentally accessible GT intensity references at the SR scale. qHiFi-SIM addresses this gap by establishing a self-supervised DL regime that uses the actual WF image as a physically grounded intensity reference. The network is trained to minimize the intensity discrepancy between this reference and a synthetically degraded WF, generated by convolving the network-predicted SR output with the system PSF. By strategically fusing the high-fidelity structural details from HiFi-SIM with the quantitative intensity from the WF reference, qHiFi-SIM achieves excellent intensity linearity (R^2^ > 0.980; mean intensity ratio near 1), while simultaneously maintaining high structural fidelity and a twofold resolution gain. Moreover, qHiFi-SIM exhibits notable transferability across diverse SIM setups and sample structures, serving as a generalizable intensity-calibration module compatible with other SOTA SIM algorithms. To our knowledge, qHiFi-SIM represents the first SIM framework to simultaneously achieve high-fidelity SR structures and reliable intensity linearity. Furthermore, qHiFi-SIM extends quantitative SR imaging to live-cell applications by enabling reliable dynamic intensity quantification and photobleaching correction (**Fig. 4**), accurate FRET parameter quantification while preserving structural fidelity (**Fig. 5**), and high-sensitivity discrimination of closely spaced FRET populations (**Fig. 6**).

qHiFi-SIM presents certain limitations. First, when applied to an imaging modality with OS capability^47,53,54^ (e.g., 3D-SIM), the network moderately enhances out-of-focus signals, thereby reducing OS performance (**Supplementary Fig. 27**). This effect arises from the inherent out-of-focus signal in the WF reference. For data with strong backgrounds, retraining with application-specific data is a key step to avoid potential resolution degradation (**Supplementary Fig. 23**), as the qHiFi-SIM network trained in this study is based on low-background datasets. Future implementations could potentially address this issue by employing alternative quantitative references (e.g., confocal image), although such adaptation would generally require hardware modifications to standard SIM setups. Second, under low SNR conditions, particularly when the average photon count falls to approximately 116 or lower (**Supplementary Fig. 22**), qHiFi-SIM remains susceptible to reconstruction artifacts typical of conventional SIM. A potential solution is to first obtain a structurally faithful initial SR image via deep learning^21-28^ (**Supplementary Fig. 28**), followed by quantitative intensity reconstruction using qHiFi-SIM. Third, while qHiFi-SIM could in principle be extended to nonlinear-SIM^55,56^ or to advanced deconvolution methods such as multi-resolution analysis (MRA)^35^ or sparse deconvolution combined with RL algorithm^15,33^, direct application may yield suboptimal results due to substantial differences in imaging principles and accessible frequency ranges relative to 2D-SIM. For applications requiring deconvolution of SR-SIM images to ∼60 nm resolution^15,35^, a feasible strategy would be to use the corresponding qHiFi-SIM image as a quantitative intensity reference, learn a degradation kernel that maps the deconvolved SR image back to this reference, and then retrain the model accordingly. Fourth, while qHiFi-SIM shows considerable transferability in our tests, as a data-driven DL model we recommend that users, particularly those with DL expertise, retrain the model with application-specific data to ensure optimal reconstruction performance.

As a preferred technique for live-cell dynamic imaging, SIM offers considerable potential for integration with quantitative imaging modalities such as FCS^3^, FRET^4,36,37^, FRAP^5^, and FLIM^6^. Such multimodal integration is expected to accelerate the development of quantitative SR imaging. Previously, combining these methods with SR-SIM has been constrained by the lack of quantitative intensity reconstruction in traditional SIM, whereas qHiFi-SIM now provides a path to overcome this bottleneck. The self-supervised learning framework of qHiFi-SIM could be extended to adaptive optics (AO)-SIM^53,57^, lattice-SIM^58,59^ and lattice light-sheet microscopy^60,61^, opening routes toward high-quality, quantitative SR imaging of thick tissues.

In summary, qHiFi-SIM enables quantitative high-fidelity SR-SIM imaging while maintaining compatibility with mainstream SIM algorithms and imaging platforms. The accompanying user-friendly software package^62^ significantly lowers the accessibility barrier for researchers in computational optics and the life sciences. We anticipate that continued refinement and dissemination of qHiFi-SIM, coupled with advances in functional probes and the growing need for dynamic imaging, would collectively catalyze a paradigm shift in quantitative SR microscopy from structural visualization toward truly functional, quantitative imaging.

## Methods

### Network architecture and implementation of qHiFi-SIM

qHiFi-SIM accepts two inputs: an SR image reconstructed using conventional SIM algorithms (e.g., HiFi-SIM^13^), and an experimental WF image serving as a physical intensity reference within the diffraction limit. Let *I*_WF_ ∈ ℝ^H×W^ denote the input WF image, *I*_SIM_ ∈ ℝ^H×W^ the input SR-SIM image, and *H*_WF_ the system PSF. A dual-branch feature extraction strategy is employed to simultaneously achieve high-fidelity structural resolution and quantitative intensity correction. This design was inspired by the physical imaging model in Eq. (**1**), wherein the WF image is mathematically related to the in-focus sample signal *S*_in_ via convolution with the system PSF. Although this intensity at the SR scale cannot be directly measured experimentally, the WF image provides a physically constrained and accurate intensity profile within the diffraction limit. The network first generates a degraded WF image by convolving the SR image with the PSF, as follows

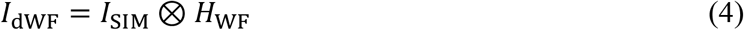

where ⊗ denotes the convolution operation implemented via a fixed convolutional layer. Then, two parallel branches are employed for feature extraction. Briefly, the WF intensity difference feature extraction branch (WIFE) processes the concatenation of *I*_WF_ and *I*_dWF_, with the mapping function defined as

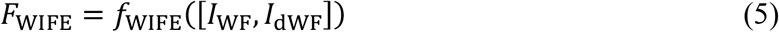

where *f*_WIFE_(*x*) = *GELU* (BN^(2)^ (Conv^(2)^ (*GELU* (BN^(1)^ (Conv^(1)^(*x*)))))), with Conv^(1)^ and Conv^(2)^ are 3×3 convolutional layers, BN^(1)^ and BN^(2)^ represent the batch normalization layers, and GELU denotes the Gaussian error linear unit activation function. The SR structural feature extraction branch (SSFE) processes the input SR image *I*_SIM_, with its mapping function defined as

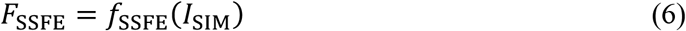

where *f*_SSFE_(*x*) = ELU(BN(Conv_SSFE_(*x*)), α = 1.0), with Conv_SSFE_ represents a 3×3 convolutional layer, BN is a batch normalization layer, and ELU denotes an exponential linear unit activation function with parameter α.

The features from both branches are concatenated channel-wise to form the combined feature maps, formulated as

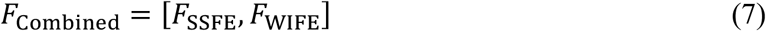

where [·] denotes the channel-wise concatenation. These combined features are processed by the SR intensity difference extraction module (SIDE), which consists of six alternatively arranged building blocks. Even-numbered blocks integrate Fourier channel attention blocks (FCABs) with residual blocks (RBs), whereas odd-numbered blocks contain solely RBs. This architectural design implements a progressive feature mapping strategy that preserves spatial information by avoiding aggressive resampling operations. The RBs facilitate stable gradient flow through identity skip connections, while the FCABs compute channel-wise attention weights using both magnitude and phase information from the Fast Fourier Transform (FFT). The spatial attention block (SAB) further refines the output by generating spatially-weighted attention maps through simultaneous channel-wise average and max pooling operations processed by a 7×7 convolutional layer, effectively highlighting regions requiring intensity correction and enhancing the precision of local intensity restoration. The RB for an input *x* is defined as

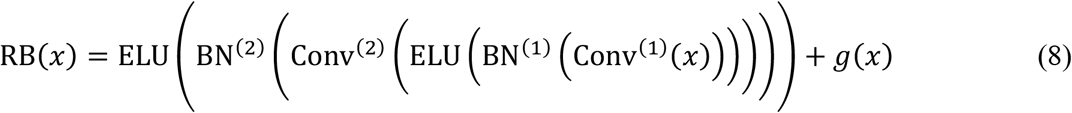

where Conv^(1)^ and Conv^(2)^ represent the 3×3 convolutional layers with batch normalization, ELU employs the parameter α = 0.45, and *g*(*x*) denotes a shortcut connection implemented via a 1×1 convolution with batch normalization when channel dimensionality requires adjustment. The operations performed by FCAB can be formulated as

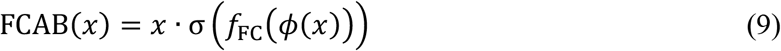

where *ϕ*(*x*) = Pool_global_([|FFT(*x*)|, ∠FFT(*x*)]) computes global average pooling across both magnitude and phase components of the FFT of input *x, f*_FC_ constitutes a sequence of fully connected layers incorporating GELU and sigmoid activations σ represents the sigmoid function that generates channel-wise attention weights. The output from the SIDE module, denoted as *F*_SIDE_, subsequently undergoes processing through a spatial attention block (SAB), as follows

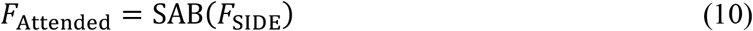

where SAB is defined as

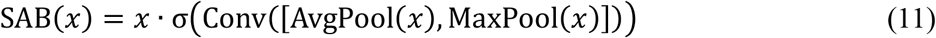

where AvgPool and MaxPool represent average and max pooling operations along the channel dimension, Conv denotes a convolutional layer with a 7×7 kernel, and σ indicates the sigmoid activation function. The SAB employs a dual-pathway architecture that integrates both average- and max-pooled features across channel dimensions to generate a discriminative attention map. This integrated approach enables the capture of widespread intensity trends through average pooling while simultaneously preserving salient local features via max pooling. The resulting attention map effectively prioritizes spatial regions exhibiting significant intensity discrepancies, thereby enhancing the precision of local intensity restoration while suppressing less informative areas.

The reconstruction module then produces the final corrected output as

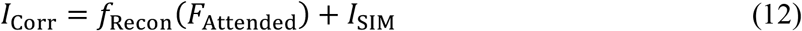

where *f*_Recon_ is defined as: *f*_Recon_(*x*) = GELU (Conv^(2)^ (GELU (Conv^(1)^(*x*)))), with Conv^(1)^ and Conv^(2)^ denoting 3×3 convolutional layers. A synthetically degraded WF image, *I*_dCorr_ = *I*_Corr_ ⊗ *H*_WF_, is also generated for loss computation. Systematic ablation studies on the SSFE branch, WIFE branch, SIDE submodules (FCAB and SAB), the identity skip connection, and the loss terms are provided in **Supplementary Figs. 4–8 and Supplementary Tables 1–7**.

### Network training and inference

The objective function of qHiFi-SIM comprises a quantitative intensity loss and a structural fidelity loss, formulated as follows

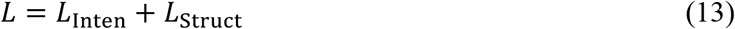

where the quantitative intensity loss, *L*_Inten_, ensures physically accurate intensities by enforcing consistency between the experimentally acquired WF image (*I*_WF_) and a synthetically degraded WF (*I*_dCorr_), defined as

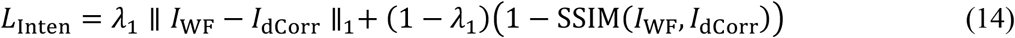

where *I*_dCorr_ is obtained by convolving the corrected output with the system PSF, yielding a self-supervised signal for intensity calibration. Here, ∥·∥_1_ denotes the L1-norm and SSIM the structural similarity index. Concurrently, the structural fidelity loss *L*_Struct_, preserves the high-resolution structure detail present in the input SR-SIM image, defined as

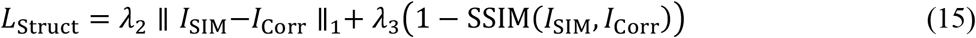

with λ_1_ = 0.5, λ_2_ = 0.1, and λ_3_ = 0.1. The weighting factors were optimized via a grid search on the validation set (**Supplementary Note 3; Supplementary Fig. 8; Supplementary Table 7**). The L1 terms primarily drive the intensity alignment and structural preservation, whereas the SSIM terms enhance perceptual quality and textural similarity.

Before being fed into the network, all input images, including the experimental WF image and the initial SR-SIM image, were normalized to the range [0, 1] to stabilize training and prevent gradient explosion. The network output (range [0, 1]) was then normalized to the maximum intensity of the experimental WF image, consistent with the PSF-convolution least-squares normalization described in **Supplementary Note 2**. All experiments were conducted using Python 3.12.0 and PyTorch 2.8.0 on an NVIDIA GeForce RTX 5090 GPU. Model training employed the Adam optimizer with a batch size of 16 and an initial learning rate of 1 × 10^-4^, and a fixed random seed of 42, over 150 epochs, requiring ∼24 h to complete. For inference, 1004 × 1004-pixel images were used, corresponding to reconstructed initial SIM images from 502 × 502-pixel raw BioSR test data. With a 31 × 31 pixels PSF and batch size 1, 50 warm-up and 100 test iterations yielded a mean latency of 37.25 ± 0.06 ms per image (min 37.20 ms, max 37.64 ms) and a peak GPU memory of 1629.7 MB. GPU inference is supported; CPU-only inference is possible but not optimized and is not recommended for routine use due to higher latency.

### PSF generation

The qHiFi-SIM network utilized the same theoretical 2D PSF model as the previously established HiFi-SIM framework^13^. During training, the 2D OTF was analytically derived from the specified imaging parameters (including numerical aperture, NA; emission wavelength, λ_em_; and effective pixel size), and the PSF was obtained as the modulus of the inverse Fourier transform of this OTF. The PSF was then convolved with both the input high-fidelity SR-SIM image and the network-generated SR image to produce corresponding degraded WF images. Owing to the robustness of qHiFi-SIM to PSF mismatch (**Supplementary Figs. 14** and **15**), a theoretical PSF generated from the imaging parameters of the target dataset is sufficient to ensure stable and high intensity linearity of qHiFi-SIM, thereby avoiding additional calibration and improving accessibility. To fully cover the main lobe and the first Airy sidelobe, the PSF was sampled up to the second Airy zero. For the F-actin training data (λ_em_ 525 nm; NA 1.41; WF pixel size 62.6 nm), the second Airy zero lies at a radius of *r*_2_ = 1.1165λ/NA ≈ 416 nm, corresponding to approximately 13.3 pixels at the SIM pixel size of 31.3 nm. Therefore, a kernel of at least 29 × 29 pixels is required. To provide additional margin, the final PSF was cropped to a central 31 × 31 pixel region. This size retains the core diffraction structure needed for accurate convolution while remaining computationally efficient. The PSF was normalized such that the sum of all kernel elements equals unity, so that convolution preserves the integrated fluorescence intensity. The convolution operation was performed with symmetric zero padding to ensure that the output retained the same spatial dimensions as the input.

### Dataset preparation and preprocessing

The experimental dataset comprised 51 groups of F-actin images from the BioSR repository^21^. From each group, we extracted the high-SNR sub-file labeled “RawSIMData_gt”. Of these, 50 groups were acquired as 502 × 502 pixel raw data, yielding 1004 × 1004 pixel SIM reconstructions; the remaining group (Cell_024; **Supplementary Table 10**) was acquired as 758 × 758 pixel raw data, yielding a 1516 × 1516 pixel SIM reconstruction, from which we cropped two 1004×1004 pixel sub-images with minimal overlap and used both as separate training samples. The final dataset comprised 52 samples (35 for training, 8 for validation, and 9 for testing), with exact sample identifiers for each split provided in **Supplementary Table 10**. SR-SIM images were reconstructed using the HiFi-SIM algorithm, while the corresponding WF images were generated by averaging the raw SIM series followed by 2× frequency-domain zero-padding, with a 4× intensity scaling to compensate for the 2D fast Fourier transform (FFT) and its inverse (iFFT) normalization convention and to restore the physical intensity scale of the experimental WF images (**Supplementary Note 1; Supplementary Fig. 1a,b**). To diversify the training data, we varied the OTF attenuation strength in HiFi-SIM to obtain multiple reconstructions with different spectral weightings. For training, 256 × 256-pixel patches were extracted via sliding-window cropping and augmented by horizontal flipping, vertical flipping and rotation, yielding approximately 40320 training patches. Similarly, the validation and test sets were expanded to 4608 and 3000 patches, respectively, via sliding-window cropping and flipping/rotation. All PSFs were normalized to unit sum for consistent physical representation.

### SIM-FRET setup

The home-built qHiFi-SIM-FRET system was developed as an upgraded version of a previously reported platform^36,37^. Donor and acceptor excitation were provided by 488 nm and 561 nm laser lines, respectively, from a multicolor laser source (L4cc, Oxxius). Both excitation channels were intensity-calibrated to a common baseline and maintained under proportional illumination mode during acquisition. The collimated beam was directed into a phase modulation unit containing a polarizing beam splitter (PBS), an achromatic half-wave plate (HWP), and a ferroelectric liquid-crystal spatial light modulator (QXGA-3DM, Forth Dimension Displays) to generate diffracted beams. These beams were focused by an achromatic lens L1 (AC254-300, Thorlabs) onto the Fourier plane, where a custom spatial filter transmitted exclusively the ±1st-order beams. A quarter-wave plate (QWP) and a customized azimuthal polarizer (VIS 038 BC3 CW01, Codixx) were employed to maintain s-polarized interference across all pattern orientations. The filtered ±1st-order beams were relayed through lenses L2 (AC254-125, Thorlabs) and L3 (AC254-200, Thorlabs) to the back focal plane of the objective lens (CFI Apochromat TIRF 60X/1.49 NA, Nikon), generating structured illumination at the sample plane. Fluorescence emitted from the sample passed through a multiband dichroic mirror (DI03-R405/488/561/635, Semrock) and two bandpass emission filters: one for the donor (ET530/30x, Chroma) and one for the acceptor (BA570-625, Olympus). Finally, donor and acceptor fluorescence were collected separately by two scientific complementary metal-oxide-semiconductor (sCMOS) cameras (Fusion BT, Hamamatsu).

### SIM-FRET calibration and validation

#### FRET reference values and FLIM validation

The FRET reference values for ActA-G17M (*E*_D_ ≈ 0.2, *R*_C_ ≈ 1.0) were previously established in our earlier study^37^. The reference values for G32M (*E*_D_ ≈ 0.17, *R*_C_ ≈ 1.0) were derived from calibrated sensitized-emission FRET (E-FRET) analysis of experimental WF images acquired under the same optical configuration (**Supplementary Note 4; Supplementary Fig. 29**). Independent TCSPC-FLIM measurements of G17M and G32M were performed using a PicoQuant LSM Upgrade Kit coupled to a Zeiss LSM 780, yielding FLIM-FRET efficiencies of 0.204 ± 0.011 and 0.170 ± 0.008, respectively (**Supplementary Note 5; Supplementary Fig. 30**).

#### Spectral crosstalk and *G*/k calibration

Donor-only and acceptor-only controls were used to determine the four spectral-crosstalk coefficients (*a, b, c*, and *d*). The *G* factor was calibrated using the modified partial-acceptor-photobleaching method (mPb-G), and the *k* factor was obtained from a 1:1 donor–acceptor tandem construct. All crosstalk and *G*/*k* parameters were experimentally determined under the same optical configuration and kept fixed for WF-FRET, HiFi-SIM-FRET, and qHiFi-SIM-FRET calculations; no parameters were re-estimated after qHiFi-SIM processing (**Supplementary Note 4; Supplementary Fig. 29**).

#### Inter-channel registration and chromatic aberration correction

Inter-channel registration accuracy was assessed using 100-nm TetraSpeck multicolor beads (T7279, Thermo Fisher Scientific) acquired under the same detection configuration. After affine registration, residual errors were 0.185 ± 0.045 pixel in the central region and 0.463 ± 0.045 pixel in the peripheral region, with all beads showing sub-pixel displacement. These measurements also assessed the effectiveness of correcting wavelength-dependent lateral misregistration across the field of view (**Supplementary Note 6; Supplementary Fig. 31**).

#### Channel-wise effect of qHiFi-SIM processing

To assess whether independent qHiFi-SIM processing of individual FRET channels alters the calibrated inter-channel intensity relationships, qHiFi-SIM was applied to DD, AA, or DA individually, or consistently to all three channels, while the same spectral-crosstalk and *G*/*k* calibration parameters were kept fixed. Single-channel substitution produced channel-dependent biases in the derived FRET efficiency, whereas consistent qHiFi-SIM processing of all three channels yielded the smallest bias and spatial dispersion relative to the WF-derived FRET reference (**Supplementary Note 7; Supplementary Fig. 32**).

### Calculation of *E*_D_ and *R*_C_ in FRET imaging

To define valid FRET analysis regions, we applied a previously established binary co-localization mask based on donor and acceptor fluorescence signals^36,37,52^. The model is defined as

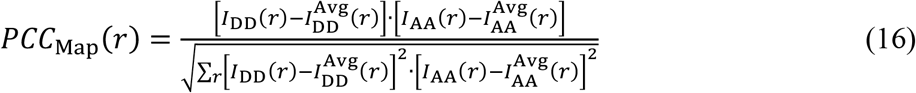

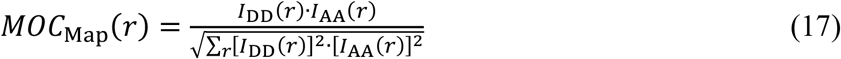

where *PCC*_Map_(*r*) and *MOC*_Map_(*r*) represent the correlation contribution maps of each pixel in donor and acceptor images to the Pearson correlation coefficient and Manders’ overlap coefficient, respectively; 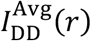 and 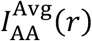 denote the mean pixel intensities of background-corrected DD and AA channel images. The hybrid map generated by multiplying *PCC*_Map_(*r*) and *MOC*_Map_(*r*) is subsequently binarized using an adaptive threshold algorithm to produce the co-localization mask *B*_mask_(*r*), which is defined as

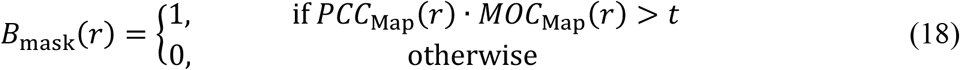

where *t* represents an empirical threshold. Following co-localization mask filtering, the donor-centric FRET efficiency (*E*_D_), the acceptor-centric FRET efficiency (*E*_A_), and the concentration ratio of total acceptor to donor (*R*_C_) were expressed as

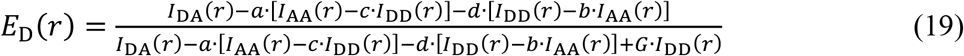

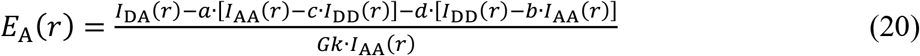

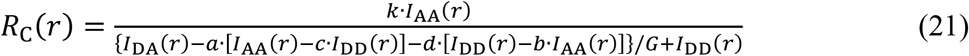

where *G* denotes the FRET sensitization-quenching transition factor, an instrument-specific calibration constant, and *k* represents the donor-to-acceptor fluorescence intensity ratio for equimolar concentrations in the absence of FRET. The calibrated values were *G* = 0.382, *k* = 2.157, *a* = 0.041, *b* = 0.001, *c* = 0.002, and *d* = 0.055.

### Data acquisition

SR-SIM imaging was conducted on several custom-built and commercial SIM platforms. 2D-SIM data for the synaptonemal complex (NA 1.49, excitation at 561 nm, emission at 605 nm, and calibrated pixel size 65 nm) and mitochondrial cristae (NA 1.49, excitation at 488 nm, emission at 525 nm, and calibrated pixel size 65 nm) were acquired with a commercial Polar-SIM system (Airy Technologies Co., Ltd.). The Argo-SIM standard slide was imaged on a DeltaVision OMX SR system (GE Healthcare) with NA 1.42, excitation at 488 nm, emission at 527 nm, and calibrated pixel size 78.6 nm. Mitochondrial outer membrane (MOM) data were collected on the custom-built SIM-FRET setup. Parameters were as follows: NA 1.49; excitation at 488 nm (DD and DA channels) or 561 nm (AA channel); emission at 525 nm (DD channel) or 600 nm (DA and AA channels); and calibrated pixel size 72.2 nm.

### Image processing

The performance of qHiFi-SIM was benchmarked against several state-of-the-art (SOTA) SIM algorithms, including HiFi-SIM^13^, fairSIM^32^, BF-SIM^18^, FlexSIM^19^, sparse deconvolution^15^, single-plane 3D-SIM procedure integrated in HiFi-SIM^13^, Open3DSIM^47^, 3I-SIM^49^, and the built-in SIM algorithm of the commercial Polar-SIM system. Total variation (TV)^34^ and Hessian^12^ regularized denoising procedures were evaluated as post-processing steps. Several deep learning–based methods were also compared, including DFCAN and DFGAN^22^, rDL-SIM^23^, SN2N^24^, and PRS^25^. Data processing was performed using Python scripts, MATLAB (v.2018a), and ImageJ (v.1.54p). All initial HiFi-SIM reconstructions fed into the qHiFi-SIM network were obtained from the MATLAB implementation of the HiFi-SIM algorithm^13^. Figures were prepared with MATLAB (v.2018a), ImageJ (v1.54p), OriginPro (v.2024b), and Adobe Illustrator (v.2020).

### Calculation of spatial resolution

Resolution was evaluated via decorrelation analysis^44^. This approach estimates the highest frequency from the local maxima of the decorrelation function, yielding an integrated metric of resolution and SNR. The decorrelation function is defined as

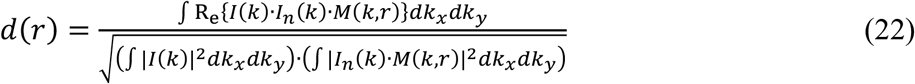

where *k* = [*k*_*x*_, *k*_*y*_] represents the spatial frequency coordinates, *I*(*k*) is the Fourier transform of the test image, *I*_*n*_(*k*) = *I*(*k*)/|*I*(*k*) is the normalized Fourier spectrum, and *M*(*k, r*) is a binary circular mask of radius *r* ∈ [0,1].

### Performance metrics

Quantitative evaluation was performed using five standard metrics: mean squared error (MSE), normalized root mean squared error (NRMSE), peak signal-to-noise ratio (PSNR), structural similarity index (SSIM), and coefficient of determination (R^2^), with Ŷ defined as the computational output and *Y* as the reference image. These metrics were computed directly from the grayscale intensity values of the test images, without additional rescaling, except where specifically indicated for particular analytical methods.

MSE is defined as

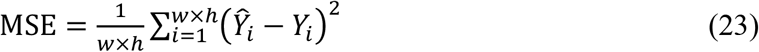

where *w* and *h* denote the image width and height in pixels.

NRMSE is defined as

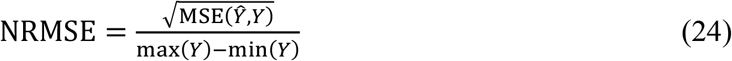

where max(*Y*) and min(*Y*) denote the maximum and minimum intensity values of the reference image *Y*, respectively.

PSNR is defined as

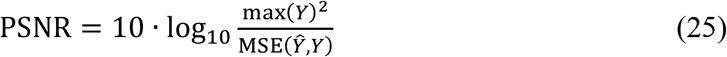

SSIM is defined as^13^

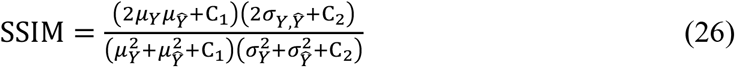

where *µ*_*Y*_ and *µ*_Ŷ_ are the mean values of *Y* and _Ŷ_, σ_*Y*_ and σ_Ŷ_ are the standard deviations, and σ_*Y*, Ŷ_ is the cross-covariance of *Y* and Ŷ, respectively. C_1_ and C_2_ are used to avoid division by a small denominator and are set as C_1_ = 0.01 and C_2_ = 0.03.

R^2^ is defined as

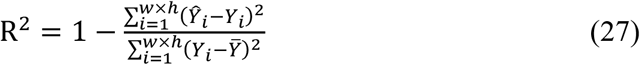

where 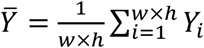 is the mean pixel intensity of the reference image *Y*.

### Calculation of the mean intensity ratio

To assess intensity linearity, we defined the mean intensity ratio as a metric for the agreement between a reconstructed (or degraded) image and its corresponding reference. For each PSF scale, 500 ROIs were randomly selected on a common coordinate grid, excluding pixels within 20 pixels of the image border to avoid edge artifacts. The ROI size was set to the WF or SIM PSF scale. The WF PSF radius at the first dark ring was *r*_WF_ = 0.61λ_em_/NA . For simplicity, the SIM PSF radius was defined assuming ideal twofold resolution improvement, i.e., *r*_SIM_ = *r*_WF_/2. In SIM pixels, for a given PSF radius *r*, the ROI size was *N* = 2⌈*r*/*p*⌉ + 1 pixels, where *p* is the SIM pixel size and ⌈·⌉ denotes the ceiling function. Thus, each ROI sampled the PSF main lobe out to approximately the first dark ring while centered on the target pixel. For every ROI, the integrated pixel intensity was calculated, and two intensity ratios were computed for the matched ROI: (i) the degraded WF (dWF or qWF) versus the experimental WF image, and (ii) the SIM reconstruction versus the corresponding GT image in simulations. The 500 ratios were used for box-plot analysis, and the mean intensity ratio was reported as mean ± SD. Parameters for the representative imaging systems and datasets used in this study are summarized in **Supplementary Table 11**.

### Synaptonemal complex preparation

The synaptonemal complex was prepared as described previously^49^. The structure comprises two lateral elements flanking a central region. Within these lateral elements, parallel protein assemblies are organized: SYCP3 proteins are spaced approximately 220 nm apart, while the C-termini of SYCP1 dimers localize to the same elements with an inter-dimer spacing of about 98 nm.

### Live-cell mitochondrial cristae labeling

Live-cell imaging of mitochondrial cristae in U2OS cells (**Supplementary Figs. 13 and 16**) was performed using PK Mito Orange (PKMO-1, PromoKine). A 250 μM stock solution was prepared by dissolving 25 nmol of the dye in 100 μL anhydrous DMSO. Immediately before staining, the stock solution was diluted 1:1000 in pre-warmed culture medium to prepare the working solution. U2OS cells grown on glass-bottom confocal dishes were gently washed once with warm medium and incubated with the PK Mito Orange working solution for 15 min at 37 °C under standard culture conditions. After incubation, the staining solution was removed, and cells were washed 1–2 times with pre-warmed medium. Live-cell fluorescence imaging was performed immediately on the commercial Polar-SIM system as described in the Data acquisition section.

### Plasmids and cloning

EGFP (Addgene #74165) and mCherry (Addgene #176016) were obtained from Addgene, while mCherry-ActA was a kind gift from David W. Andrews. The FRET reference constructs G17M and G32M were generated by inserting a linker oligonucleotides into the 5′ end of mCherry cDNA and subcloning the product into the GFP-C3 vector. To generate the mitochondrial-targeted FRET construct ActA-G17M, the ActA targeting sequence was fused to the C-terminus of G17M after removal of its stop codon. All constructs were verified by DNA sequencing and restriction enzyme digestion.

### Cell Culture and Transfection

U2OS cells (NCACC, China) were maintained in DMEM (Gibco) supplemented with 10% FBS and 1% Gentamicin-amphotericin B at 37°C with 5% CO_2_. Cells were seeded in 20-mm glass-bottom dishes and grown to 50–60% confluence before transfection. All plasmids were transfected for 24 hours using TurboFect (Fermentas) following the manufacturer’s protocol. Mycoplasma contamination was tested on the same frozen batch of U2OS and HeLa cells used in this study, and both tested negative (PCR-based detection).

## Data availability

The published BioSR dataset was used to train the qHiFi-SIM model for F-actin (https://figshare.com/articles/dataset/BioSR/13264793)^22^. The specific paired F-actin WF and reconstructed SIM images used for training the qHiFi-SIM network, together with some representative data underlying the figures (Fig. 1c–r, Figs. 2–4, and Fig. 5a), are publicly available at https://doi.org/10.6084/m9.figshare.34751451. Source data are provided with this paper.

## Code availability

The source codes of qHiFi-SIM, representative pre-trained models, as well as some example images for testing are publicly accessible via https://github.com/Yujun-DL4Micro/QHiFi-SIM.

## Author Contributions

G.W. and Y.T. conceived the research and designed the implementations. G.W., P.X., Ce.C., and T.C. supervised the project. Y.T. implemented the corresponding software and performed simulations. Z.L., X.Z., W.W., X.G., and M.L. prepared samples. G.W., Z.L., W.W., and X.G. performed imaging experiments. G.W., Y.T., Co.C., and Z.L. processed and analyzed relevant imaging data with advice from P.X., Ce.C., and T.C. G.W. composed the figures under the supervision of P.X. and T.C. G.W. wrote the manuscript with input from all authors. All authors discussed the results and commented on the paper.

## Acknowledgments

This work was supported by the National Key R&D Program of China (2023YFF1205700), the National Natural Science Foundation of China (12674364, 62665008, 62605150, 62205367 and 62141506), the Suzhou Basic Research Pilot Project (SSD2023006), Yunnan Fundamental Research Kunming Medical University Projects (No. 202601AY070001-031), and the Yunnan Revitalization Talent Support Program.

