## Supplemental Data 1 for "Physics-driven self-supervised learning for quantitative high-fidelity structured illumination microscopy"

### Table of Contents

**Supplementary Note 1:** Typical factors causing intensity deviation from true values in SIM image reconstruction

**Supplementary Note 2:** Comparison of normalization methods for SIM reconstruction intensity alignment

**Supplementary Note 3:** Roles and setting guidelines for the loss weights $\lambda_1$, $\lambda_2$, and $\lambda_3$ in qHiFi-SIM training

**Supplementary Note 4:** Calibration of spectral crosstalk and *G*/*k* factors for quantitative FRET

**Supplementary Note 5:** Independent validation of FRET efficiency by TCSPC-FLIM

**Supplementary Note 6:** Validation of inter-channel registration and chromatic aberration correction

**Supplementary Note 7:** Channel-wise effect of qHiFi-SIM processing on quantitative FRET

**Supplementary Fig. 1 |** Analysis of the impact of frequency-domain zero-padding and spatial-domain normalization on image intensity in SIM reconstruction.

**Supplementary Fig. 2 |** Comparison of normalization methods on a single raw SIM dataset.

**Supplementary Fig. 3 |** Network architecture of the qHiFi-SIM algorithm.

**Supplementary Fig. 4 |** Ablation study of the SSFE branch in qHiFi-SIM for reconstruction quality.

**Supplementary Fig. 5 |** Ablation study of the WIFE branch in qHiFi-SIM for reconstruction quality.

**Supplementary Fig. 6 |** Ablation study of the channel attention mechanism: training and validation loss curves.

**Supplementary Fig. 7 |** Ablation study of the channel attention mechanism: test set performance metrics.

**Supplementary Fig. 8 |** Optimization strategy for hyperparameters $\lambda_1$, $\lambda_2$, and $\lambda_3$ in qHiFi-SIM.

**Supplementary Fig. 9 |** Schematic illustration of region-of-interest (ROI) intensity sampling corresponding to Fig. 1.

**Supplementary Fig. 10 |** Assessing the intensity linearity of qHiFi-SIM reconstructions using experimental F-actin data from the same sample in Fig. 1.

**Supplementary Fig. 11 |** Validation of qHiFi-SIM reconstruction intensity linearity using synthetic microtubule simulation data.

**Supplementary Fig. 12 |** Validation of qHiFi-SIM reconstruction intensity linearity using natural scene simulation data.

**Supplementary Fig. 13 |** Validation of qHiFi-SIM reconstruction intensity linearity using mitochondrial cristae simulation data.

**Supplementary Fig. 14 |** Evaluating the impact of PSF mismatch on qHiFi-SIM reconstruction intensity linearity using simulated ER data.

**Supplementary Fig. 15 |** Evaluating the impact of PSF mismatch on qHiFi-SIM reconstruction intensity linearity using experimentally measured F-actin data.

**Supplementary Fig. 16 |** Evaluating the transferability of qHiFi-SIM through cross-training and validation.

**Supplementary Fig. 17 |** Assessing the intensity linearity of qHiFi-SIM reconstruction using endoplasmic reticulum structures in U2OS live cells.

**Supplementary Fig. 18 |** Assessing the intensity linearity of qHiFi-SIM reconstructions using synaptonemal complex sample.

**Supplementary Fig. 19 |** Evaluating the intensity linearity of different SIM reconstruction algorithms using an open-source lipid droplet dataset.

**Supplementary Fig. 20 |** Evaluating the intensity linearity of qHiFi-SIM reconstructions under gradient exposure times using an Argo-SIM standard slide.

**Supplementary Fig. 21 |** Evaluating the spatial resolution of qHiFi-SIM reconstructions using a resolution line-pair pattern on an Argo-SIM standard slide.

**Supplementary Fig. 22 |** Quantitative evaluation of qHiFi-SIM reconstruction intensity linearity under varying signal-to-noise ratio (SNR) levels.

**Supplementary Fig. 23 |** Quantitative evaluation of qHiFi-SIM reconstruction intensity linearity under varying out-of-focus background levels.

**Supplementary Fig. 24 |** Evaluation of intensity linearity in regularized denoising and FlexSIM reconstruction.

**Supplementary Fig. 25 |** Effects of OTF attenuation and normalization methods on the intensity linearity of fairSIM reconstructions.

**Supplementary Fig. 26 |** Effects of OTF attenuation and normalization methods on the intensity linearity of HiFi-SIM reconstructions.

**Supplementary Fig. 27 |** Evaluation of the limitations of qHiFi-SIM for intensity correction of full 3D-SIM data.

**Supplementary Fig. 28 |** Comparison of reconstruction intensity linearity between state-of-the-art deep learning–based algorithms and qHiFi-SIM.

**Supplementary Fig. 29 |** Experimental calibration of spectral crosstalk and $G/k$ factors for quantitative FRET imaging.

**Supplementary Fig. 30 |** FLIM-FRET validation using reference FRET constructs in HeLa cells.

**Supplementary Fig. 31 |** Validation of inter-channel registration accuracy.

**Supplementary Fig. 32 |** Channel-wise ablation of qHiFi-SIM processing for quantitative FRET.

**Supplementary Fig. 33 |** Large field-of-view SIM-FRET imaging data related to Fig. 6.

**Supplementary Fig. 34 |** Quantitative analysis results of large field-of-view SIM-FRET imaging data related to Fig. 6.

**Supplementary Table 1 |** Ablation study of the SSFE branch: spatial resolution comparison.

**Supplementary Table 2 |** Ablation study of the WIFE branch: SNR comparison.

**Supplementary Table 3 |** Ablation study of the WIFE branch: $R^2$ comparison.

**Supplementary Table 4 |** Ablation study of the WIFE branch: PSNR, SSIM, MSE, NRMSE comparison.

**Supplementary Table 5 |** Ablation study of the Spatial Attention mechanism: spatial resolution and SNR comparison.

**Supplementary Table 6 |** Ablation study of the identity skip connection: SNR comparison.

**Supplementary Table 7 |** Performance metrics of qHiFi-SIM under different combinations of loss weights ($\lambda_1$, $\lambda_2$, $\lambda_3$).

**Supplementary Table 8 |** Quantitative comparison of image quality metrics between the degraded WF and measured WF before and after qHiFi-SIM correction at different noise levels.

**Supplementary Table 9 |** Quantitative comparison of image quality metrics between degraded WF and measured WF before and after qHiFi-SIM correction under different background levels.

**Supplementary Table 10 |** Exact sample identifiers for the training, validation, and test splits.

**Supplementary Table 11 |** PSF-scale ROI sizes for mean intensity ratio analysis across imaging systems.

#### Supplementary Note 1: Typical factors causing intensity deviation from true values in SIM image reconstruction

##### 1.1 Standard Wiener-SIM reconstruction

Standard SIM reconstruction follows the generalized Wiener deconvolution procedure established by Heintzmann and Gustafsson[1,2] (hereafter referred to as Wiener-SIM). Briefly, starting from two-dimensional (2D) raw SIM data represented as[3]

$$I_{\theta,n}(r) \cong \left\{S_{\text{in}}(r)\cdot\left[1+m_\theta\cdot\cos\left(2\pi k_\theta r+\phi_\theta+\frac{2\pi(n-1)}{3}\right)\right]\right\}\otimes H(r)+ S_{\text{out}}(r)+N(r) \tag{1}$$

where the subscripts $\theta$ and $n$ denote the orientation and phase of the illumination pattern ($\theta = 1,2,3$, $n = 1,2,3$); $r$ represents spatial coordinates in the image; $S_{\text{in}}$ corresponds to the in-focus sample signal at the objective focal plane; $m_\theta$, $k_\theta$, and $\phi_\theta$ refer to the modulation depth, pattern wavevector, and initial phase of the illumination pattern, respectively; symbol $\otimes$ indicates the convolution operation; $H$ is the actual point spread function (PSF) of the imaging system; $S_{out}$ denotes the out-of-focus background signal, and $N$ represents the noise signal.

Applying the Fourier transform (FT) to Eq. (**1**) yields the frequency-domain representation (spectrum) of the corresponding raw SIM image, expressed as

$$\hat{I}_{\theta,n}(k) \cong \left\{\begin{matrix}\frac{m_\theta}{2}\hat{S}_{\text{in}}(k-k_\theta)\,e^{j\left[\phi_\theta+\frac{2\pi(n-1)}{3}\right]}+\hat{S}_{\text{in}}(k)\\ +\frac{m_\theta}{2}\hat{S}_{\text{in}}(k+k_\theta)e^{-\text{j}\left[\phi_\theta+\frac{2\pi(n-1)}{3}\right]}\end{matrix}\right\}\hat{H}(k)+\hat{S}_{\text{out}}(k)+\hat{N}(k) \tag{2}$$

where $\hat{S}_{in}$ denotes the spectrum of the in-focus sample; $\hat{S}_{in}(k \pm k_\theta)$ corresponds to the spectrum components containing the otherwise unresolvable high-frequency signals; $\hat{H}$ is the FT of $H$, representing the optical transfer function (OTF) of the system; and $\hat{S}_{out}$ and $\hat{N}$ represent the spectra of the out-of-focus background and the noise, respectively.

In 2D-SIM imaging, which typically involves three orientations each with three distinct phase shifts (nine raw images in total), all spectrum components in Eq. (**2**) can be separated by solving the following equation

$$\begin{bmatrix} \hat{S}_{\theta,0}(k) \\ \hat{S}_{\theta,+1}(k-k_\theta) \\ \hat{S}_{\theta,-1}(k+k_\theta) \end{bmatrix} = \begin{bmatrix} 1 & \frac{m_\theta}{2}e^{j\phi_\theta} & \frac{m_\theta}{2}e^{-j\phi_\theta} \\ 1 & \frac{m_\theta}{2}e^{j\left(\phi_\theta+\frac{2\pi}{3}\right)} & \frac{m_\theta}{2}e^{-j\left(\phi_\theta+\frac{2\pi}{3}\right)} \\ 1 & \frac{m_\theta}{2}e^{j\left(\phi_\theta+\frac{4\pi}{3}\right)} & \frac{m_\theta}{2}e^{-j\left(\phi_\theta+\frac{4\pi}{3}\right)} \end{bmatrix}^{-1} \cdot \begin{bmatrix} \hat{I}_{\theta,1}(k) \\ \hat{I}_{\theta,2}(k) \\ \hat{I}_{\theta,3}(k) \end{bmatrix} \quad (3)$$

where $\hat{S}_0$, $\hat{S}_{+1}$ and $\hat{S}_{-1}$ denote the separated 0th- and ±1st-order spectrum components, respectively.

Accurate separation of the spectrum components requires precise determination of the illumination pattern parameters, including $m_\theta$, $k_\theta$, and $\phi_\theta$, from the raw SIM images. These parameters are commonly estimated using cross-correlation-based methods[3-6]. After separation, the out-of-focus background and noise spectrum are distributed across the separated frequency components as follows

$$\begin{cases} \hat{S}_{\theta,0}(k) = \hat{S}_{\text{in}}(k)\hat{H}(k) + \left[\hat{S}_{\text{out},0}(k) + \hat{N}_0(k)\right] \\ \hat{S}_{\theta,+1}(k-k_\theta) = \hat{S}_{\text{in}}(k-k_\theta)\hat{H}(k) + \left[\hat{S}_{\text{out},+1}(k) + \hat{N}_{+1}(k)\right] \\ \hat{S}_{\theta,-1}(k+k_\theta) = \hat{S}_{\text{in}}(k+k_\theta)\hat{H}(k) + \left[\hat{S}_{\text{out},-1}(k) + \hat{N}_{-1}(k)\right] \end{cases} \quad (4)$$

where $\hat{S}_{\text{out},l}$ and $\hat{N}_l$ denote the out-of-focus background and noise, respectively, contained in the separated components of order $l$ (= −1, 0, +1). These separated components are then shifted back to their correct frequency positions. Because the residual out-of-focus background and noise in Eq. (**4**) are shifted along with the signal, the shifted components can be expressed as

$$\begin{cases} \hat{C}_{\theta,0}(k) = \hat{S}_{\text{in}}(k)\hat{H}(k) + \left[\hat{S}_{\text{out},0}(k) + \hat{N}_0(k)\right] \\ \hat{C}_{\theta,+1}(k+k_\theta) \cong \hat{S}_{\text{in}}(k)\hat{H}(k+k_\theta) + \left[\hat{S}_{\text{out},+1}(k+k_\theta) + \hat{N}_{+1}(k+k_\theta)\right] \\ \hat{C}_{\theta,-1}(k-k_\theta) \cong \hat{S}_{\text{in}}(k)\hat{H}(k-k_\theta) + \left[\hat{S}_{\text{out},-1}(k-k_\theta) + \hat{N}_{-1}(k-k_\theta)\right] \end{cases} \quad (5)$$

Finally, the super-resolution (SR) SIM image can be reconstructed by applying generalized Wiener-filter deconvolution[3-5], given by

$$I_{\mathrm{SIM}}(r) = \mathcal{F}^{-1}\left\{\frac{\sum_{\theta,L}\hat{C}_{\theta,L}(k+Lk_{\theta})\cdot\hat{H}^{*}(k+Lk_{\theta})}{\sum_{\theta,L}|\hat{H}(k+Lk_{\theta})|^{2}+w^{2}}\cdot\hat{A}(k)\right\}(r) \tag{6}$$

where $L$ indicates the order of the shifted component (for 2D-SIM data, $L = -1$, 0, +1); $\hat{A}$ is the apodization function; $w$ is the deconvolution Wiener constant, an empirically determined parameter.

Notably, under ideal conditions where the out-of-focus background and noise are absent [i.e., $\hat{S}_{\mathrm{out},l}$ and $\hat{N}_l$ in Eqs. (**4**) and (**5**) are zero] and the PSF used matches the actual imaging conditions, the SR-SIM image reconstructed via Eq. (**6**) preserves excellent quantitative intensity, as previously discussed[7]. However, out-of-focus and noise terms inevitably exist in both the raw WF and SIM spectrum [Eq. (**5**)]. Consequently, the total photon budget of the WF image is the superposition of in-focus, out-of-focus photons, and noise. Nevertheless, the WF image remains a robust intensity reference, laying a physical foundation for the subsequent development of quantitative SIM techniques. In practice, however, these unavoidable out-of-focus and noise terms ultimately render SIM images highly prone to complex artifacts[3,6-9]. As previously discussed[3,7], the out-of-focus signal in Eq. (**5**) compromises both the optical sectioning (OS) capability of the SR-SIM image and promotes the formation of honeycomb artifacts[6-9]. To mitigate this, the OTF attenuation strategy employs a Gaussian function to suppress the out-of-focus component at the center of each shifted spectrum, expressed as[3]

$$\hat{g}(k+Lk_{\theta}) = 1 - attStrength\cdot e^{-\frac{(k+Lk_{\theta})^{2}}{(0.5attWidth)^{2}}} \tag{7}$$

where $attStrength$ and $attWidth$ denote the attenuation strength and width, respectively. Both are empirically determined constants whose values depend on the quality of the raw data.

Substituting Eq. (**7**) into Eq. (**6**), the final SIM image can be expressed as

$$I_{\mathrm{SIM}}(r) = \mathcal{F}^{-1}\left\{\frac{\sum_{\theta,L}\hat{g}_{\theta,L}(k+Lk_\theta)\cdot\hat{C}_{\theta,L}(k+Lk_\theta)\cdot\hat{H}^*(k+Lk_\theta)}{\sum_{\theta,L}\hat{g}_{\theta,L}(k+Lk_\theta)\cdot|\hat{H}(k+Lk_\theta)|^2+w^2}\cdot\hat{A}(k)\right\}(r) \quad (8)$$

#### 1.2 Typical factors influencing intensity fidelity in SIM reconstruction

Based on the aforementioned principles of SIM reconstruction, several factors commonly compromise intensity fidelity in the reconstructed images. First, the attenuation function used to suppress out-of-focus background [e.g., Eq. (**7**)] concurrently modifies the spectrum weighting of the in-focus signal. This redistribution changes the relative intensities of sample structures across the field-of-view (FoV) and degrades the intensity fidelity of the final SR-SIM image (**Supplementary Figs. 25** and **26**).

A separate challenge arises from high-frequency noise, which readily generates random discontinuous artifacts such as hammerstroke patterns in SIM reconstruction[3,8]. To this end, several deconvolution algorithms, including total-variation (TV) denoising[10], Hessian regularization[6], Richardson–Lucy (RL) deconvolution[11], sparse deconvolution[12], and multi-resolution analysis (MRA) deconvolution[13], are commonly used to pre-process raw data or post-process reconstructed SIM images. Aside from TV- and Hessian-based smoothing, which only mildly affect intensity (**Supplementary Fig. 24**), the remaining methods tend to alter the relative spectrum weighting in a nonlinear manner, ultimately causing the reconstructed SIM intensity to diverge from the true sample distribution[7].

PSF mismatch presents another major source of intensity error in SIM reconstruction[3,6]. First, it biases the estimation of pattern parameters from raw data, notably the modulation depth $m_\theta$ in Eq. (**1**). Since $m_\theta$ determines the relative proportion of zero-order to higher-order spectrum components, any error directly affects the intensity map. Second, it causes the Wiener deconvolution in Eqs. (**6**) and (**8**) to employ an inaccurate OTF, introducing additional intensity distortions. As such, for applications demanding high quantitative accuracy, SIM reconstruction should be performed using a calibrated experimental PSF that matches the actual imaging conditions.

Currently, widely adopted spectrum optimization strategies in SIM algorithms, such as HiFi-SIM[3], HiFi-NL-SIM[14], and direct-SIM[15], effectively suppress common reconstruction artifacts. Yet the optimization functions they employ inevitably alter the relative weighting of frequency components within the reconstructed spectrum. As a result, although these methods can preserve high structural fidelity, they often fail to maintain reliable intensity fidelity.

In addition, practical implementation details such as zero-padding of the WF spectrum (**Supplementary Fig. 1**) and normalization of SIM algorithm outputs (**Supplementary Note 2**; **Supplementary Fig. 2**) do not affect the intensity linearity of the SR images, but can introduce global intensity deviations if the associated scaling is not properly accounted for. For example, zero-padding a 512 × 512-pixel WF image to 1024 × 1024-pixel reduces intensity values by a factor of four due to the two-dimensional fast Fourier transform (2D FFT) and inverse 2D FFT (2D iFFT) normalization convention; this scaling must be corrected to restore the physical intensity scale. For a detailed discussion of the normalization procedures used in this study, see **Supplementary Note 2**.

#### Supplementary Note 2: Comparison of normalization methods for SIM reconstruction intensity alignment

Quantitative comparison of fluorescence intensities across different SIM reconstruction methods requires proper normalization to a common reference. Throughout this study, the experimentally acquired wide-field (WF) image serves as the diffraction-limited intensity reference, as it represents the underlying fluorescence distribution convolved with the system point spread function (PSF). However, the choice of normalization method can markedly affect the intensity-based quantitative analysis of different SIM reconstructions. Here, we analyze two normalization approaches: (i) PSF-convolution least-squares normalization; and (ii) WF max-intensity normalization.

##### 2.1 Method 1: PSF-convolution least-squares normalization

The PSF-convolution least-squares normalization method seeks a scalar factor $\hat{c}$ that minimizes the squared difference between the SIM reconstruction after convolution with the system PSF and the WF reference image

$$I_{\mathrm{SIM}}^{\mathrm{norm}} = \hat{c} \cdot I_{\mathrm{SIM}},\ \hat{c} = \arg\min_{c \in \mathbb{R}_+} \| c \cdot I_{\mathrm{SIM}} \otimes H - I_{\mathrm{WF}} \|_2^2 \tag{1}$$

The optimization problem admits a closed-form solution

$$\hat{c} = \frac{\langle x, b \rangle}{\langle x, x \rangle} \tag{2}$$

where $x = \mathrm{vec}(I_{\mathrm{SIM}} \otimes H)$ and $b = \mathrm{vec}(I_{\mathrm{WF}})$ are the vectorized forms of the PSF-convolved SIM reconstruction and the WF reference, respectively. Eq. (**2**) directly encodes the optical imaging formation model: the WF image results from convolving the underlying high-resolution intensity distribution with the PSF. Solving for $\hat{c}$ yields the globally optimal scaling factor that minimizes the least-squares difference between the PSF-blurred SIM reconstruction and the actual WF measurement. This scaling accounts for the fact that SIM reconstruction intensities should be higher than WF intensities to compensate for the PSF spreading effect.

##### 2.2 Method 2: WF max-intensity normalization

The max-intensity normalization method scales the SIM reconstruction to match the maximum intensity of the WF reference image

$$I_{\text{SIM}}^{\text{norm}} = \frac{I_{\text{SIM}}}{\max(I_{\text{SIM}})} \cdot \max(I_{\text{WF}}) \tag{3}$$

where $I_{\text{SIM}}$ is the initial SIM reconstruction, and $I_{\text{WF}}$ is the WF image. This approach is computationally inexpensive and aligns the dynamic range of the SIM reconstruction with that of the WF image. However, it relies on the implicit assumption that $\max(I_{\text{SIM}})$ and $\max(I_{\text{WF}})$ are directly proportional. In practice, because the WF image is a convolved version of the underlying high-resolution distribution, the PSF spreading effect means that $\max(I_{\text{SIM}})$ should exceed $\max(I_{\text{WF}})$. Consequently, max-intensity normalization tends to underestimate the true intensity of SIM reconstructions.

##### 2.3 Application scenarios in this study

In this study, we applied the two normalization methods in different contexts depending on the experimental scenario. **Supplementary Fig. 2** provides a quantitative comparison of the two methods on representative datasets. The PSF-convolution least-squares method yields smaller errors between the degraded WF and the measured WF. Importantly, as shown in **Supplementary Figs. 25** and **26**, the two methods yield identical $R^2$ values across all OTF attenuation settings, confirming that the choice of normalization method does not affect the intensity linearity metrics ($R^2$) reported in this study.

**Scenario 1: Single-frame SIM image comparative analyses**

For all baseline reconstructions used in single-frame SIM image comparative analyses (excluding qHiFi-SIM images), including HiFi-SIM[3], fairSIM[5], BF-SIM[16], FlexSIM[17], TV-regularized[10], Hessian-regularized[6], sparse deconvolution[12], single-plane 3D-SIM[3,18], 3I-SIM[19], and deep learning-based methods, such as DFCAN and DFGAN[20], rDL-SIM[21], SN2N[22], and PRS[23], we uniformly applied the PSF-convolution least-squares normalization [Eqs. (**1**, **2**)] to ensure fair and physically consistent intensity comparison across different algorithms. In this study, all baseline reconstructions corresponding to **Figs. 1–3** and **Supplementary Figs. 10–28** were processed using this normalization.

**Scenario 2: qHiFi-SIM network output**

We normalized the qHiFi-SIM output (range [0, 1]) to the maximum intensity of the experimental WF (Eq. (**3**)). This treatment yields results consistent with the PSF-convolution least-squares normalization described above. This is described in the Methods section.

**Scenario 3: Time-lapse live-cell dynamic imaging**

For time-lapse imaging (**Fig. 4**), we applied no normalization to the outputs of the SIM algorithms compared in this study (fairSIM, BF-SIM, HiFi-SIM). Applying normalization in this context would multiply each time point by a different scalar, artificially altering the relative intensities across the time series and compromising temporal dynamics. We therefore retained the raw algorithm outputs to preserve the integrity of dynamic intensity trends. This is explicitly stated in the caption of **Fig. 4**.

**Scenario 4: SIM-FRET imaging**

For SIM-FRET imaging, we applied no normalization to the outputs of the SIM algorithms compared in this study (fairSIM, BF-SIM, and HiFi-SIM for **Fig. 5**; HiFi-SIM for **Fig. 6**). Applying normalization would multiply each FRET channel (DD, DA, AA) by a different scalar, altering the intrinsic inter-channel intensity ratios essential for FRET quantification. We therefore retained the raw algorithm outputs to preserve these inter-channel intensity ratios. This is explicitly stated in the captions of **Figs. 5** and **6**.

#### Supplementary Note 3: Roles and setting guidelines for the loss weights $\lambda_1$, $\lambda_2$, and $\lambda_3$ in qHiFi-SIM training

The qHiFi-SIM network is trained by minimizing a composite loss function defined in Eq. (**3**) of the main text and reproduced here as Eq. (**1**)

$$L = \lambda_1 \parallel I_{\mathrm{WF}} - f_\theta(I_{\mathrm{WF}}, I_{\mathrm{SIM}}) \otimes H_{\mathrm{WF}} \parallel_1$$
$$+(1-\lambda_1)[1 - \mathrm{SSIM}(I_{\mathrm{WF}}, f_\theta(I_{\mathrm{WF}}, I_{\mathrm{SIM}}) \otimes H_{\mathrm{WF}})]$$
$$+\lambda_2 \parallel I_{\mathrm{SIM}} - f_\theta(I_{\mathrm{WF}}, I_{\mathrm{SIM}}) \parallel_1$$
$$+\lambda_3\left[1 - \mathrm{SSIM}\left(I_{\mathrm{SIM}}, f_\theta(I_{\mathrm{WF}}, I_{\mathrm{SIM}})\right)\right] \quad (1)$$

where $I_{\mathrm{WF}}$ denotes the input experimental WF reference, $I_{\mathrm{SIM}}$ denotes the input HiFi-SIM reconstruction, $f_\theta(\cdot)$ denotes the network output, $\otimes$ denotes convolution, and $H_{\mathrm{WF}}$ is the system PSF. The first two terms correspond to Eq. (**14**), and the last two terms correspond to Eq. (**15**) of the main text. Equation (**1**) is the full four-term form of the training loss.

##### 3.1 Roles of the four loss terms

**Term 1 (intensity fidelity, L1):** $\lambda_1 \parallel I_{\mathrm{WF}} - f_\theta(I_{\mathrm{WF}}, I_{\mathrm{SIM}}) \otimes H_{\mathrm{WF}} \parallel_1$

It enforces photon-count conservation at the WF PSF scale between the experimental WF and the synthetic WF (qWF), generated by convolving the network output with the system PSF. Larger $\lambda_1$ drives the network output to match the WF photon budget more strictly.

**Term 2 (intensity fidelity, SSIM):** $(1-\lambda_1)[1 - \mathrm{SSIM}(I_{\mathrm{WF}}, f_\theta(I_{\mathrm{WF}}, I_{\mathrm{SIM}}) \otimes H_{\mathrm{WF}})]$

It ensures that the degraded WF image remains perceptually similar to the experimental WF, suppressing global intensity deviations that L1 alone may not capture. The weight $(1-\lambda_1)$ is complementary to $\lambda_1$, so that Terms 1 and 2 jointly constrain the photon budget and perceptual consistency at the WF PSF scale.

**Term 3 (structural fidelity, L1):** $\lambda_2 \parallel I_{\text{SIM}} - f_\theta(I_{\text{WF}}, I_{\text{SIM}}) \parallel_1$

It preserves the structural continuity of the HiFi-SIM reconstruction at the SIM PSF scale, providing spatial constraints on where photons should be allocated. Larger $\lambda_2$ keeps the network output closer to HiFi-SIM structural prior.

**Term 4 (structural fidelity, SSIM):** $\lambda_3\left[1 - \text{SSIM}\left(I_{\text{SIM}}, f_\theta(I_{\text{WF}}, I_{\text{SIM}})\right)\right]$

It preserves perceptual structural consistency with the HiFi-SIM reconstruction at the SIM PSF scale, suppressing structural artifacts. Larger $\lambda_3$ enhances visual quality; however, this SSIM term must remain non-zero, as setting $\lambda_3 = 0$ leads to severe artifacts (**Supplementary Fig. 8h**).

##### 3.2 Balancing the two competing physical priors

A central challenge in tuning the loss weights lies in balancing two competing physical priors. The WF image provides a physically accurate intensity reference but contains out-of-focus background [the term $S_{\text{out}}$ in Eq. (**1**) of the main text]. The HiFi-SIM reconstruction offers reliable structural continuity at the SIM PSF scale, yet its intensity values may deviate from the true distribution due to nonlinear operations such as spectrum optimization. Thus, the network must leverage the WF intensity reference without being misled by its out-of-focus components, while simultaneously benefiting from HiFi-SIM structural fidelity without inheriting its intensity bias.

This balance is achieved through two complementary mechanisms. The first is the WF PSF-scale intensity constraint. Terms 1 and 2 jointly constrain the network output (after convolution with the system PSF) to match the WF reference at the WF PSF scale in both photon budget and perceptual consistency. This constraint operates on the photon budget within each WF PSF region rather than on single-pixel intensities, ensuring that the total photon count matches that of the WF. Since the HiFi-SIM structural prior provides spatial constraints at the SIM PSF scale on where sharp structures are located, the network does not reconstruct the slowly varying out-of-focus background as structural features. The out-of-focus signal, as part of the physical WF measurement, is correctly accounted for at the WF PSF scale in terms of photon counts, but is not misinterpreted as structural detail. For most datasets tested in this study, the out-of-focus background is relatively low, and its impact on reconstruction quality is limited. As shown in **Supplementary Fig. 23**, notable resolution degradation is observed only under strong out-of-focus backgrounds, a limitation also noted in the Discussion of the main text. The second is the differential weighting of the two physical priors across scales. The structural terms (Terms 3 and 4), which act at the SIM PSF scale, are intentionally assigned smaller weights compared to the intensity terms (Terms 1 and 2), which act at the WF PSF scale. In the default configuration ($\lambda_1$ = 0.5, $\lambda_2$ = 0.1, $\lambda_3$ = 0.1; **Supplementary Fig. 8**), the intensity terms dominate, ensuring that HiFi-SIM serves primarily as a structural guide at the SIM PSF scale, providing spatial constraints on where photons should be allocated within each SIM PSF-scale region, rather than as an intensity reference. The intensity values are still governed by the WF photon budget through Terms 1 and 2. Consequently, the network inherits the high-resolution structural details of HiFi-SIM while substantially suppressing its intensity deviations, which might otherwise be propagated through the structural loss terms. Together, these two mechanisms enable qHiFi-SIM to simultaneously achieve high structural fidelity and reliable intensity linearity at the SIM PSF scale.

##### 3.3 Effects of imbalanced settings

As shown in **Supplementary Fig. 8** and **Supplementary Table 7**, imbalanced settings of the loss weights lead to distinct failure modes:

- **$\lambda_1$ too small with $\lambda_2$ and $\lambda_3$ too large** (e.g., $\lambda_1$ = 0.1, $\lambda_2$ = 0.9, $\lambda_3$ = 0.9)

The network overfits to the HiFi-SIM structural prior, inheriting its intensity deviations. Intensity linearity drops sharply: $R^2$ = 0.9533 at the WF

PSF scale and 0.9395 at the SIM PSF scale, with global SSIM decreasing to 0.6803.

- **$\lambda_2$ too small with $\lambda_1$ and $\lambda_3$ too large** (e.g., $\lambda_1$ = 0.8, $\lambda_2$ = 0.1, $\lambda_3$ = 0.9)

  The L1 structural preservation is weakened, leading to intensity linearity degradation: $R^2$ = 0.9575 at the WF PSF scale and 0.9454 at the SIM PSF scale, with global SSIM decreasing to 0.5745.

- **$\lambda_3$ too small with $\lambda_1$ and $\lambda_2$ too large** (e.g., $\lambda_1$ = 0.8, $\lambda_2$ = 0.9, $\lambda_3$ = 0.1)

  Insufficient SSIM weighting fails to maintain perceptual structural consistency: PSNR drops to 23.84 dB and SSIM to 0.9347, confirming that the SSIM term is essential for preserving visual quality.

- **All three $\lambda$ balanced but moderate** (e.g., $\lambda_1$ = 0.5, $\lambda_2$ = 0.5, $\lambda_3$ = 0.5)

  Performance degrades considerably when all three $\lambda$ values are moderately balanced. Compared to the optimal configuration ($\lambda_1$ = 0.5, $\lambda_2$ = 0.1, $\lambda_3$ = 0.1), $R^2$ at the WF PSF scale drops from 0.9999 to 0.9630, and $R^2$ at the SIM PSF scale drops from 0.9999 to 0.9570. PSNR decreases by 27.6 dB, SSIM decreases by 0.25, and MSE increases by over 570-fold. This indicates that $\lambda_1$ and $\lambda_2$ are competitive and cannot both be set to large values; they require careful balancing, together with a non-zero $\lambda_3$, to avoid severe performance degradation.

- **$\lambda_3$ = 0 (removing the SSIM constraint)**

  Removing the SSIM term increases PSNR to 49.92 dB but introduces severe structural artifacts (**Supplementary Fig. 8h**), confirming that SSIM is essential for artifact suppression.

- **$\lambda_2$ = $\lambda_3$ = 0 (removing all structural constraints)**

  Removing all structural constraints further increases PSNR to 51.11 dB but causes even more severe artifacts (**Supplementary Fig. 8i**), demonstrating that structural terms are indispensable for maintaining visual quality.

##### 3.4 Setting guidelines

Based on the systematic grid search performed on the validation set, we recommend the following guidelines:

**(1) Default configuration** ($\lambda_1$ = 0.5, $\lambda_2$ = 0.1, $\lambda_3$ = 0.1)

This combination achieves optimal balance among the four loss terms: $\lambda_1$ = 0.5 balances photon-count conservation and perceptual consistency at the WF PSF scale, while $\lambda_2$ = 0.1 and $\lambda_3$ = 0.1 preserve HiFi-SIM structural details while substantially suppressing its intensity deviations. Keeping $\lambda_3$ non-zero is critical for suppressing artifacts (**Supplementary Fig. 8h,i**). Under this configuration, the network achieves $R^2 > 0.999$ at both WF and SIM PSF scales, PSNR ≈ 47.09 dB, and SSIM ≈ 0.9909, with no structural artifacts.

**(2) Guidelines for parameter adjustment**

- **If higher intensity fidelity is desired:** increase $\lambda_1$ while monitoring for potential structural degradation. A larger $\lambda_1$ strengthens the WF PSF-scale photon-budget constraint, improving photon-number conservation. However, setting $\lambda_1$ too high may suppress structural variations, leading to over-smoothing or loss of fine details.

- **If better structural preservation is desired:** moderately increase $\lambda_2$ or $\lambda_3$ while monitoring intensity linearity. Increasing $\lambda_2$ or $\lambda_3$ reinforces the HiFi-SIM structural prior at the SIM PSF scale. However, excessive weighting of these structural terms may cause the network to inherit intensity deviations from HiFi-SIM, compromising intensity linearity.

- **Do not set $\lambda_3$ to 0:** the SSIM constraint is essential for suppressing structural artifacts. As shown in **Supplementary Fig. 8h,i**, removing the SSIM term ($\lambda_3$ = 0) leads to severe artifacts despite an apparent increase in PSNR.

- **Keep $\lambda_1$ and $\lambda_2/\lambda_3$ in a cooperative balance:** avoid extreme ratios where one term overwhelmingly dominates. Based on our grid search results (**Supplementary Table 7**), configurations with $\lambda_1$ = 0.5 and $\lambda_2$ = $\lambda_3$ = 0.1 achieve optimal balance.

**(3) Practical recommendations**

For most users and datasets, the default configuration ($\lambda_1$ = 0.5, $\lambda_2$ = 0.1, $\lambda_3$ = 0.1) is recommended as a robust starting point. This setting consistently delivers high intensity linearity and high structural fidelity at the SIM PSF scale across a wide range of imaging conditions and sample types without requiring manual adjustment. For users with specific requirements, such as working with low-SNR data or samples with unusual structural characteristics, application-specific fine-tuning is possible. However, this is not necessary for typical biological imaging applications.

#### Supplementary Note 4: Calibration of spectral crosstalk and *G*/*k* factors for quantitative FRET

Quantitative sensitized-emission FRET requires correction of spectral crosstalk among the DD, DA, and AA channels and calibration of instrument-dependent factors that relate sensitized acceptor emission, donor quenching, and fluorophore concentration. We therefore determined the spectral-crosstalk coefficients using donor-only and acceptor-only controls and calibrated the FRET sensitized-quenching transition factor $G$ using the modified partial-acceptor-photobleaching method (mPb-G)[24]. The $k$ factor was then obtained from a 1:1 donor–acceptor tandem reference construct using the calibrated *G* value. Representative donor-only and acceptor-only control images, the distributions of the four spectral-crosstalk coefficients, the donor-acceptor tandem construct images before and after partial acceptor photobleaching, and the resulting *G*- and *k*-factor distributions are shown in **Supplementary Fig. 29**.

##### 4.1 Spectral-crosstalk calibration

Background-subtracted fluorescence intensities were measured in the three acquisition channels: $I_{\mathrm{DD}}$ (donor excitation/donor emission), $I_{\mathrm{DA}}$ (donor excitation/acceptor emission), and $I_{\mathrm{AA}}$ (acceptor excitation/acceptor emission). The four spectral-crosstalk coefficients were determined from acceptor-only (A) and donor-only (D) controls as

$$a = \frac{I_{\mathrm{DA}}(\mathrm{A})}{I_{\mathrm{AA}}(\mathrm{A})}, b = \frac{I_{\mathrm{DD}}(\mathrm{A})}{I_{\mathrm{AA}}(\mathrm{A})}, c = \frac{I_{\mathrm{AA}}(\mathrm{D})}{I_{\mathrm{DD}}(\mathrm{D})}, d = \frac{I_{\mathrm{DA}}(\mathrm{D})}{I_{\mathrm{DD}}(\mathrm{D})} \quad (1)$$

The corresponding control images and the distributions of these four coefficients are presented in **Supplementary Fig. 29a,b**. The corrected sensitized acceptor emission attributable to FRET was calculated as

$$F_C = I_{\mathrm{DA}} - a(I_{\mathrm{AA}} - cI_{\mathrm{DD}}) - d(I_{\mathrm{DD}} - bI_{\mathrm{AA}}) \quad (2)$$

Therefore, these single-color controls explicitly quantify donor spectral bleed-through and acceptor cross-excitation contributions that contaminate the raw DA signal.

##### 4.2 *G* factor calibration using the mPb-G method

The *G* factor represents the conversion between FRET-sensitized acceptor emission and the corresponding donor fluorescence quenched by FRET. In the mPb-G calibration, the donor–acceptor tandem reference construct was first imaged before acceptor photobleaching to obtain $I_{\mathrm{DD}}$, $I_{\mathrm{AA}}$, and $I_{\mathrm{DA}}$, from which the pre-bleach corrected sensitized emission $F_{\mathrm{C}}$ was calculated. The acceptor was then partially photobleached, and the post-bleach donor and acceptor intensities, $I_{\mathrm{DD}}^{\mathrm{post}}$ and $I_{AA}^{\mathrm{post}}$, were measured. The fractional acceptor photobleaching was estimated from the decrease in the AA intensity

$$X = \frac{I_{\mathrm{AA}} - I_{\mathrm{AA}}^{\mathrm{post}}}{I_{\mathrm{AA}}} \tag{3}$$

Assuming that the AA intensity is proportional to the amount of fluorescent acceptor, *X* represents the fraction of acceptors photobleached. Accordingly, the loss of FRET-sensitized acceptor emission caused by partial acceptor photobleaching is estimated as $XF_{\mathrm{C}}$, whereas the corresponding recovery of donor fluorescence is $I_{\mathrm{DD}}^{\mathrm{post}} - I_{\mathrm{DD}}$. The *G* factor is therefore calculated as

$$G = \frac{X \cdot F_{\mathrm{C}}}{I_{\mathrm{DD}}^{\mathrm{post}} - I_{\mathrm{DD}}} \tag{4}$$

##### 4.3 *k* factor calibration and quantitative FRET calculation

After determining *G*, the quenched donor fluorescence was numerically restored by adding the FRET-dependent donor loss, $F_{\mathrm{C}}/G$, to the measured donor intensity. For the 1:1 donor–acceptor tandem construct, the *k* factor was calculated as

$$k = \frac{I_{\mathrm{DD}} + F_{\mathrm{C}}/G}{I_{\mathrm{AA}}} \tag{5}$$

The experimentally calibrated crosstalk coefficients, *G*, and *k* were subsequently used for quantitative FRET analysis. The donor-referenced FRET efficiency and acceptor-to-donor concentration ratio were calculated as

$$E_{\mathrm{D}} = \frac{F_{\mathrm{C}}}{F_{\mathrm{C}} + G \cdot I_{\mathrm{DD}}} \tag{6}$$

$$R_{\mathrm{C}} = \frac{k \cdot I_{\mathrm{AA}}}{I_{\mathrm{DD}} + F_{\mathrm{C}}/G} \tag{7}$$

All crosstalk and *G*/*k* calibration parameters were determined experimentally under the same optical configuration and were kept fixed for subsequent WF-FRET, HiFi-SIM-FRET, and qHiFi-SIM-FRET calculations. They were not re-estimated after qHiFi-SIM processing, thereby avoiding absorption of reconstruction-dependent intensity changes into the calibration parameters and allowing a direct assessment of whether qHiFi-SIM preserves the experimentally calibrated inter-channel intensity relationships. The resulting calibration constants were: $G = 0.382$, $k = 2.157$, $a = 0.041$, $b = 0.001$, $c = 0.002$, and $d = 0.055$.

**Supplementary Fig. 29c–f** shows representative images acquired before and after partial acceptor photobleaching, together with the resulting distributions of the calibrated *G*- and *k*-factors.

#### Supplementary Note 5: Independent validation of FRET efficiency by TCSPC-FLIM

Sensitized-emission FRET quantification depends on the relative fluorescence intensities measured in the DD, DA, and AA channels and therefore requires appropriate correction for spectral crosstalk and instrument-dependent calibration factors. To provide an independent quantitative reference for the FRET efficiencies used in this study, we performed time-correlated single-photon counting fluorescence lifetime imaging microscopy (TCSPC-FLIM) on the G17M and G32M reference constructs expressed in HeLa cells. FLIM measurements were acquired using a PicoQuant LSM Upgrade Kit coupled to a Zeiss LSM 780. Because FLIM estimates FRET from donor-lifetime shortening rather than from the DD/DA/AA intensity ratios, this measurement provides an orthogonal validation that is independent of donor bleed-through, acceptor direct excitation, and *G*/*k* calibration used in sensitized-emission FRET.

The donor-only reference lifetime used for the FLIM-FRET calculation was 2.710 ns. For datasets fitted with a multicomponent fluorescence-decay model, the amplitude-weighted donor lifetime was used as the effective donor lifetime in the presence of the acceptor. The corresponding expressions are

$$\tau_{\mathrm{DA}} = \tau_{\mathrm{AV,Amp}} = \frac{\sum_i A_i \tau_{\mathrm{i}}}{\sum_i A_i} \quad (1)$$

$$E_{\mathrm{FLIM}} = 1 - \frac{\tau_{\mathrm{DA}}}{\tau_{\mathrm{D}}} \quad (2)$$

where $A_i$ and $\tau_{\mathrm{i}}$ denote the fitted amplitude and lifetime of the $i$-th decay component, respectively; $\tau_{\mathrm{DA}}$ is the amplitude-weighted donor lifetime in the FRET construct; and $\tau_{\mathrm{D}} = 2.710\ ns$ is the donor-only reference lifetime.

Representative donor-intensity, fluorescence-lifetime, and FLIM-derived FRET-efficiency maps are shown in Supplementary **Fig. 30a–c**. The pixel-level distributions from the representative cells further illustrate the systematic separation between the two constructs (**Supplementary Fig. 30d**). These pixel-level distributions are provided to visualize the intracytoplasmic lifetime and FRET distributions and were not treated as independent biological replicates. At the cell level, G17M exhibited a shorter donor lifetime of 2.156 ± 0.030 ns (mean ± SD, $n$ = 10), whereas G32M exhibited a longer donor lifetime of 2.247 ± 0.020 ns ($n$ = 5). Using the same donor-only reference lifetime, these values corresponded to FLIM-FRET efficiencies of 0.204 ± 0.011 for G17M and 0.170 ± 0.008 for G32M (**Supplementary Fig. 30e**). Thus, FLIM independently resolved the higher- and lower-FRET regimes represented by G17M and G32M, respectively.

The purpose of this FLIM experiment is to provide a construct-level quantitative reference rather than a spatially resolved benchmark for the super-resolved FRET maps. The TCSPC-FLIM measurements were acquired on a diffraction-limited laser-scanning microscope and therefore do not resolve the finest subcellular structures accessible to SIM-FRET. Nevertheless, because the lifetime-based FRET efficiencies are obtained through an intensity-independent measurement principle, the observed separation between G17M and G32M provides independent support for the quantitative FRET levels used to evaluate sensitized-emission SIM-FRET in this study.

#### Supplementary Note 6: Validation of inter-channel registration and chromatic aberration correction

Quantitative SIM-FRET analysis requires accurate spatial correspondence among the DD, DA, and AA fluorescence channels because intensities from different channels are combined on a pixel-by-pixel basis. We therefore evaluated inter-channel registration accuracy using 100-nm multicolor TetraSpeck fluorescent beads (T7279, Thermo Fisher Scientific) acquired under the same detection configuration used for the SIM-FRET measurements. Representative bead images are shown in **Supplementary Fig. 31a–c**. Without loss of generality, the DD channel was used as the reference channel, and the DA and AA channels were registered to it.

Inter-channel geometric displacement was corrected using an affine transformation. For a target-channel coordinate $(x, y)$, the registered coordinate $(x', y')$ was obtained as

$$\begin{bmatrix} x' \\ y' \\ 1 \end{bmatrix} = \begin{bmatrix} a_{11} & a_{12} & t_x \\ a_{21} & a_{22} & t_y \\ 0 & 0 & 1 \end{bmatrix} \begin{bmatrix} x \\ y \\ 1 \end{bmatrix} \tag{1}$$

where $a_{11}$, $a_{12}$, $a_{21}$, and $a_{22}$ describe the linear transformation, while $t_x$ and $t_y$ represent inter-channel translation. The affine model corrects the dominant lateral geometric differences between channels, including translation, rotation, scaling, and shear.

The center of each fluorescent bead was localized with subpixel precision in both channels. After registration, the residual displacement was calculated as

$$\begin{cases} \Delta x_i = x_{\mathrm{ref},i} - x'_{\mathrm{i}} \\ \Delta y_i = y_{\mathrm{ref},i} - y'_{\mathrm{i}} \end{cases} \tag{2}$$

where $(x_{\mathrm{ref},i}, y_{\mathrm{ref},i})$ is the center coordinate of the *i*-th bead in the reference channel, and $(x', y')$ is its registered coordinate in the target channel. The radial residual registration error was defined as

$$r_i = \sqrt{(\Delta x_i)^2 + (\Delta y_i)^2} \quad (3)$$

To assess whether wavelength-dependent distortion increased toward the field periphery, beads were further divided into central and peripheral regions of the FoV (**Supplementary Fig. 31d**). The residual displacement vectors remained clustered around the zero-displacement position after affine registration (**Supplementary Fig. 31e**). In the central region, the residual registration error was 0.185 ± 0.045 pixel (mean ± SD, $n$ = 9), with an RMS error of 0.190 pixel. In the peripheral region, the residual error increased to 0.463 ± 0.045 pixel ($n$ = 9), with a root-mean-square (RMS) error of 0.465 pixel (**Supplementary Fig. 31f**). The peripheral residual error was larger than that in the central region, indicating a modest field-dependent increase in the remaining wavelength-dependent displacement. However, all 18 analyzed beads exhibited residual errors below one pixel across the entire FoV.

Because lateral chromatic aberration in multichannel fluorescence imaging manifests as wavelength-dependent spatial displacement and field-dependent geometric distortion, the multicolor-bead measurements also provide an experimental assessment of the effectiveness of the chromatic registration correction. Although an affine transformation does not explicitly model arbitrary higher-order optical distortions, the residual displacement remained within the subpixel range throughout the analyzed field. These results demonstrate that the registration procedure provides sufficiently accurate inter-channel spatial correspondence for the quantitative SIM-FRET measurements presented in this study.

#### Supplementary Note 7: Channel-wise effect of qHiFi-SIM processing on quantitative FRET

Quantitative sensitized-emission FRET depends on the calibrated relative intensities of the DD, DA, and AA channels rather than on the fidelity of any single channel in isolation. Because qHiFi-SIM is applied independently to each reconstructed fluorescence channel, a potential concern is that channel-dependent intensity correction could alter the inter-channel intensity relationships used to calculate FRET. We therefore performed a channel-wise ablation analysis using the same FRET dataset presented in **Fig. 5** to directly assess how applying qHiFi-SIM to individual channels propagates into the final FRET result.

Five reconstruction combinations were analyzed. The original HiFi-SIM reconstruction of all three channels served as the baseline. qHiFi-SIM was then substituted into DD only, AA only, or DA only while the other two channels remained as HiFi-SIM reconstructions. Finally, qHiFi-SIM was applied consistently to all three channels:

$$\text{HiFi-SIM}\ (\mathrm{DD_H}, \mathrm{AA_H}, \mathrm{DA_H})$$

$$\text{qHiFi-SIM}_{\mathrm{DD}}\ \left(\mathrm{DD_q}, \mathrm{AA_H}, \mathrm{DA_H}\right)$$

$$\text{qHiFi-SIM}_{\mathrm{AA}}\ \left(\mathrm{DD_H}, \mathrm{AA_q}, \mathrm{DA_H}\right)$$

$$\text{qHiFi-SIM}_{\mathrm{DA}}\ \left(\mathrm{DD_H}, \mathrm{AA_H}, \mathrm{DA_q}\right)$$

$$\text{qHiFi-SIM}\ \left(\mathrm{DD_q}, \mathrm{AA_q}, \mathrm{DA_q}\right)$$

where the subscript H denotes HiFi-SIM reconstruction and q denotes qHiFi-SIM reconstruction. This design isolates the effect of channel-wise qHiFi-SIM substitution from that of consistent three-channel processing. All combinations were analyzed using exactly the same experimentally determined spectral-crosstalk and *G*/*k* calibration parameters. No calibration coefficient was re-estimated for any hybrid reconstruction. This design isolates the effect of reconstruction-dependent intensity changes and prevents such changes from being absorbed into method-specific recalibration.

For each combination, the corrected sensitized acceptor emission and donor-referenced FRET efficiency were calculated using the same quantitative FRET equations as defined in **Supplementary Note 4** [Eqs. (**2**) and (**6**)].

To evaluate reconstruction-dependent FRET deviation, the experimentally measured WF result from the same sample was used as the reference for photometric comparison. The pixel-wise FRET deviation was defined as

$$\Delta E_{\mathrm{D}}(x,y) = E_{\mathrm{D}}^{\mathrm{method}}(x,y) - E_{\mathrm{D}}^{\mathrm{WF}}(x,y) \tag{1}$$

Here, the WF result is used specifically as an experimental reconstruction reference for evaluating intensity-dependent deviations and is not treated as an independent absolute FRET ground truth. The independent quantitative validation of the FRET level is provided separately by the FLIM-FRET measurements described in **Supplementary Note 5**.

As shown in **Supplementary Fig. 32**, the original HiFi-SIM reconstruction showed an overall positive FRET bias of 0.0436 ± 0.0623 (mean ± SD of the pixel-wise $\Delta E_{\mathrm{D}}$ distribution within the analyzed FRET region). Applying qHiFi-SIM only to the DD channel shifted the FRET result toward a pronounced negative bias of −0.1552 ± 0.0379, whereas applying qHiFi-SIM only to the AA channel produced a positive bias of 0.1726 ± 0.0613. Applying qHiFi-SIM only to DA substantially reduced the mean bias to 0.0130 ± 0.0605, but the residual spatial dispersion remained comparable to that of the original HiFi-SIM result. In contrast, consistent qHiFi-SIM processing of DD, AA, and DA produced the smallest deviation from the WF-derived reference, with a mean bias of 0.0031 ± 0.0239.

The opposite error directions observed when only DD or AA was corrected demonstrate that improving the photometric fidelity of one channel alone does not necessarily improve the derived FRET efficiency. This behavior is expected because $E_D$ is a nonlinear quantity determined by the coupled DD, DA, and AA intensities after spectral-crosstalk correction. Replacing only one channel changes the balance among the calibrated intensity terms and can therefore introduce a new FRET bias even when that individual channel is itself more photometrically faithful. The much smaller bias and reduced spatial dispersion obtained when all three channels were processed by qHiFi-SIM indicate that the quantitative improvement does not arise from an arbitrary adjustment of a single FRET channel. Rather, consistent qHiFi-SIM processing restores the inter-channel intensity relationship required by the fixed FRET calibration framework. These results directly address the potential channel-dependence of qHiFi-SIM processing and support the use of coordinated three-channel qHiFi-SIM reconstruction for quantitative SIM-FRET analysis.

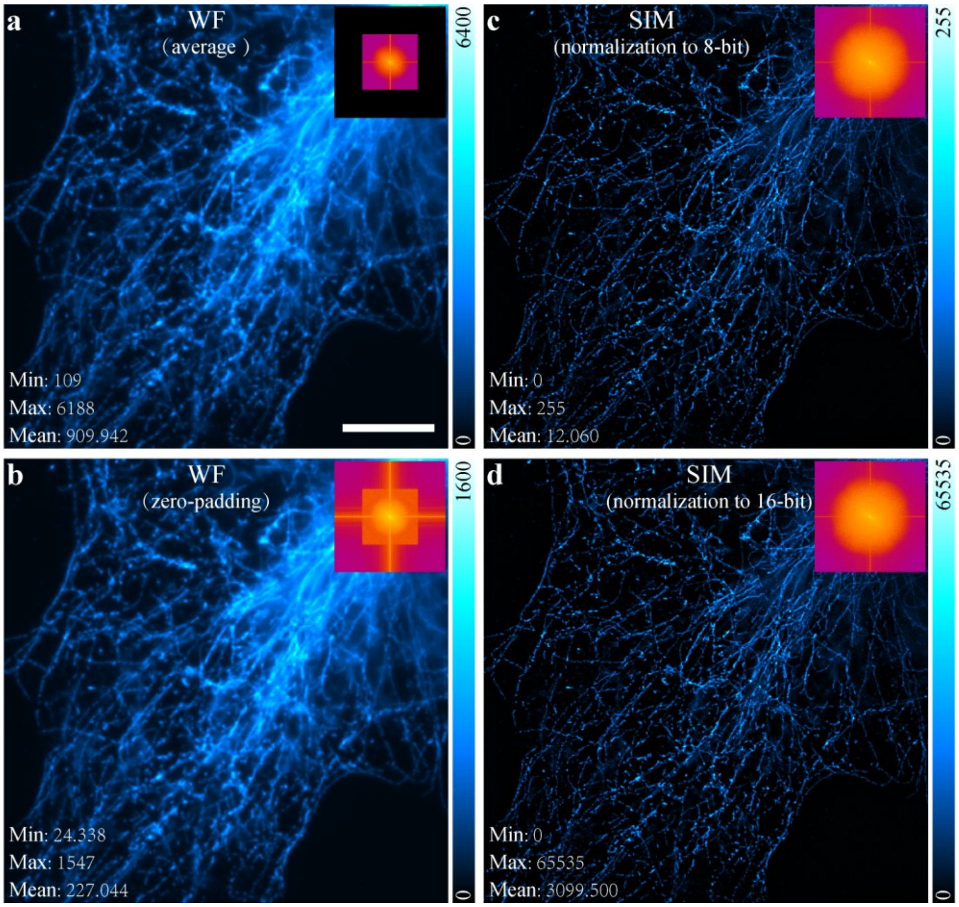


**Supplementary Fig. 1 | Analysis of the impact of frequency-domain zero-padding and spatial-domain normalization on image intensity in SIM reconstruction.** Microtubule data from fixed COS-7 cells were derived from a published, open-source 2D-SIM dataset[3]. **a**, Wide-field (WF) image obtained by averaging the nine raw SIM images; its corresponding Fourier spectrum is shown in the inset. **b**, WF image generated by applying frequency-domain zero-padding to the spectrum in **a**. The 4× reduction in intensity arises from the 2D FFT / 2D iFFT normalization convention: zero-padding from $N \times N$ to $2N \times 2N$ changes the 2D iFFT normalization factor from $1/N^2$ to $1/(2N)^2$, requiring a 4× correction to restore the physical WF intensity scale. As such, in qHiFi-SIM, intensities in **b** are multiplied by four to restore the physical WF intensity scale before network training. **c**,**d**, Reconstructed SIM images normalized to 8-bit (0–255) and 16-bit (0–65535) ranges, respectively. While normalization does not alter structural information, arbitrary scaling can affect intensity comparisons. For a more detailed discussion of normalization methods for SIM reconstructions, see **Supplementary Note 2**. Scale bar: 8 μm.

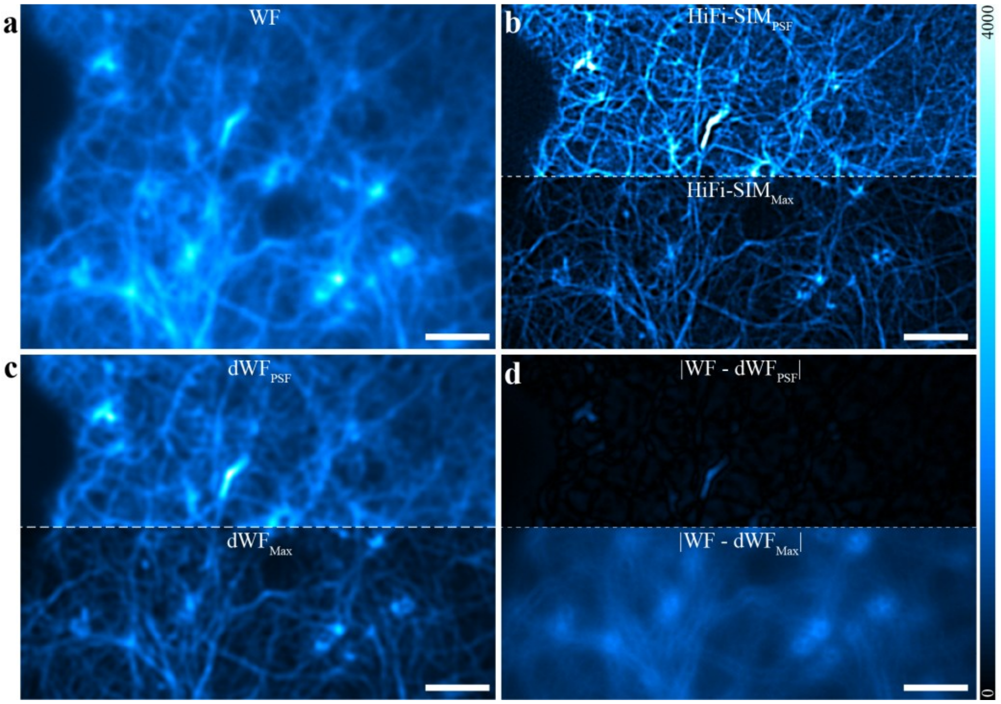


**Supplementary Fig. 2 | Comparison of normalization methods on a single raw SIM dataset.** **a**, WF image of F-actin filaments. **b**, SIM reconstructions obtained using PSF-convolution least-squares normalization (top) and WF max-intensity normalization (bottom), respectively. **c**, Degraded WF images, $\mathrm{dWF_{PSF}}$ (top) and $\mathrm{dWF_{MAX}}$ (bottom), generated by convolving the corresponding SIM images in **b** with the system PSF. **d**, Error maps between the degraded WF images in **c** and the measured WF reference. Compared with WF max-intensity normalization, PSF-convolution least-squares normalization yields smaller errors, indicating better agreement of reconstructed intensities with the WF reference. Scale bars: **a**–**d** 2 µm.

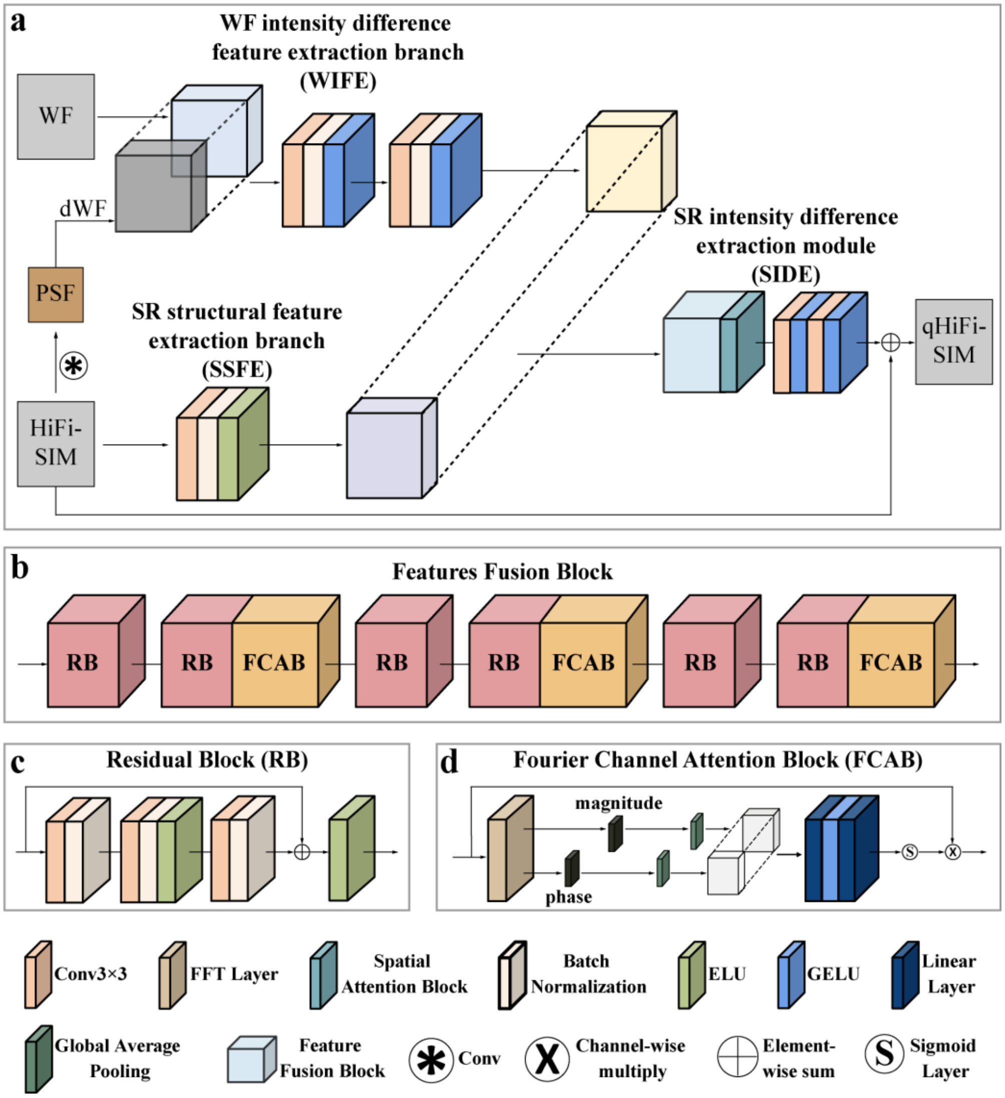


**Supplementary Fig. 3 | Network architecture of the qHiFi-SIM algorithm. a**, The network architecture consists of two parallel feature-extraction branches and a fusion module. **b**, Architecture of the feature fusion module. **c**, Architecture of the Residual Block (**RB**). **d**, Architecture of the Fourier Channel Attention Block (**FCAB**). For a detailed description of the qHiFi-SIM network architecture and its implementation, refer to **Fig. 1a,b** and the **Methods** section.

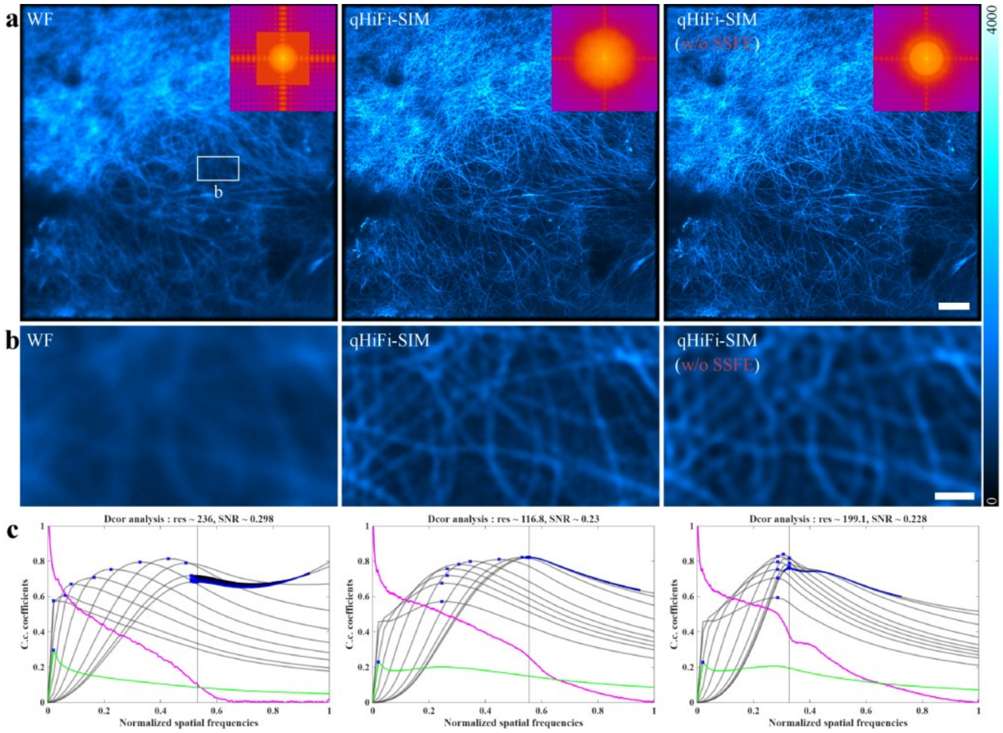


**Supplementary Fig. 4 | Ablation study of the SSFE branch in qHiFi-SIM for reconstruction quality. a**, Images of F-actin structures from the BioSR dataset: WF image (left), qHiFi-SIM reconstruction with the SSFE branch (middle), and qHiFi-SIM reconstruction without the SSFE branch (right). Insets show the corresponding Fourier spectra, revealing a substantial loss of high-frequency components upon SSFE removal. **b**, Magnified views of the white-boxed region in **a**. **c**, Spatial resolutions of the images in panel **a** were estimated by decorrelation analysis[25]. For the representative dataset shown here, the resolution degrades from 116.8 nm to 199.1 nm upon SSFE removal. Across nine independent test datasets, the mean resolution drops from 116.1 ± 2.7 nm to 159.4 ± 22.7 nm (**Supplementary Table 1**). Scale bars: **a** 3 μm; **b** 0.5 μm.

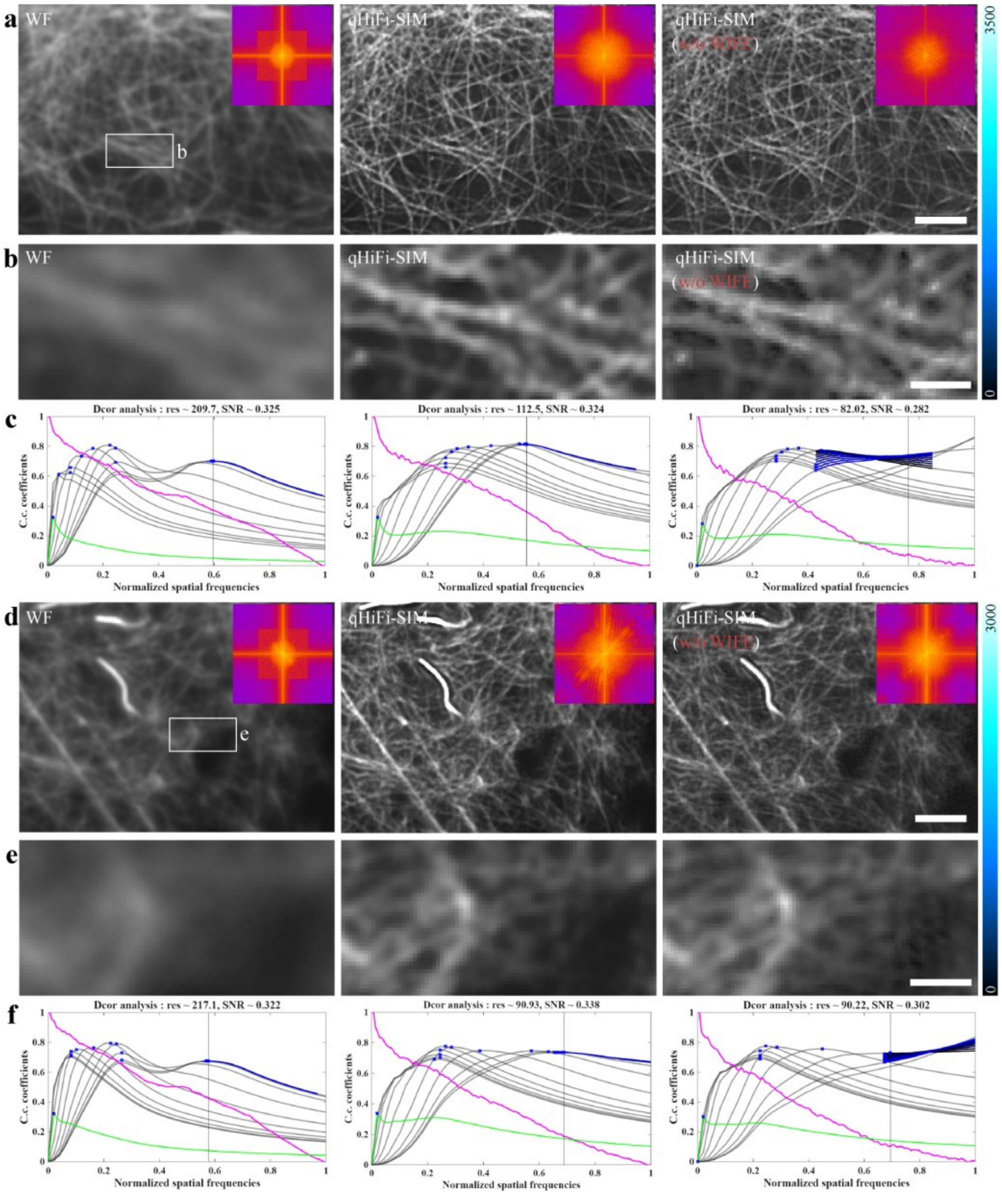


**Supplementary Fig. 5 | Ablation study of the WIFE branch in qHiFi-SIM for reconstruction quality. a**, **d**, Images of two distinct F-actin structures from the BioSR dataset: WF images (left), qHiFi-SIM reconstructions with the WIFE branch (middle), and qHiFi-SIM reconstructions without the WIFE branch (right). Insets show the corresponding Fourier spectra. **b**, **e**, Magnified views of the white-boxed regions in **a** and **d**, respectively. **c**, **f**, Spatial resolutions of the images in

**a** and **d** were estimated by decorrelation analysis, demonstrating that the WIFE branch facilitated the preservation of spatial resolution and, in particular, the improvement of SNR in qHiFi-SIM reconstructions (**Supplementary Table 2**). Quantitative comparisons between the degraded WF images and the measured WF images further demonstrated that the WIFE branch is essential for maintaining intensity linearity and SNR in qHiFi-SIM reconstructions (**Supplementary Tables 3** and **4**). Scale bars: **a** 2 μm; **b**, **e** 0.5 μm; **d** 1 μm.

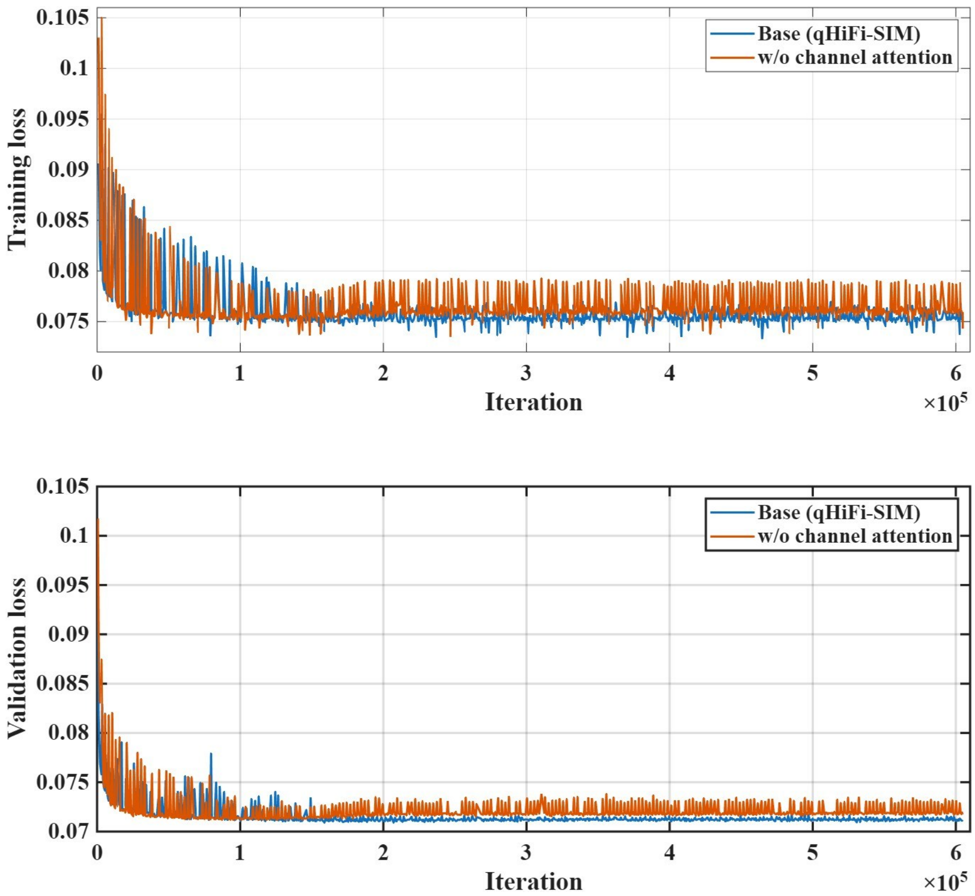


**Supplementary Fig. 6 | Ablation study of the channel attention mechanism: training and validation loss curves.** Training and validation loss curves as a function of iteration for the base qHiFi-SIM model (with channel attention) and the model without channel attention (w/o channel attention). Top, training loss; bottom, validation loss. The base model exhibits a smoother and more stable convergence, whereas the model without channel attention shows notable fluctuations during both training and validation. These results indicate that the channel attention module enhances training stability. All losses were computed on the same training and validation datasets.

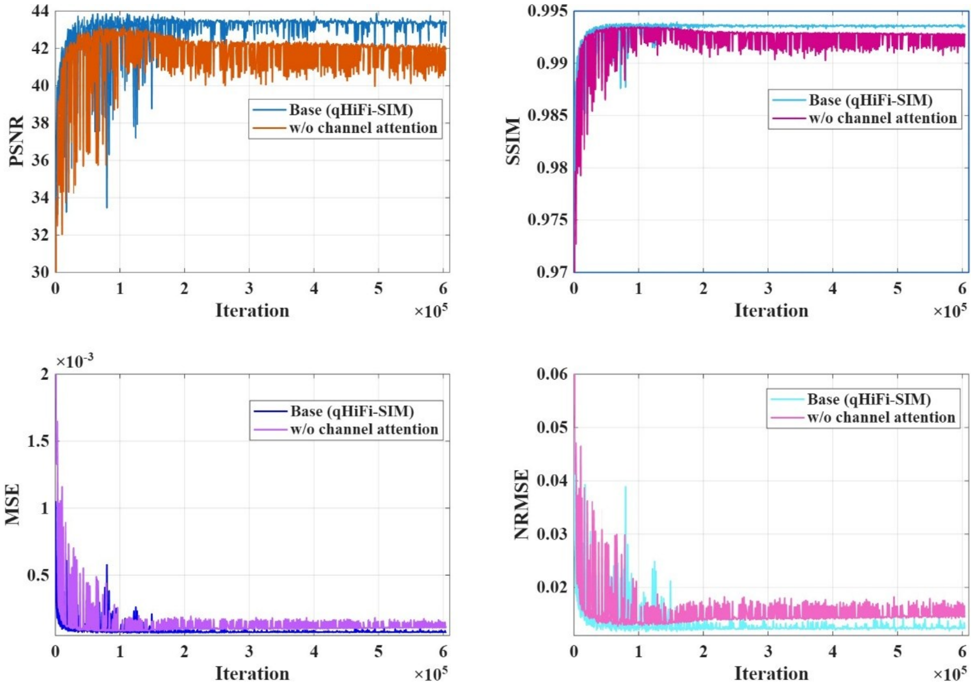


**Supplementary Fig. 7 | Ablation study of the channel attention mechanism: test set performance metrics.** Comparison of the base qHiFi-SIM model (with channel attention) and the variant without channel attention (w/o channel attention) on the test set. Performance was evaluated using PSNR, SSIM, MSE, and NRMSE. The base model consistently outperforms the variant without channel attention across all metrics, whereas the latter exhibits pronounced fluctuations and performance degradation. These results underscore the essential role of the channel attention module in preserving reconstruction fidelity and training stability, as removing it leads to substantial performance loss across all evaluated metrics. All metrics were computed on the independent test set.

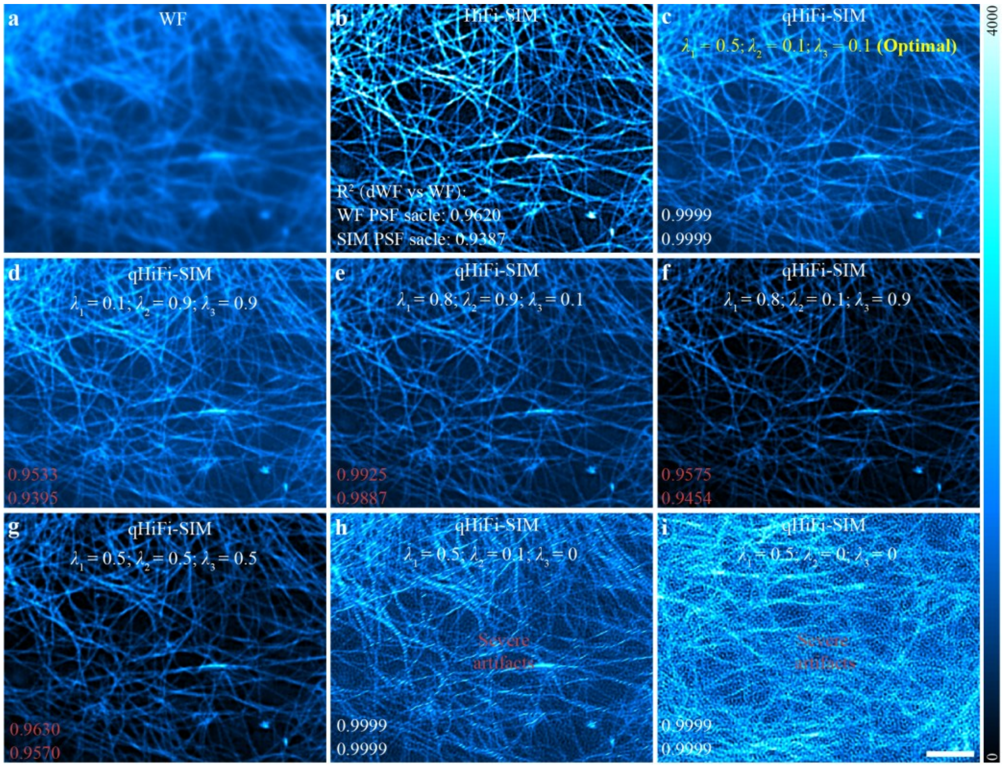


**Supplementary Fig. 8 | Optimization strategy for hyperparameters $\boldsymbol{\lambda_1}$, $\boldsymbol{\lambda_2}$, and $\boldsymbol{\lambda_3}$ in qHiFi-SIM. a,** WF image. **b,** initial HiFi-SIM reconstruction of F-actin structures. **c**, qHiFi-SIM output with the optimal hyperparameter combination ($\lambda_1$ = 0.5, $\lambda_2$ = 0.1, $\lambda_3$ = 0.1). It achieves high intensity linearity, with $R^2$ of 0.9999 at both WF and SIM PSF scales, as well as a PSNR of 47.0851 and SSIM of 0.9909 between the degraded WF (qWF) image and the measured WF image (**Supplementary Table 7**). **d**, Combination ($\lambda_1$ = 0.1, $\lambda_2$ = 0.9, $\lambda_3$ = 0.9): small $\lambda_1$ with large $\lambda_2$ and $\lambda_3$ imposes an overly strong constraint on the HiFi-SIM image, degrading $R^2$ to 0.9533 and 0.9395 at the WF and SIM PSF scales, respectively. **e–g**, Other imbalanced combinations also compromise performance (**Supplementary Table 7**). **e,** Large $\lambda_1$, $\lambda_2$ with small $\lambda_3$ yields $R^2$ of 0.9925 and 0.9887, accompanied by a marked decrease in PSNR and SSIM to 23.8421 and 0.9347. **f,** Small $\lambda_2$ with large $\lambda_1$, $\lambda_2$ yields $R^2$ of 0.9575 and 0.9454. **g,** Balanced $\lambda_1$–$\lambda_3$ values yield $R^2$ of 0.9630 and 0.9570. **h**, Removing the structural similarity constraint ($\lambda_3$ = 0) increases PSNR to 49.92 dB but exhibits severe artifacts. **i**, Removing the full structure loss constraint $L_{\mathrm{struct}}$ ($\lambda_2$ = $\lambda_3$ = 0) further exacerbates structural artifacts. These results demonstrate that a balanced parameter setting is essential for suppressing artifacts and maintaining intensity fidelity. Scale bar: 2 μm.

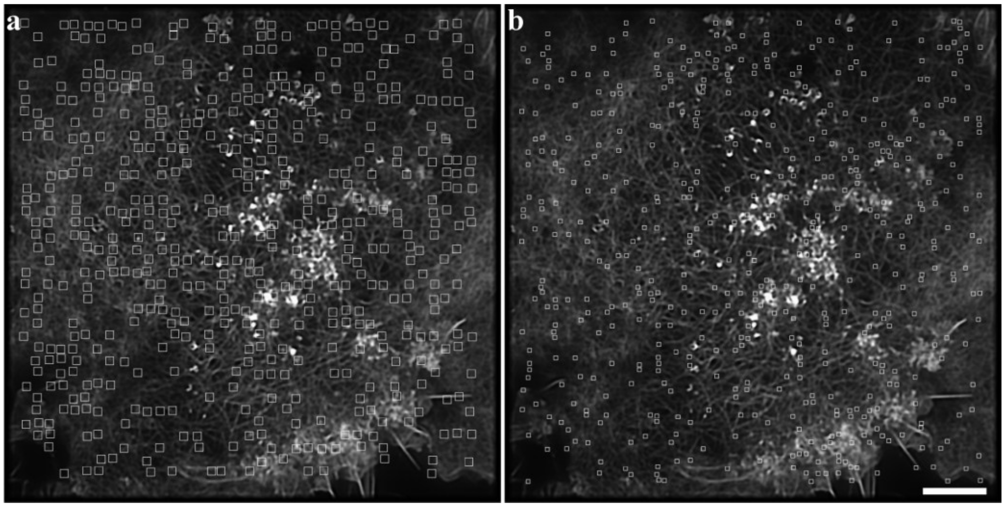


**Supplementary Fig. 9 | Schematic illustration of region-of-interest (ROI) intensity sampling corresponding to Fig. 1.** A total of 500 ROIs were randomly selected at both the WF PSF scale (**a**) and the SIM PSF scale (**b**) to quantify intensity linearity ($R^2$) and mean intensity ratios by summing the intensities within each ROI. Scale bar: 4 μm.

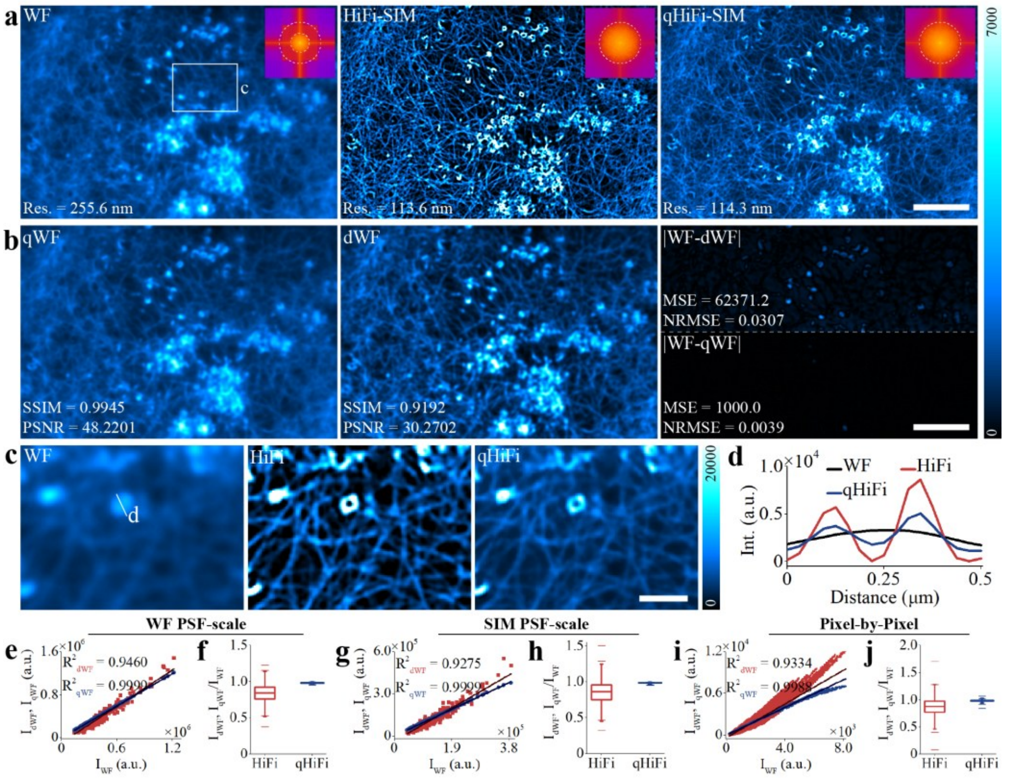


**Supplementary Fig. 10 | Assessing the intensity linearity of qHiFi-SIM reconstructions using experimental F-actin data from the same sample in Fig. 1. a**, WF, initial HiFi-SIM, and qHiFi-SIM reconstructions of the F-actin sample; Insets show the corresponding Fourier spectra, and spatial resolutions estimated by decorrelation analysis are annotated in the lower left. **b**, Degraded WF images qWF (left) and dWF (middle), generated by convolving the corresponding SR images in **a** with the system PSF. Error maps relative to measured WF (right). **c**, Magnified views of the white-box region in **a**. **d**, Fluorescence intensity profiles along the white line in **c**. **e–j**, Quantitative comparison of degraded WF images (dWF and qWF) against the measured WF across three spatial scales: WF PSF scale (**e**, **f**), SIM PSF scale (**g**, **h**), and single-pixel scale (**i**, **j**). $R^2$ values (**e**, **g**, **i**) and box plots of mean intensity ratios (**f**, **h**, **j**) are derived from 500 randomly selected ROIs at each scale. Specifically, at the WF PSF scale, HiFi-SIM yields an $R^2$ of 0.9460 and a mean intensity ratio of 0.843 ± 0.126, whereas qHiFi-SIM achieves an $R^2$ of 0.9999 and a mean intensity ratio of 0.981 ± 0.008. At the SIM PSF scale, HiFi-SIM shows an $R^2$ of 0.9275 and a mean intensity ratio of 0.853 ± 0.161, compared to qHiFi-SIM with an $R^2$ of 0.9999 and a mean intensity ratio of 0.982 ± 0.008. At the single-pixel scale, HiFi-SIM presents an $R^2$ of 0.9334 and a mean intensity ratio of 0.865 ± 0.169, while qHiFi-SIM achieves an $R^2$ of 0.9988 and a mean intensity ratio of 0.980 ± 0.018. Overall, qHiFi-SIM consistently maintains an $R^2$ > 0.998 and a mean intensity ratio ≥ 0.980 across all three scales. Scale bars: **a**, **b** 4 μm; **c** 1 μm.

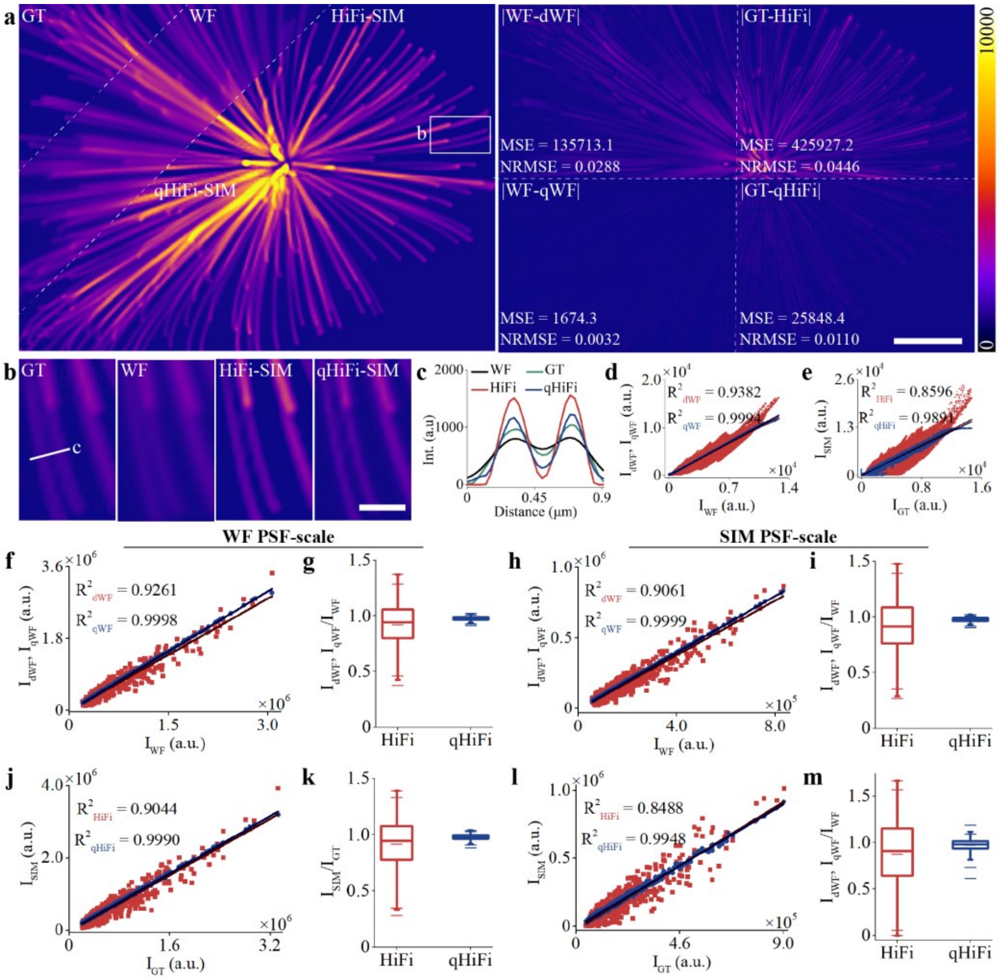


**Supplementary Fig. 11 | Validation of qHiFi-SIM reconstruction intensity linearity using synthetic microtubule simulation data.** The ground truth (GT) image was downloaded from an external dataset[26], and the structured illumination imaging was simulated under conditions matching those of the F-actin sample in the BioSR dataset (emission wavelength: 525 nm; numerical aperture: 1.41; pixel size: 62.6 nm). **a**, GT, WF, HiFi-SIM, and qHiFi-SIM reconstructions of the synthetic microtubule sample (left); error maps of degraded WF images relative to measured WF, as well as HiFi-SIM and qHiFi-SIM relative to GT (right), quantitatively showing that qHiFi-SIM achieves substantially lower MSE and NRMSE compared to HiFi-SIM. **b**, Magnified views of the white-boxed region in **a**. **c**, Fluorescence intensity profiles along the white line in **b**. **d**, **e**, $R^2$ analysis between the degraded WF images (dWF and qWF) and the measured WF image (**d**), and between the SIM reconstructions (HiFi-SIM and qHiFi-SIM) and the GT image (**e**) at the single-pixel scale. **f–m**, Quantitative comparison of degraded WF images and SIM reconstructions across the WF PSF scale (**f**, **g**, **j**, **k**) and SIM PSF scale (**h**, **i**, **l**, **m**). $R^2$ values and box plots of mean intensity ratios at each scale are derived from 500 randomly selected ROIs. Specifically, at the WF PSF scale, HiFi-SIM yields an $R^2$ of 0.9261 and a mean intensity ratio of 0.922 ± 0.185 relative to the measured WF, while qHiFi-SIM achieves an $R^2$ of 0.9998 and a mean intensity ratio of 0.978 ± 0.016 (**f**, **g**); HiFi-SIM yields an $R^2$ of 0.9044 and a mean intensity ratio of 0.915 ± 0.227 relative to GT, while qHiFi-SIM achieves an $R^2$ of 0.9990 and a mean intensity ratio of 0.977 ± 0.025 (**j**, **k**). At the SIM PSF scale, HiFi-SIM shows an $R^2$ of 0.9061 and a mean intensity ratio of 0.911 ± 0.234 relative to the measured WF, while qHiFi-SIM achieves an $R^2$ of 0.9999 and a mean intensity ratio of 0.976 ± 0.019 (**h**, **i**); HiFi-SIM shows an $R^2$ of 0.8488 and a mean intensity ratio of 0.875 ± 0.364 relative to GT, while qHiFi-SIM achieves an $R^2$ of 0.9948 and a mean intensity ratio of 0.970 ± 0.072 (**l**, **m**). Collectively, these results demonstrate that qHiFi-SIM yields mean intensity ratios closer to unity and higher $R^2$ values across all assessed scales, confirming its superior intensity linearity. Scale bars: **a** 2 μm; **b** 0.5 μm.

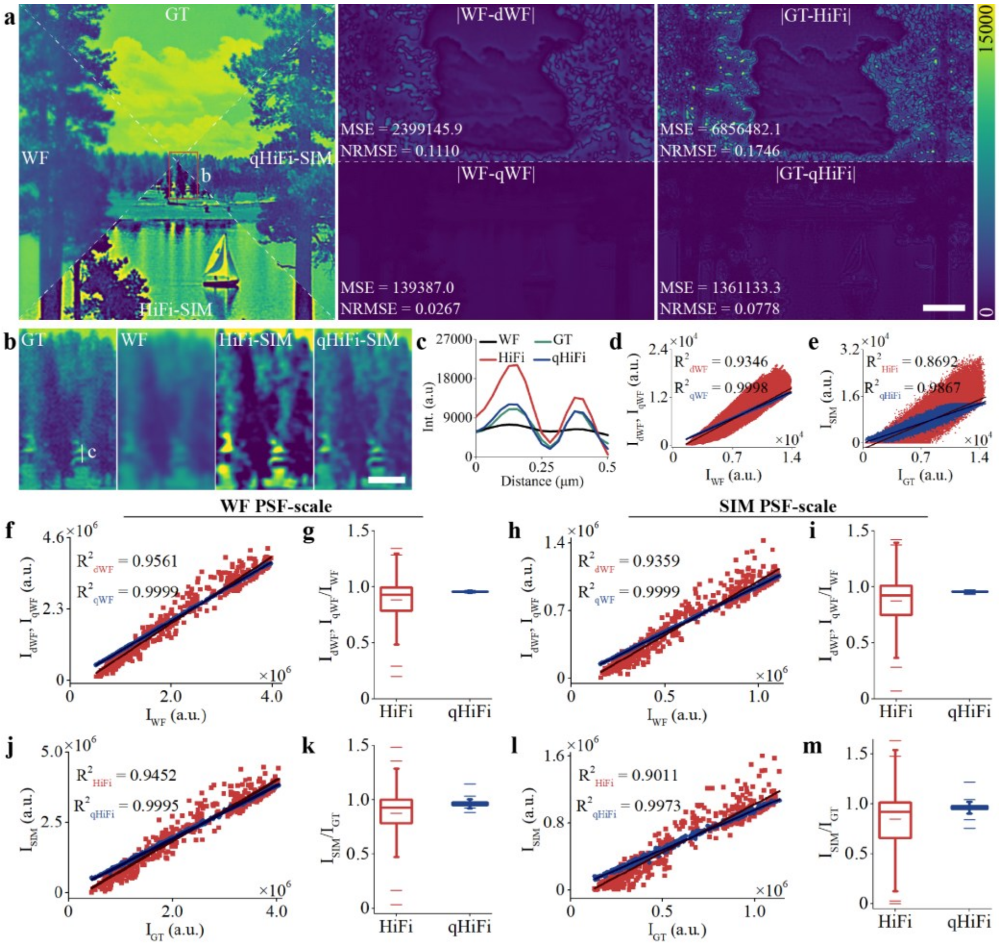


**Supplementary Fig. 12 | Validation of qHiFi-SIM reconstruction intensity linearity using natural scene simulation data.** The GT image was downloaded from an external database[27], and the structured illumination imaging was simulated under conditions matching those of the F-actin sample in the BioSR dataset (emission wavelength: 525 nm; numerical aperture: 1.41; pixel size: 62.6 nm). **a**, GT, WF, HiFi-SIM, and qHiFi-SIM reconstructions of the natural scene image (left); error maps of degraded WF images relative to the measured WF (middle), as well as HiFi-SIM and qHiFi-SIM relative to GT (right), showing that qHiFi-SIM achieves substantially lower MSE and NRMSE compared to HiFi-SIM. **b**, Magnified views of the red-boxed region in **a**. **c**, Fluorescence intensity profiles along the white line in **b**. **d**, **e**, $R^2$ analysis between the degraded WF images (dWF and qWF) and the measured WF image (**d**), and between the SIM reconstructions (HiFi-SIM and qHiFi-SIM) and the GT image (**e**) at the single-pixel scale. **f**–**m**, Quantitative comparison of degraded WF images and SIM reconstructions across the WF PSF scale (**f**, **g**, **j**, **k**) and SIM PSF scale (**h**, **i**, **l**, **m**). $R^2$ values and box plots of mean intensity ratios at each scale are derived from 500 randomly selected ROIs. Specifically, at the WF PSF scale, HiFi-SIM yields an $R^2$ of 0.9561 and a mean intensity ratio of 0.884 ± 0.194 relative to the measured WF, while qHiFi-SIM achieves an $R^2$ of 0.9999 and a mean intensity ratio of 0.958 ± 0.004 (**f**, **g**); HiFi-SIM yields an $R^2$ of 0.9452 and a mean intensity ratio of 0.876 ± 0.225 relative to GT, while qHiFi-SIM achieves an $R^2$ of 0.9995 and a mean intensity ratio of 0.964 ± 0.021 (**j**, **k**). At the SIM PSF scale, HiFi-SIM shows an $R^2$ of 0.9359 and a mean intensity ratio of 0.874 ± 0.228 relative to the measured WF, while qHiFi-SIM achieves an $R^2$ of 0.9999 and a mean intensity ratio of 0.958 ± 0.005 (**h**, **i**); HiFi-SIM shows an $R^2$ of 0.9011 and a mean intensity ratio of 0.847 ± 0.309 relative to GT, while qHiFi-SIM achieves an $R^2$ of 0.9973 and a mean intensity ratio of 0.961 ± 0.037 (**l**, **m**). Scale bars: **a** 4 μm; **b** 1 μm.

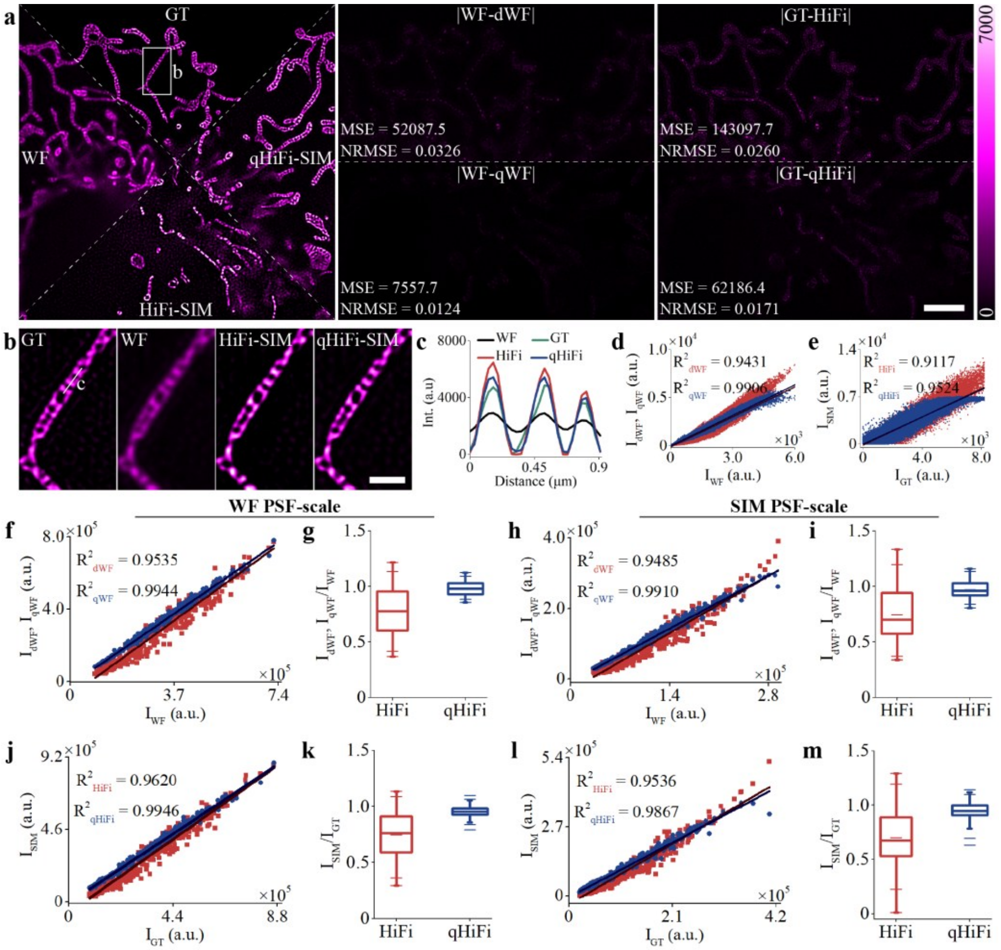


**Supplementary Fig. 13 | Validation of qHiFi-SIM reconstruction intensity linearity using mitochondrial cristae simulation data.** The GT image (mitochondrial cristae structures) was acquired from live U2OS cells using a Polar-SIM system, and the structured illumination imaging was simulated under the following conditions: emission wavelength, 525 nm; numerical aperture, 1.49; pixel size, 65 nm. **a**, GT, WF, HiFi-SIM, and qHiFi-SIM reconstructions of the mitochondrial cristae sample (left); error maps of degraded WF images relative to measured WF (middle), as well as HiFi-SIM and qHiFi-SIM relative to GT (right). **b**, Magnified views of the white-boxed region in **a**. **c**, Fluorescence intensity profiles along the white line in **b**. **d**, **e**, $R^2$ analysis between the degraded WF images (dWF and qWF) and the measured WF image (**d**), and between the SIM reconstructions (HiFi-SIM and qHiFi-SIM) and the GT image (**e**) at the single-pixel scale. **f**–**m**, Quantitative comparison of degraded WF images and SIM reconstructions across the WF PSF scale (**f**, **g**, **j**, **k**) and SIM PSF scale (**h**, **i**, **l**, **m**). $R^2$ values and box plots of mean intensity ratios at each scale are derived from 500 randomly selected ROIs. Specifically, at the WF PSF scale, HiFi-SIM yields an $R^2$ of 0.9535 and a mean intensity ratio of 0.777 ± 0.197 relative to the measured WF, while qHiFi-SIM achieves an $R^2$ of 0.9944 and a mean intensity ratio of 0.983 ± 0.056 (**f, g**); HiFi-SIM yields an $R^2$ of 0.9620 and a mean intensity ratio of 0.747 ± 0.191 relative to GT, while qHiFi-SIM achieves an $R^2$ of 0.9946 and a mean intensity ratio of 0.956 ± 0.043 (**j**, **k**). At the SIM PSF scale, HiFi-SIM shows an $R^2$ of 0.9485 and a mean intensity ratio of 0.748 ± 0.218 relative to the measured WF, while qHiFi-SIM achieves an $R^2$ of 0.9910 and a mean intensity ratio of 0.974 ± 0.069 (**h**, **i**); HiFi-SIM shows an $R^2$ of 0.9536 and a mean intensity ratio of 0.702 ± 0.239 relative to GT, while qHiFi-SIM achieves an $R^2$ of 0.9867 and a mean intensity ratio of 0.945 ± 0.079 (**l**, **m**). Despite the distinct differences in sample structures and system parameters compared to the training conditions, qHiFi-SIM consistently maintains an $R^2$ > 0.986 and a mean intensity ratio ≥ 0.945 at the SIM PSF scale, demonstrating its robust transferability for intensity linearity. Scale bars: **a** 4 μm; **b** 1 μm.

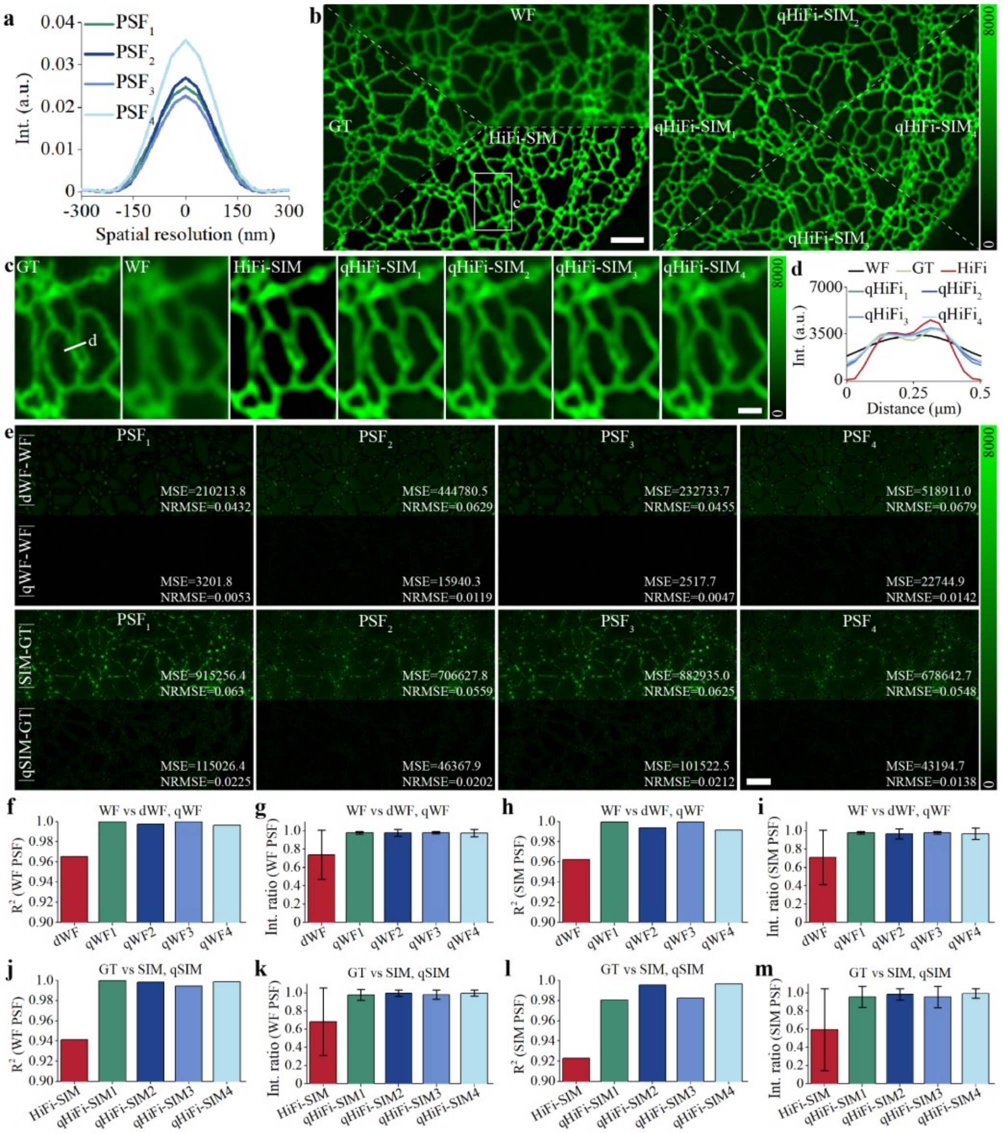


**Supplementary Fig. 14 | Evaluating the impact of PSF mismatch on qHiFi-SIM reconstruction intensity linearity using simulated ER data.** The GT ER image was obtained from Wiener-SIM reconstruction of the BioSR dataset. **a**, Comparison of four PSFs. PSF$_1$ matches the training conditions (emission wavelength, 525 nm; numerical aperture, 1.41; pixel size, 62.6 nm); PSF$_2$–PSF$_4$ correspond to the imaging parameters of the commercial Polar-SIM (emission wavelength, 525 nm; numerical aperture, 1.49; pixel size, 65 nm), home-built SIM-FRET (emission wavelength, 600 nm, numerical aperture, 1.49, pixel size, 72.2

nm), and GE Healthcare 3D-SIM (emission wavelength, 527 nm; numerical aperture, 1.42; pixel size, 78.6 nm) systems used in this study, respectively. These four PSFs were generated using the same theoretical PSF model as in the HiFi-SIM algorithm. The structured illumination imaging was simulated under conditions matching $PSF_1$. **b**, GT, WF, and HiFi-SIM reconstructions of the ER sample (left); qHiFi-SIM reconstructions using the four PSFs (right). **c**, Magnified views of the white-boxed region in **b**. **d**, Fluorescence intensity profiles along the white line in **c**. **e**, Error maps of degraded WF images (dWF and qWF) relative to the measured WF (upper row), and SIM reconstructions (HiFi-SIM and qHiFi-SIM) relative to GT (lower row) under the four PSF conditions. **f**–**i**, Quantitative comparison of $R^2$ values (**f**, **h**) and mean intensity ratios (**g**, **i**) of degraded WF images relative to the measured WF at the WF PSF scale (**f**, **g**) and SIM PSF scale (**h**, **i**); $R^2$ values and box plots of mean intensity ratios at each scale are derived from 500 randomly selected ROIs. **j**–**m**, Quantitative comparison of $R^2$ values (**j**, **l**) and mean intensity ratios (**k**, **m**) of SIM reconstructions relative to GT at the WF PSF scale (**j**, **k**) and SIM PSF scale (**l**, **m**). Overall, qHiFi-SIM consistently outperforms HiFi-SIM across all metrics at both scales even under sample structure variations and PSF-mismatched conditions, maintaining an $R^2 > 0.989$ and a mean intensity ratio ≥ 0.966 at the SIM PSF scale, demonstrating its robust immunity to imaging PSF mismatches. Scale bars: **b**, **e** 2 μm; **c** 0.5 μm.

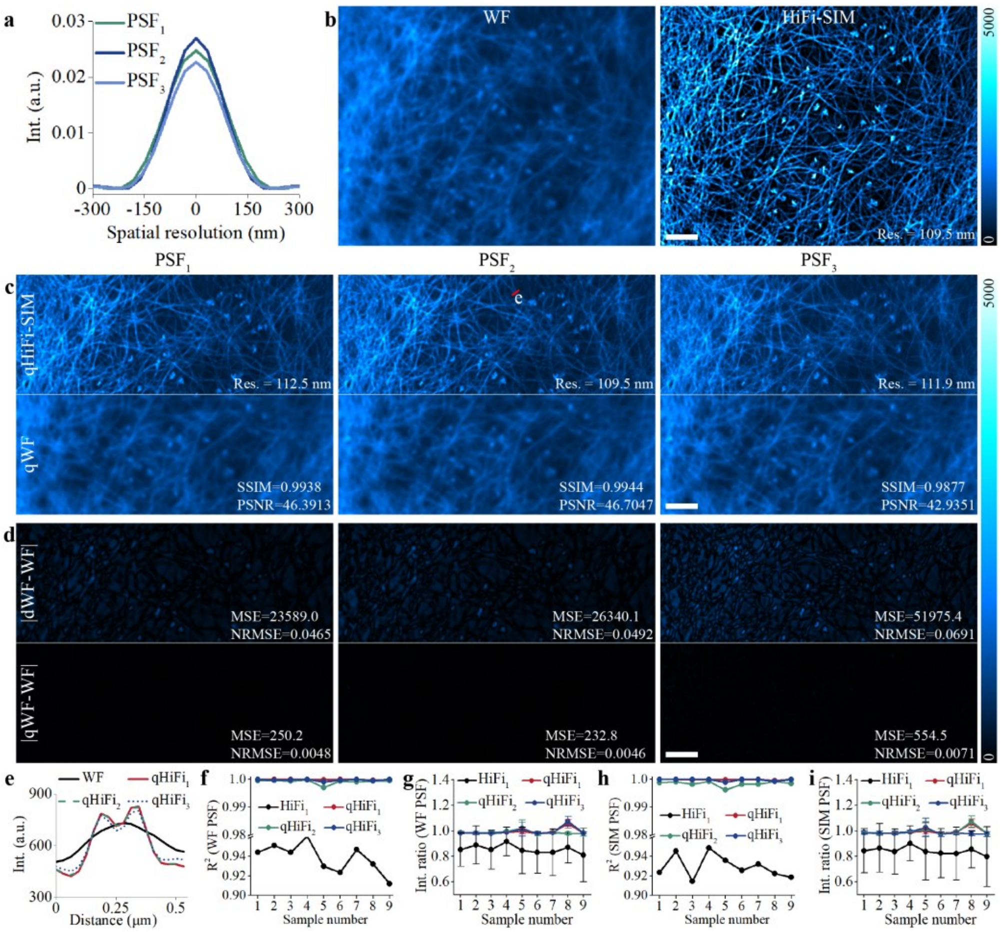


**Supplementary Fig. 15 | Evaluating the impact of PSF mismatch on qHiFi-SIM reconstruction intensity linearity using experimentally measured F-actin data.** The data were obtained from the BioSR dataset[20]. **a**, Comparison of three PSFs. $PSF_1$ matches the training conditions (emission wavelength, 525 nm; numerical aperture, 1.41; pixel size, 62.6 nm); $PSF_2$ and $PSF_3$ correspond to the imaging parameters of the commercial Polar-SIM (Em 525 nm, NA 1.49, pixel size 65 nm) and the home-built SIM-FRET (Em 600 nm, NA 1.49, pixel size 72.2 nm) systems used in this study, respectively. These three PSFs were generated using the same theoretical PSF model as in the HiFi-SIM algorithm. **b**, WF image (left) and corresponding HiFi-SIM reconstruction using $PSF_1$ (right). The spatial resolution of the HiFi-SIM image was determined as 109.5 nm through decorrelation analysis. **c**, qHiFi-SIM reconstructions (upper row) of the HiFi-SIM image in **b** using $PSF_1$, $PSF_2$, and $PSF_3$, and their corresponding degraded WF images (lower row). Comparable performance among the three PSFs was observed in terms of achieved resolution, PSNR, and SSIM. **d**, Error maps of the degraded WF images (dWF and qWF) relative to the measured WF. **e**, Fluorescence intensity profiles along the red line in **c**. **f**–**i**, Quantitative statistics of $R^2$ values (**f**, **h**) and mean intensity ratios (**g**, **i**) across nine test samples at the WF PSF scale (**f**, **g**) and SIM PSF scale (**h**, **i**). Overall, qHiFi-SIM consistently outperforms HiFi-SIM across all metrics at both scales even under PSF-mismatched conditions, maintaining an $R^2 > 0.996$ and a mean intensity ratio $\geq 0.974$ at the SIM PSF scale, demonstrating its robust immunity to imaging PSF mismatches. Collectively, although an experimentally matched PSF provides optimal accuracy, a theoretical PSF based on the imaging parameters is sufficient for high-quality reconstruction, circumventing specialized calibration and improving the accessibility and practicality of qHiFi-SIM. Scale bars: **b**–**d** 2 μm.

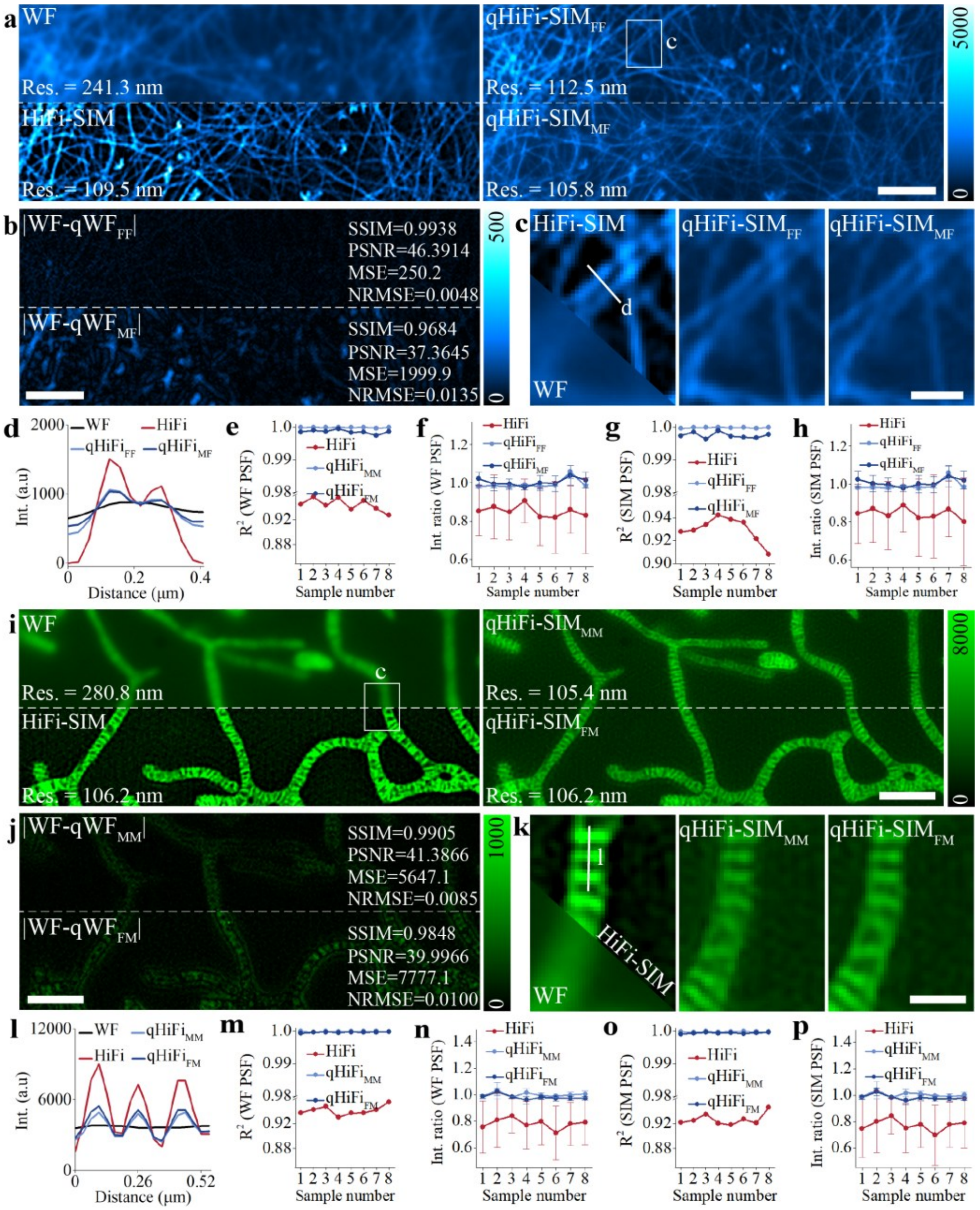


**Supplementary Fig. 16 | Evaluating the transferability of qHiFi-SIM through cross-training and validation.** To assess cross-structure transferability, two independent qHiFi-SIM networks were trained on F-actin data (BioSR: emission wavelength, 525 nm; numerical aperture, 1.41; pixel size, 62.6 nm) and mitochondrial cristae data from U2OS cells (commercial Polar-SIM: emission wavelength, 525 nm; numerical aperture, 1.49; pixel size, 65 nm), and then applied to reconstruct eight samples of each type. Transferability was quantified by comparing homologous versus heterologous reconstructions. **a**, For an F-actin sample: WF (upper left), HiFi-SIM (lower left), and qHiFi-SIM reconstructions trained on F-actin dataset (qHiFi-$SIM_{FF}$; upper right) and mitochondrial cristae dataset (qHiFi-$SIM_{MF}$; lower right). **b**, Error maps and quantitative metrics between the measured WF and the degraded WF images ($qWF_{FF}$ and $qWF_{MF}$) of the qHiFi-SIM reconstructions in **a**. **c**, Magnified views of the white-boxed region in **a**. **d**, Fluorescence intensity profiles along the white line in **c**. **e**–**h**, $R^2$ (**e**, **g**) and mean intensity ratio (**f**, **h**) statistics across eight F-actin test samples at the WF PSF (**e**, **f**) and SIM PSF (**g**, **h**) scales. Both networks maintained $R^2 > 0.995$ and mean intensity ratios ≥ 0.979 at the SIM PSF scale, indicating comparable intensity linearity between the two networks. **i**, For a mitochondrial cristae sample: WF (upper left), HiFi-SIM (lower left), and qHiFi-SIM reconstructions trained on mitochondrial cristae dataset (qHiFi-$SIM_{MM}$; upper right) and F-actin dataset (qHiFi-$SIM_{FM}$; lower right). **j**, Corresponding error maps and metrics for the degraded WF images ($qWF_{MM}$ and $qWF_{FM}$) of the qHiFi-SIM reconstructions in **i**. **k**, Magnified views of the white-boxed region in **i**. **l**, Fluorescence intensity profiles along the white line in **k**. **m**–**p**, $R^2$ (**m**, **o**) and mean intensity ratio (**n**, **p**) statistics across eight mitochondrial cristae test samples at the WF PSF (**m**, **n**) and SIM PSF (**o**, **p**) scales. Both networks maintained $R^2 > 0.999$ and mean intensity ratios ≥ 0.972 at the SIM PSF scale. Collectively, qHiFi-SIM exhibits consistently high intensity linearity and reconstruction quality regardless of homologous or heterologous training, confirming robust transferability across sample structures and imaging conditions. Scale bars: **a**, **b**, **i**, **j** 2 μm; **c**, **k** 0.5 μm.

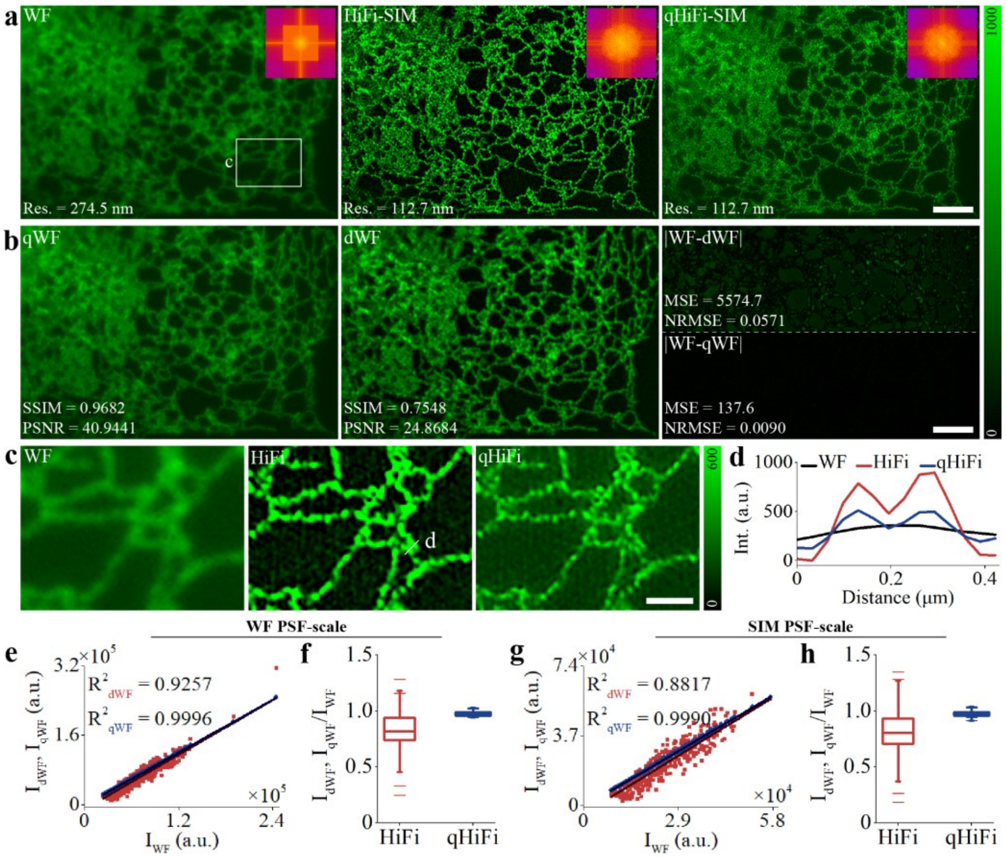


**Supplementary Fig. 17 | Assessing the intensity linearity of qHiFi-SIM reconstruction using endoplasmic reticulum structures in U2OS live cells.** Endoplasmic reticulum (ER) data were acquired using a home-built 2D-SIM system with the following parameters: emission wavelength, 525 nm; numerical aperture, 1.49; pixel size, 65 nm. **a**, WF, HiFi-SIM, and qHiFi-SIM reconstructions of the ER sample; Insets show the corresponding Fourier spectra, and spatial resolutions estimated by decorrelation analysis are annotated in the lower left. **b**, Degraded WF images qWF (left) and dWF (middle), generated by convolving the corresponding SR images in **a** with the system PSF; error maps relative to measured WF (right). **c**, Magnified views of the white-box region in **a**. **d**, Fluorescence intensity profiles along the white line in **c**. **e–h**, Quantitative comparison of degraded WF images (dWF and qWF)

relative to the measured WF across the WF PSF scale (**e**, **f**) and SIM PSF scale (**g**, **h**). $R^2$ values (**e**, **g**) and box plots of mean intensity ratios (**f**, **h**) are derived from 500 randomly selected ROIs at each scale. Specifically, at the WF PSF scale, HiFi-SIM yields an $R^2$ of 0.9257 and a mean intensity ratio of 0.824 ± 0.170, whereas qHiFi-SIM achieves an $R^2$ of 0.9996 and a mean intensity ratio of 0.977 ± 0.019 (**e**, **f**). At the SIM PSF scale, HiFi-SIM shows an $R^2$ of 0.8817 and a mean intensity ratio of 0.804 ± 0.208, compared to qHiFi-SIM with an $R^2$ of 0.9990 and a mean intensity ratio of 0.976 ± 0.021 (**g**, **h**). Overall, despite the distinct differences in sample structures and system parameters compared to the training conditions, qHiFi-SIM consistently maintains an $R^2 > 0.999$ and a mean intensity ratio ≥ 0.976 at the SIM PSF scale, demonstrating its robust transferability for intensity linearity. Scale bars: **a**, **b** 3 μm; **c** 1 μm.

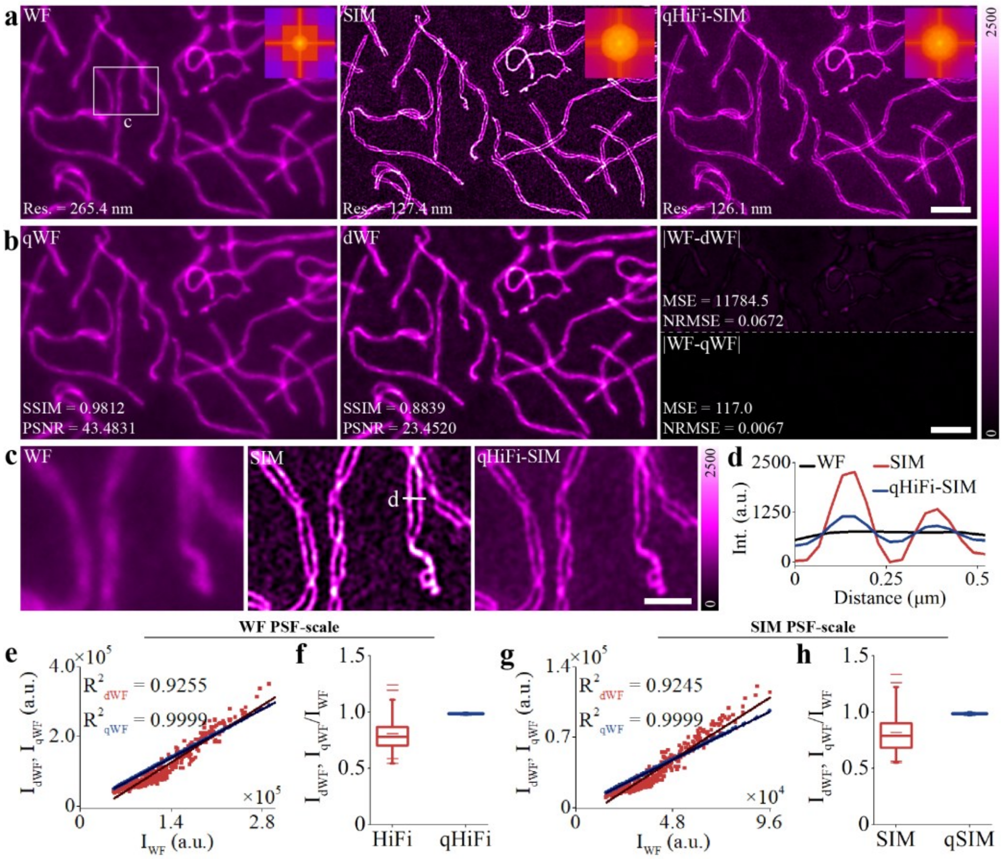


**Supplementary Fig. 18 | Assessing the intensity linearity of qHiFi-SIM reconstructions using synaptonemal complex sample.** Synaptonemal complex data were acquired using a commercial Polar-SIM system (Airy Technologies Co., Ltd.) with the following parameters: emission wavelength, 605 nm; numerical aperture, 1.49; pixel size, 65 nm. SIM images were reconstructed using the built-in algorithm of the system. **a**, WF, SIM, and qHiFi-SIM reconstructions of the synaptonemal complex sample; Insets show the corresponding Fourier spectra, and spatial resolutions estimated by decorrelation analysis are annotated in the lower left. **b**, Degraded WF images qWF (left) and dWF (middle), generated by convolving the corresponding SR images in a with the system PSF; error maps relative to measured WF (right). **c**, Magnified views of the white-boxed region in **a**. **d**, Fluorescence intensity profiles along the white line in **c**. **e**–**h**, Quantitative comparison of degraded WF images (dWF and qWF) relative to the measured WF across the WF PSF scale (**e**, **f**) and SIM PSF scale (**g**, **h**). $R^2$ values (**e**, **g**) and box plots of mean intensity ratios (**f**, **h**) are derived from 500 randomly selected ROIs at each scale. Specifically, at the WF PSF scale, the built-in algorithm (SIM) yields an $R^2$ of 0.9255 and a mean intensity ratio of 0.808 ± 0.138, whereas qHiFi-SIM achieves an $R^2$ of 0.9999 and a mean intensity ratio of 0.987 ± 0.004 (**e**, **f**). At the SIM PSF scale, the built-in algorithm (SIM) shows an $R^2$ of 0.9245 and a mean intensity ratio of 0.819 ± 0.174, compared to qHiFi-SIM with an $R^2$ of 0.9999 and a mean intensity ratio of 0.987 ± 0.005 (**g**, **h**). Overall, despite the differences in sample structures, imaging conditions, and reconstruction algorithms compared to the training conditions, qHiFi-SIM consistently maintains an $R^2 > 0.999$ and a mean intensity ratio ≥ 0.987 at the SIM PSF scale, confirming its excellent transferability across different experimental setups and sample types. Scale bars: **a**, **b** 3 μm; **c** 1 μm.

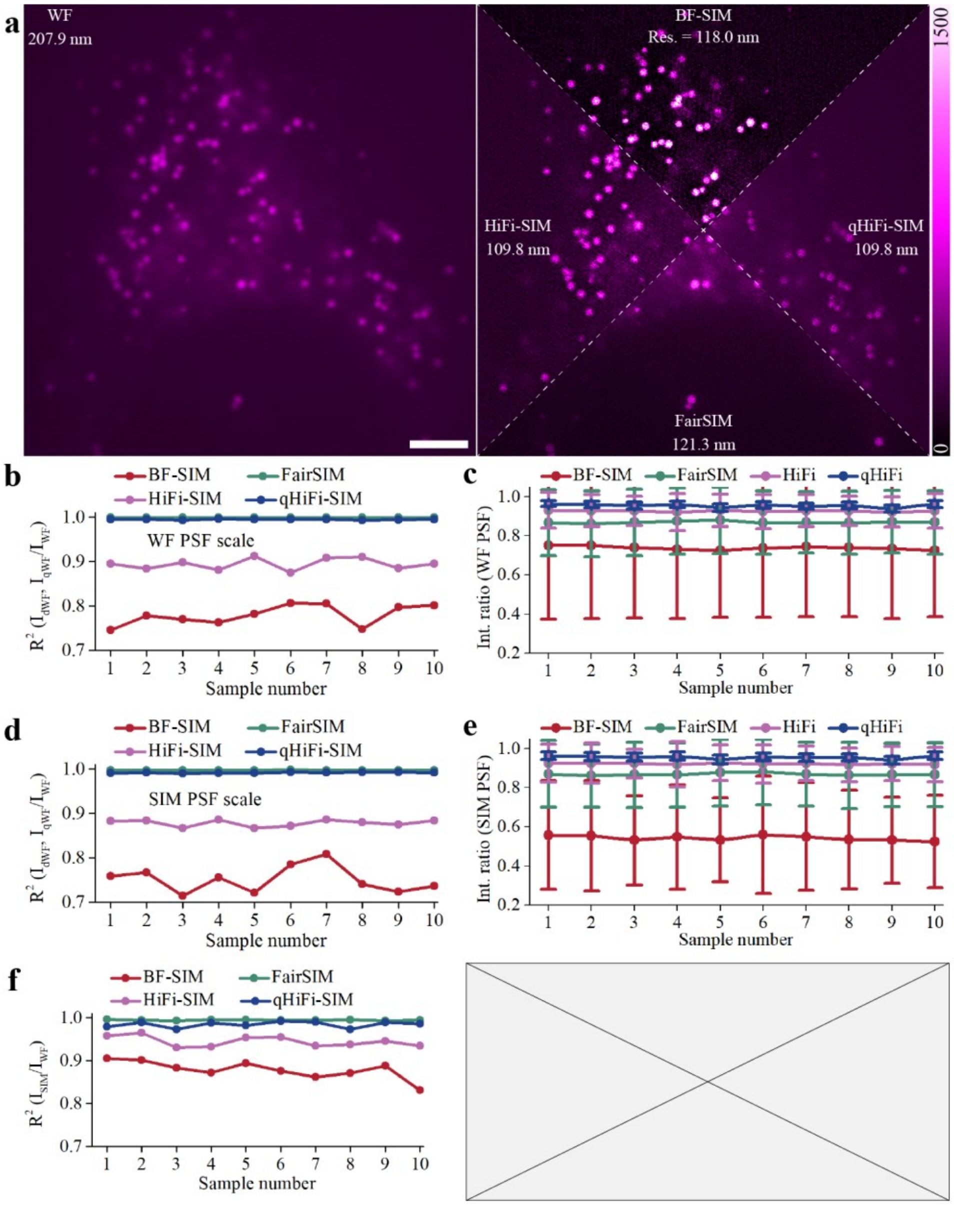


**Supplementary Fig. 19 | Evaluating the intensity linearity of different SIM reconstruction algorithms using an open-source lipid droplet dataset.** The lipid droplet data were sourced from an open-access repository established for BF-SIM benchmarking[16]**.** **a**, WF image of the lipid droplet sample (left), and reconstructions by BF-SIM, fairSIM, HiFi-SIM, and qHiFi-SIM (right, separated by white dashed lines), with corresponding spatial resolutions annotated in each region. **b**–**e**,

Quantitative comparison of $R^2$ values (**b**, **d**) and mean intensity ratios (**c**, **e**) of reconstructions from four different algorithms at the WF PSF scale (**b**, **c**) and SIM PSF scale (**d**, **e**). These metrics at each scale are derived from 500 randomly selected ROIs across ten consecutive time points. qHiFi-SIM achieves $R^2$ values comparable to fairSIM (> 0.990) at both scales, while exhibiting the mean intensity ratios closest to unity; in contrast, BF-SIM shows the lowest performance in both metrics, implying the largest relative fluctuation in its reconstruction intensity. **f**, $R^2$ values calculated between the SIM images and the measured WF images within the segmented lipid droplet regions, following the analysis approach used in BF-SIM[16]. The results show that BF-SIM exhibits the lowest $R^2$ values, while qHiFi-SIM achieves $R^2$ values comparable to fairSIM (≥ 0.973), demonstrating its superior reconstruction intensity linearity. Scale bar: 4 μm.

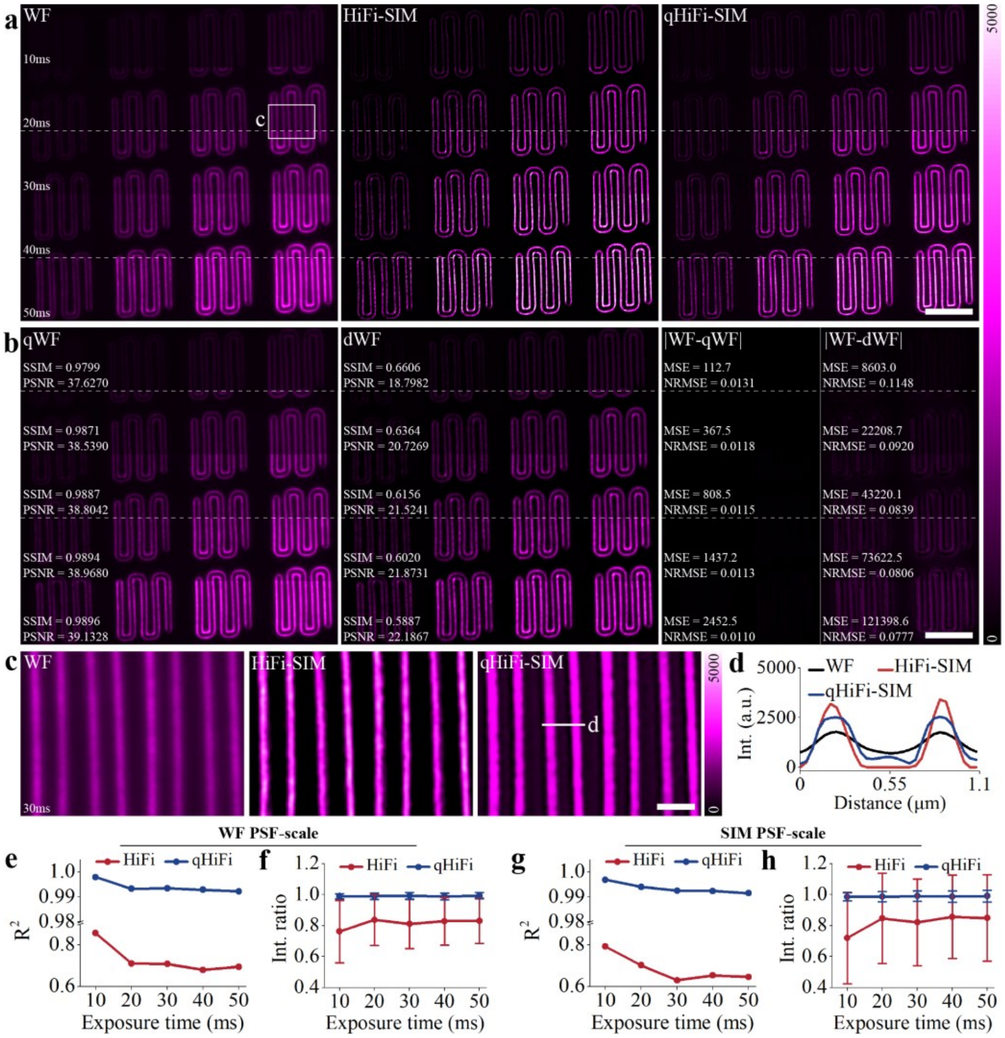


**Supplementary Fig. 20 | Evaluating the intensity linearity of qHiFi-SIM reconstructions under gradient exposure times using an Argo-SIM standard slide.** SIM imaging of the paperclip-shaped pattern on an Argo-SIM standard slide was performed at gradient exposure times ranging from 10 to 50 ms using a DeltaVision OMX SR system (GE Healthcare; emission wavelength, 527 nm; numerical aperture, 1.42; pixel size, 78.6 nm). **a**, WF (left), HiFi-SIM (middle), and qHiFi-SIM (right) reconstructions of the paperclip-shaped pattern at gradient exposure times (10–50 ms). **b**, Corresponding degraded WF images qWF (left) and dWF (middle), along with their error maps relative to the measured

WF (right). **c**, Magnified views of the white-boxed region in **a**. **d**, Fluorescence intensity profiles along the white line in **c**. **e**–**h**, Quantitative comparison of $R^2$ values (**e**, **g**) and mean intensity ratios (**f**, **h**) at the WF PSF scale (**e**, **f**) and SIM PSF scale (**g**, **h**) across gradient exposure times. These metrics at each scale are derived from 500 randomly selected ROIs. The results show that the $R^2$ and mean intensity ratios of HiFi-SIM reconstructions exhibit notable fluctuations across different exposure times and are substantially lower than those of qHiFi-SIM; in contrast, qHiFi-SIM reconstructions consistently achieve an $R^2 > 0.991$ across all five exposure times, with a mean intensity ratio ≥ 0.985 at the SIM PSF scale. Scale bars: **a**, **b** 6 μm; **c** 1 μm.

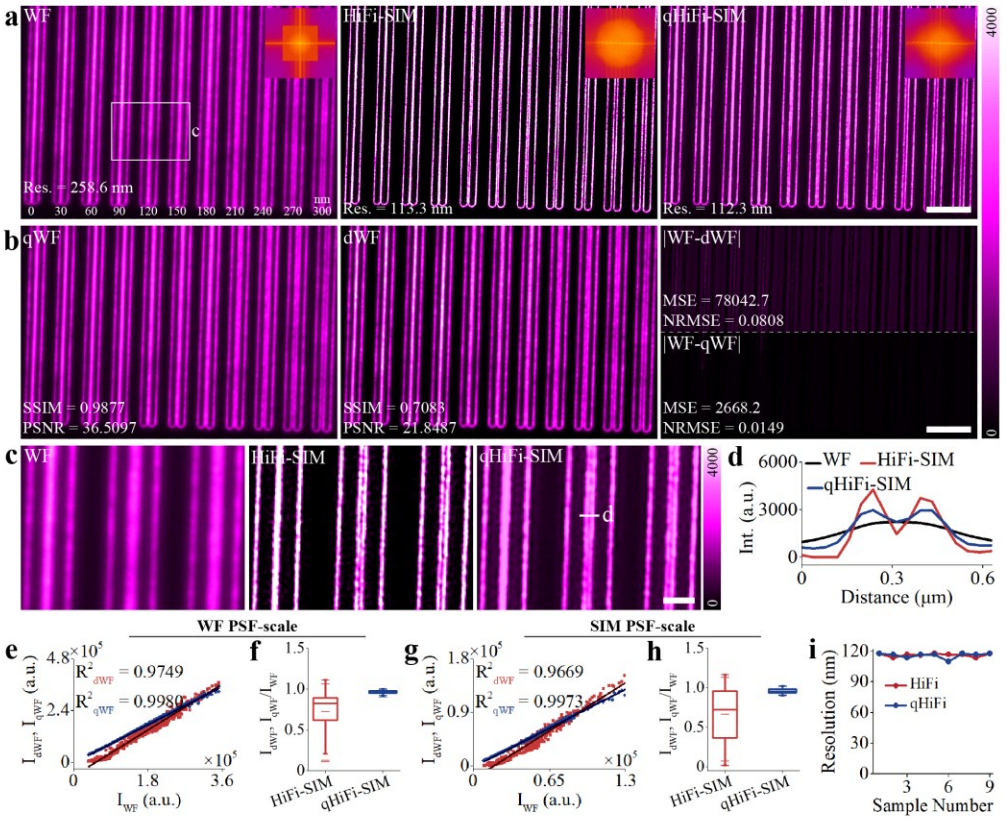


**Supplementary Fig. 21 | Evaluating the spatial resolution of qHiFi-SIM reconstructions using a resolution line-pair pattern on an Argo-SIM standard slide.** The line-pair pattern on an Argo-SIM standard slide was imaged using a DeltaVision OMX SR system (GE Healthcare; emission wavelength, 527 nm; numerical aperture, 1.42; pixel size, 78.6 nm). **a**, WF (left), HiFi-SIM (middle), and qHiFi-SIM (right) reconstructions of the line-pair pattern; Insets show the corresponding Fourier spectra, and spatial resolutions estimated by decorrelation analysis are annotated in the lower left. **b**, Corresponding degraded WF images qWF (left) and dWF (middle), along with their error maps relative to the measured WF (right), quantitatively showing that qWF achieves substantially higher SSIM/PSNR compared to dWF. **c**, Magnified views of the white-boxed region in **a**. **d**, Fluorescence intensity profiles along the white line in **c**. The results demonstrate that both HiFi-SIM and qHiFi-

SIM are capable of resolving line pairs with a 120 nm spacing, which is further corroborated by the resolution estimates from decorrelation analysis. **e**–**h**, Quantitative comparison of $R^2$ values (**e**, **g**) and mean intensity ratios (**f**, **h**) of degraded WF images (dWF and qWF) relative to the measured WF at the WF PSF scale (**e**, **f**) and SIM PSF scale (**g**, **h**). These metrics at each scale are derived from 500 randomly selected ROIs. The results show that qHiFi-SIM reconstruction consistently achieves an $R^2 > 0.997$, with mean intensity ratios of approximately 0.960 at the SIM PSF scale. **i**, Statistical comparison of spatial resolution between HiFi-SIM and qHiFi-SIM reconstructions across nine test F-actin samples. Collectively, qHiFi-SIM achieves comparable spatial resolution to HiFi-SIM while exhibiting superior intensity linearity. Scale bars: **a**, **b** 4 μm; **c** 1 μm.

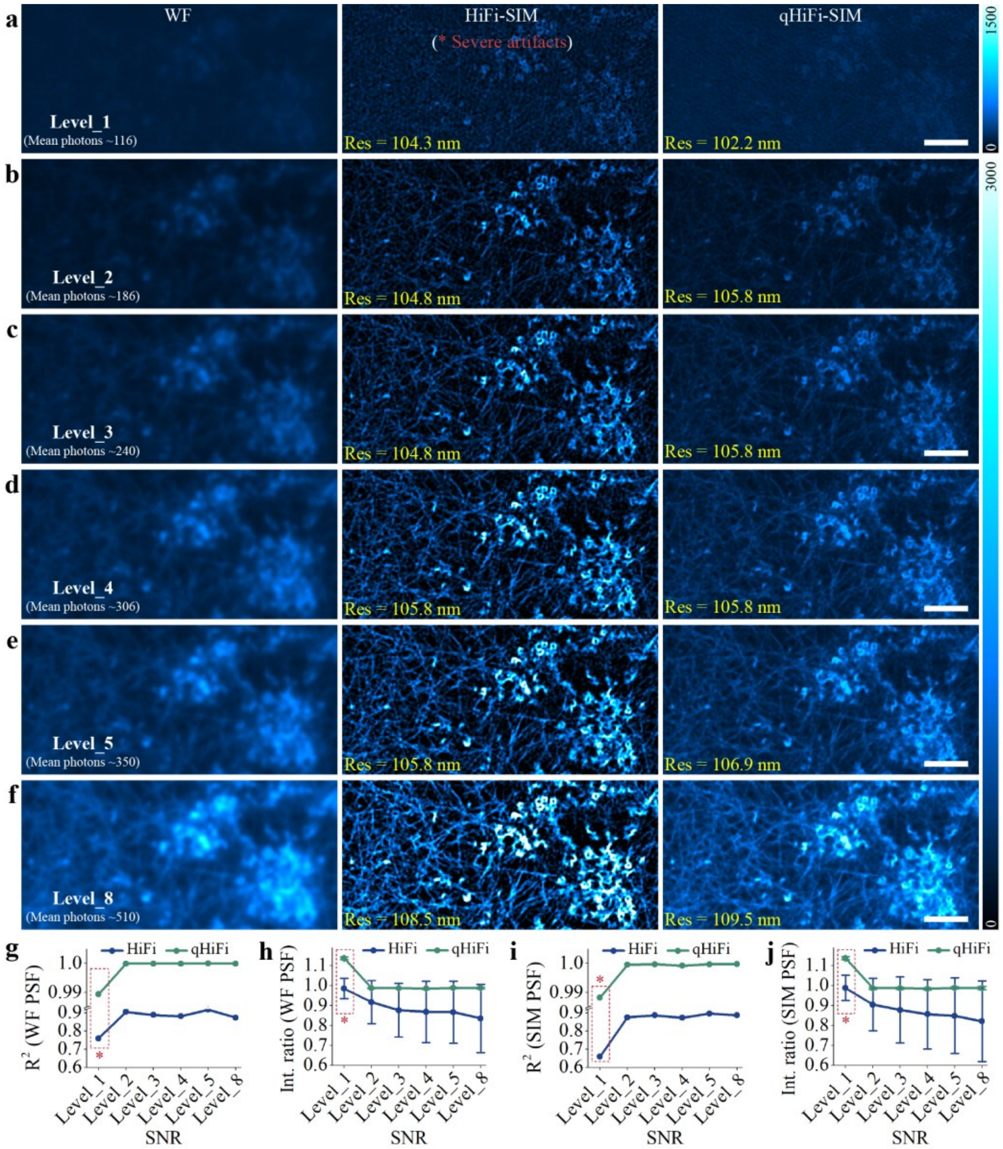


**Supplementary Fig. 22 | Quantitative evaluation of qHiFi-SIM reconstruction intensity linearity under varying signal-to-noise ratio (SNR) levels.** Test data from the F-actin sample in the BioSR dataset were analyzed at six different SNR levels (Level 1–5 and Level 8, with mean photon counts ranging from 116 to 510). **a**–**f**, WF (left), HiFi-SIM (middle), and qHiFi-SIM (right) reconstructions of F-actin structures at six SNR levels; the mean photon counts are annotated in the WF images, and spatial resolutions are indicated in the lower right of the SIM reconstructions. Severe artifacts are observed in the HiFi-SIM reconstruction of the raw data with low SNR (Level_1 group, labeled as "* Severe artifacts"). **g**–**j**, Quantitative comparison of $R^2$ values (**g**, **i**) and mean intensity ratios (**h**, **j**) between qHiFi-SIM and HiFi-SIM at the WF PSF scale (**g**, **h**) and SIM PSF scale (**i**, **j**). The results show that qHiFi-SIM effectively improves the intensity linearity of the initial HiFi-SIM reconstructions under Levels 2–5 and Level 8, with substantial improvements in key metrics (e.g., SSIM and PSNR) of the corresponding degraded WF images (**Supplementary Table 8**). Meanwhile, under Levels 2–5 and Level 8, qHiFi-SIM achieves stable $R^2$ values and mean intensity ratios, with an $R^2 > 0.999$ and a mean intensity ratio close to unity at the SIM PSF scale. Notably, at Level_1, the low SNR of the raw SIM data induced severe structure-related artifacts in the initial HiFi-SIM reconstruction, which ultimately resulted in abnormal $R^2$ and mean intensity metrics in the corresponding qHiFi-SIM reconstruction. Therefore, we recommend applying qHiFi-SIM for intensity linearity correction when the initial SIM image maintains high structural fidelity. For scenarios where the reconstruction itself inherently contains severe structural artifacts, intensity linearity correction is not recommended, as the structural fidelity is already compromised and discussing intensity fidelity loses its physical meaning. Scale bars: **a**–**f** 2 μm.

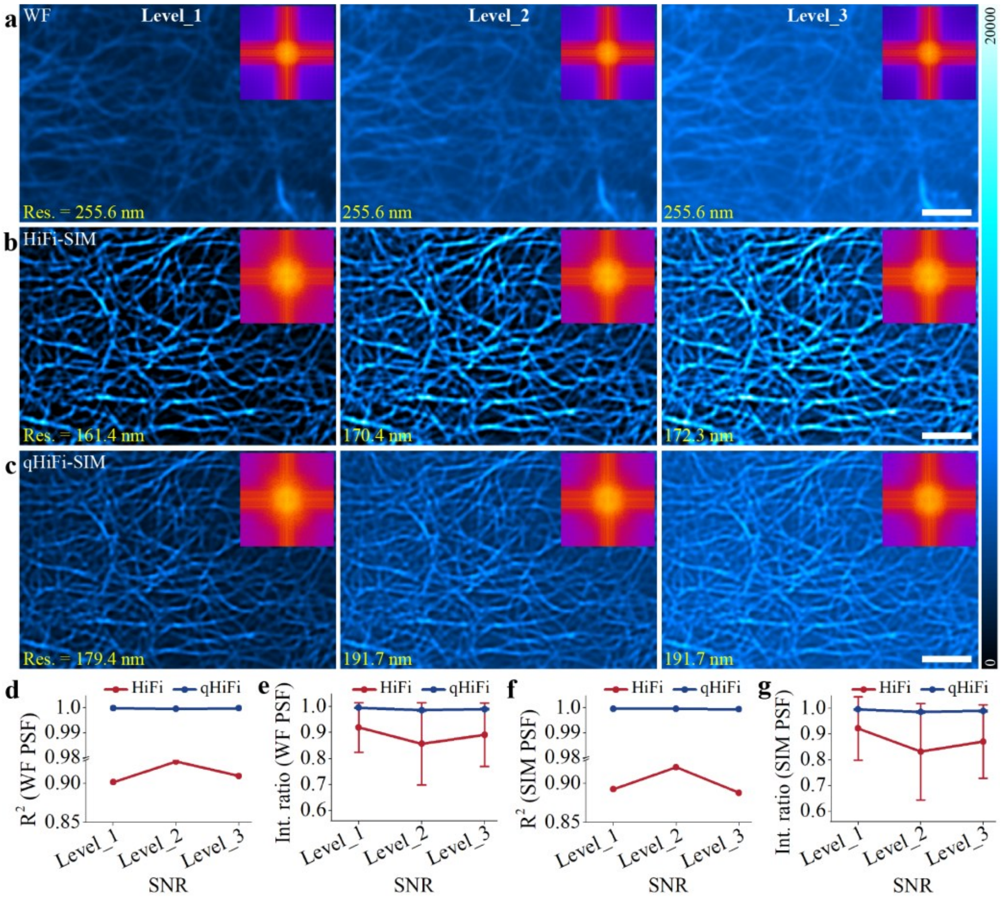


**Supplementary Fig. 23 | Quantitative evaluation of qHiFi-SIM reconstruction intensity linearity under varying out-of-focus background levels.** The SR image of the test F-actin sample from the BioSR dataset reconstructed by HiFi-SIM was utilized as the ground truth (GT). Structured illumination imaging was simulated under three out-of-focus background levels (Level_1–Level_3) with conditions matching those of the F-actin data used for network training. **a**–**c**, WF (**a**), HiFi-SIM (**b**), and qHiFi-SIM (**c**) reconstructions at the three out-of-focus background levels; Insets show the corresponding Fourier spectra, and spatial resolutions estimated by decorrelation analysis are annotated in the lower left. **d**–**g**, Quantitative comparison of $R^2$ values (**d**, **f**) and mean intensity ratios (**e**, **g**) between qHiFi-SIM and HiFi-SIM at the WF PSF scale (**d**, **e**) and SIM PSF scale (**f**, **g**). The results show that qHiFi-SIM consistently maintains stable and high $R^2$ values and mean intensity ratios close to unity across all out-of-focus levels, accompanied by substantial improvements in key metrics (e.g., SSIM and PSNR) of the corresponding degraded WF images (**Supplementary Table 9**). At the SIM PSF scale, it achieves an $R^2 > 0.999$ and mean intensity ratios ≥ 0.985, demonstrating excellent intensity linearity. However, as the out-of-focus background level increases, the high-frequency details in qHiFi-SIM reconstructions are progressively obscured by the background, leading to a shrinkage of the effective high-frequency range and a noticeable degradation in spatial resolution (**b**, **c**). This limitation arises because the qHiFi-SIM network was trained on high-quality BioSR datasets with low background signals, resulting in relatively constrained capability in preserving spatial resolution for strong out-of-focus background scenarios. Scale bars: **a**–**c** 2 μm.

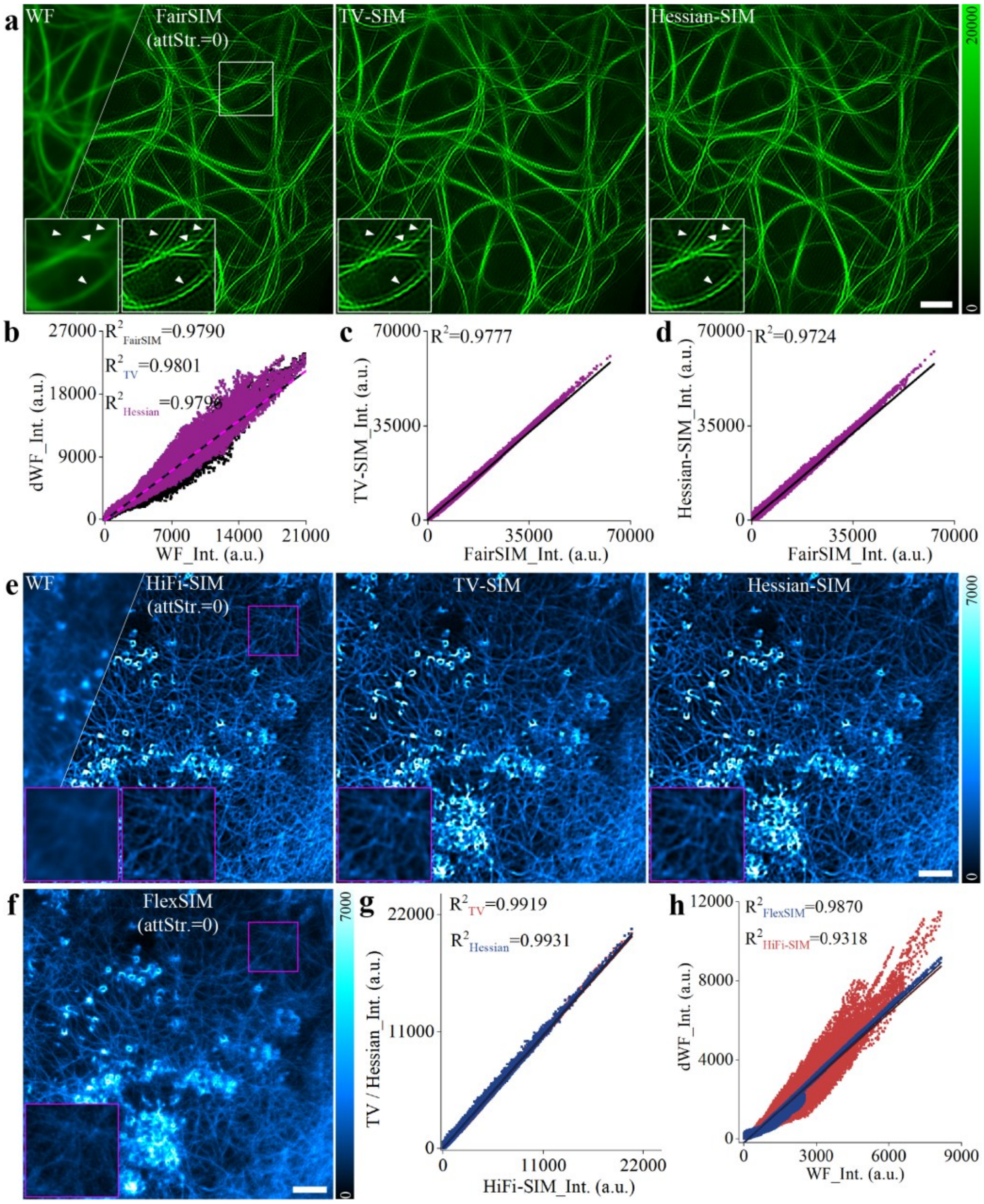


**Supplementary Fig. 24 | Evaluation of intensity linearity in regularized denoising and FlexSIM reconstruction. a**, WF (upper left), fairSIM reconstruction with OTF attenuation disabled (left), and its TV-regularized (TV-SIM; middle) and Hessian-regularized (Hessian-SIM; right) denoised reconstructions, showing that regularization cannot eliminate the sidelobe artifacts present in the input SIM image. **b–d**, $R^2$ analysis of the corresponding degraded WF images against the measured WF (**b**), and of TV-SIM (**c**) and Hessian-SIM (**d**) against fairSIM, indicating that the regularization processing has a relatively minor impact on intensity linearity. **e**, WF (upper left), HiFi-SIM reconstruction (left), and its TV-regularized (middle) and Hessian-regularized (right) denoised reconstructions. **f**, FlexSIM reconstruction with OTF attenuation disabled. **g**, $R^2$ analysis of TV-SIM and Hessian-SIM reconstructions against HiFi-SIM from **e**. **h**, $R^2$ analysis of the degraded WF images corresponding to HiFi-SIM from **e** and FlexSIM from **f** against the measured WF, showing that FlexSIM achieves better intensity linearity ($R^2$ = 0.9870) than HiFi-SIM ($R^2$ = 0.9318) when OTF attenuation is disabled. However, FlexSIM exhibits a lower SNR, and many fine structures clearly visible in the HiFi-SIM reconstruction are not resolved in FlexSIM. Scale bars: **a**, **e**, **f** 2 μm.

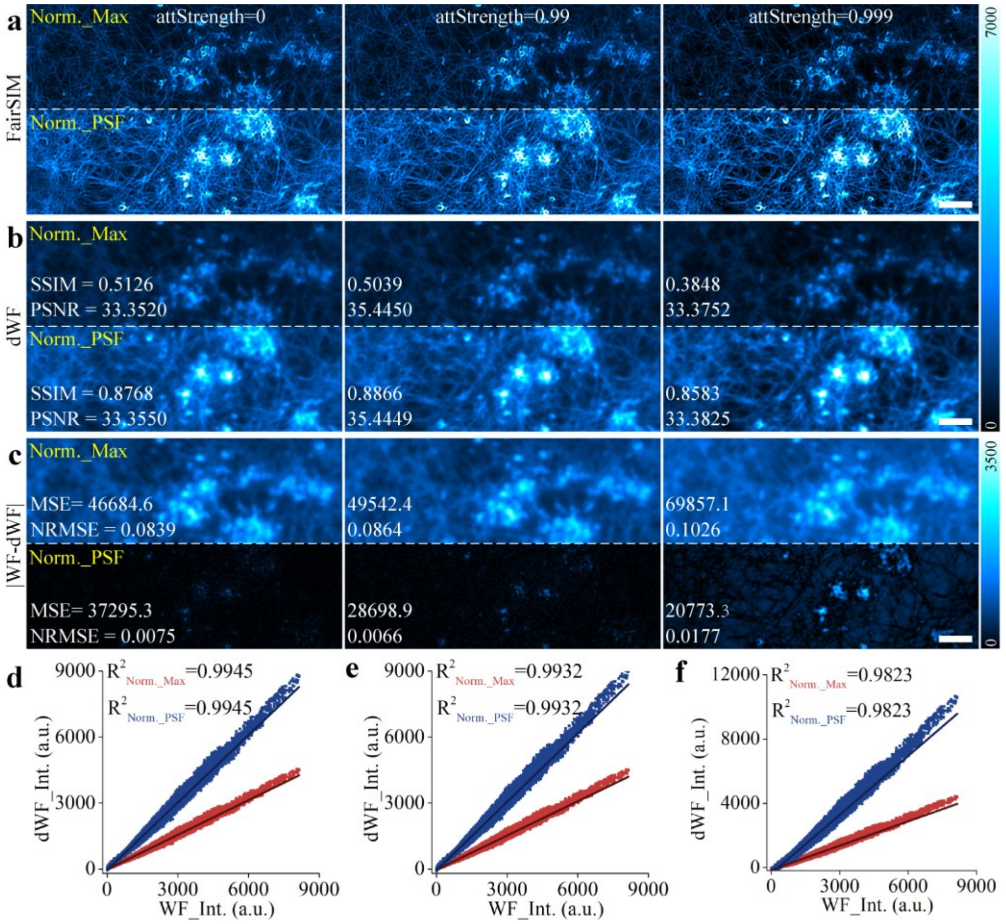


**Supplementary Fig. 25 | Effects of OTF attenuation and normalization methods on the intensity linearity of fairSIM reconstructions.** Using fairSIM reconstructions as an example, we evaluated the effects of OTF attenuation and two normalization methods, WF maximum intensity (Norm._Max) and PSF-convolution least-squares (Norm._PSF), on intensity linearity. **a**, fairSIM reconstructions processed with the two normalization methods at three OTF attenuation strengths (attStrength = 0, 0.99, and 0.999). **b**, Corresponding degraded WF images (dWF). **c**, Error maps between dWF and the measured WF. **d**–**f**, $R^2$ analysis between dWF and the measured WF at different OTF attenuation strengths. Intensity linearity progressively decreases as the OTF attenuation strength increases ($R^2$ dropping from 0.9945 to 0.9823), whereas the two normalization methods yield identical $R^2$ values across all attenuation strengths, indicating that normalization does not affect intensity linearity. Scale bars: **a**–**c** 2 μm.

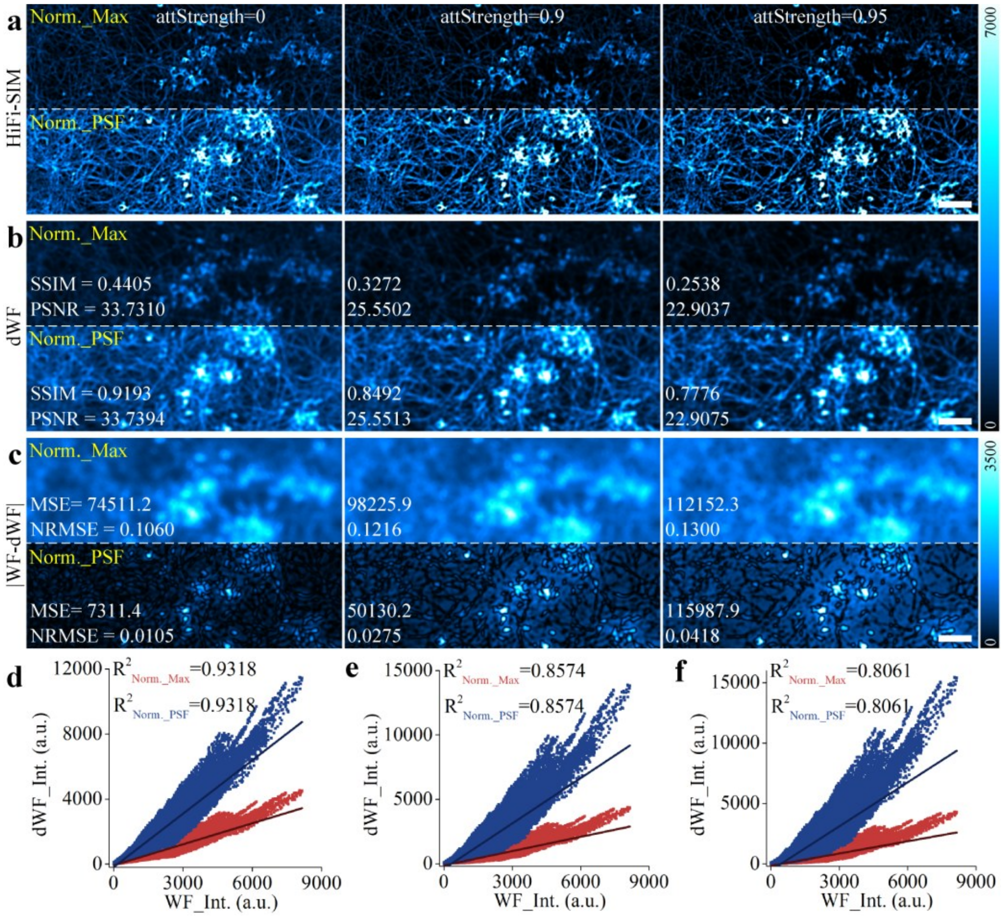


**Supplementary Fig. 26 | Effects of OTF attenuation and normalization methods on the intensity linearity of HiFi-SIM reconstructions.** Using HiFi-SIM reconstructions as an example, we evaluated the effects of OTF attenuation and two normalization methods, WF maximum intensity (Norm._Max) and PSF-convolution least-squares (Norm._PSF), on intensity linearity. **a**, HiFi-SIM reconstructions processed with the two normalization methods at three OTF attenuation strengths (attStrength = 0, 0.9, and 0.95). **b**, Corresponding degraded WF images (dWF). **c**, Error maps between dWF and the measured WF. **d**–**f**, $R^2$ analysis between dWF and the measured WF at different OTF attenuation strengths. Intensity linearity progressively decreases as the OTF attenuation strength increases ($R^2$ dropping from 0.9318 to 0.8061), whereas the two normalization methods yield identical $R^2$ values across all attenuation strengths, indicating that normalization does not affect intensity linearity. Scale bars: **a**–**c** 2 μm.

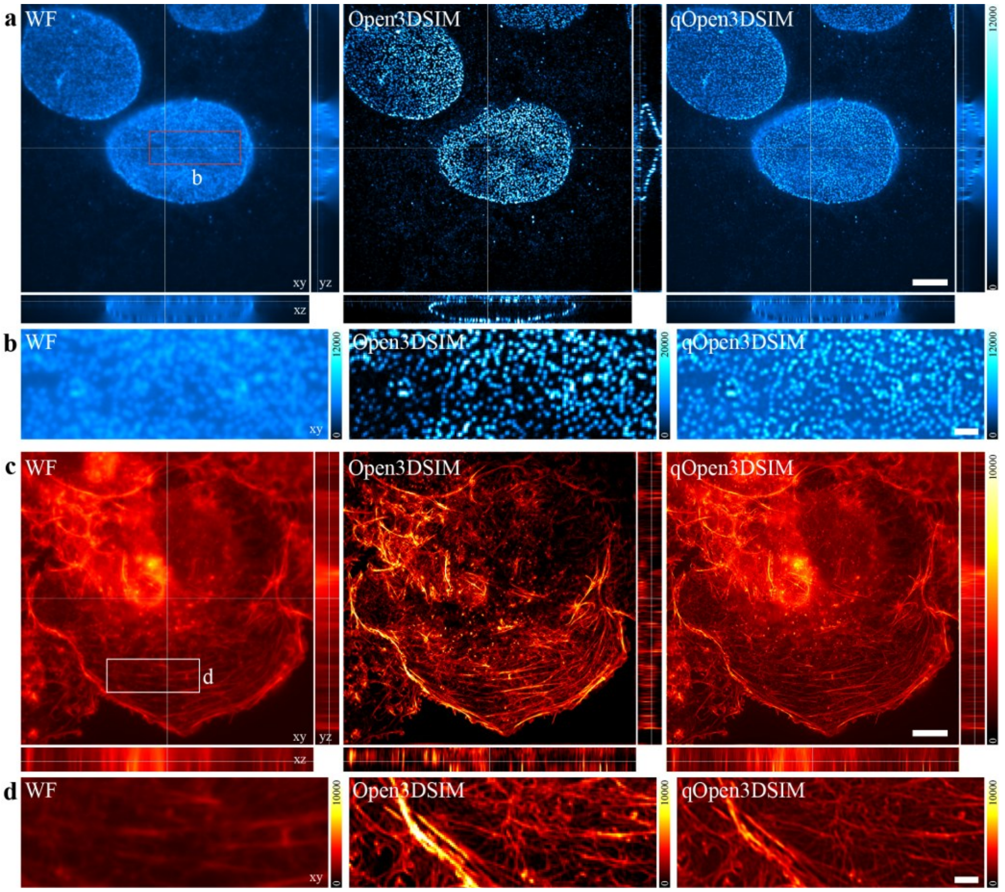


**Supplementary Fig. 27 | Evaluation of the limitations of qHiFi-SIM for intensity correction of full 3D-SIM data.** Nuclear pore data were obtained using Open3DSIM, whereas actin filament data in U2OS osteosarcoma cells were obtained using fairSIM. **a**, 3D WF (left), Open3DSIM (middle), and qOpen3DSIM (right) reconstructions of nuclear pore complexes. **b**, Magnified view of the red-boxed region in **a**. **c**, 3D WF (left), Open3DSIM (middle), and qOpen3DSIM (right) reconstructions of actin filaments. **d**, Magnified view of the white-boxed region in **c**. Since qHiFi-SIM uses the measured WF image as the intensity reference, and the WF image contains out-of-focus background that is removed by full 3D-SIM imaging, directly applying qHiFi-SIM for intensity correction reintroduces out-of-focus signals and thus compromises the optical sectioning of full 3D-SIM. Collectively, qHiFi-

SIM is not recommended for direct intensity correction of full 3D-SIM imaging. Scale bars: **a**, **c** 5 μm; **b, d** 1 μm.

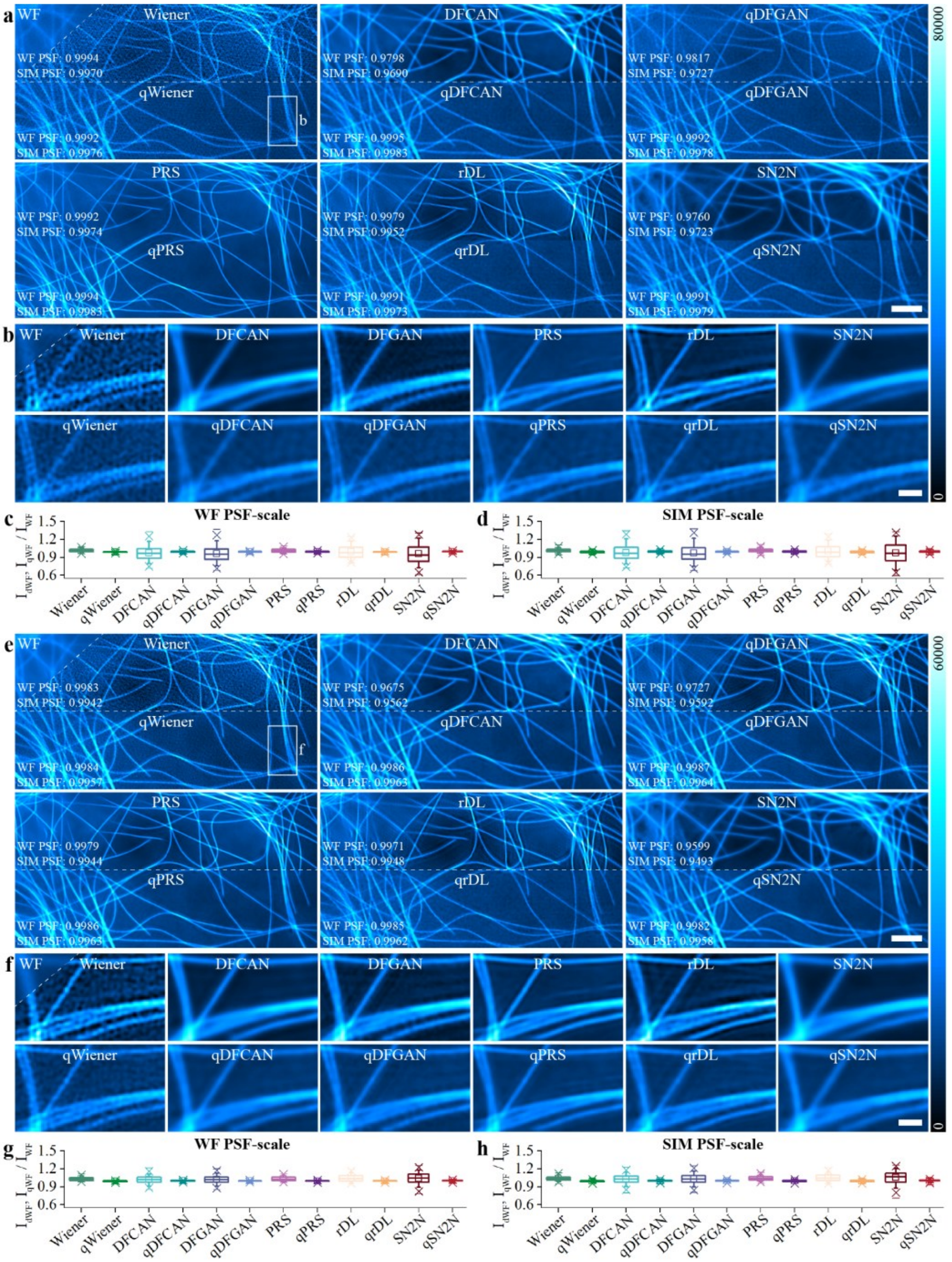


**Supplementary Fig. 28 | Comparison of reconstruction intensity linearity between state-of-the-art deep learning–based algorithms and qHiFi-SIM.** Microtubule data were obtained from the BioSR dataset. SIM images were reconstructed using the open-source standard Wiener-SIM algorithm from BF-SIM[16]. DFCAN and DFGAN[20], SN2N[22], and PRS[23] denoising models were then applied to denoise the initial SR images reconstructed by Wiener-SIM; and the rDL[21] model was applied to denoise and enhance the raw SIM data, followed by SR reconstruction using the same Wiener-SIM algorithm. The prefix "q" denotes the results further intensity-corrected by qHiFi-SIM. **a**, **e**, Reconstructions of low-SNR (**a**) and high-SNR (**e**) test data by the above algorithms and their qHiFi-SIM post-processed versions; $R^2$ values at the WF PSF and SIM PSF scales are annotated in the lower left of each image. **b**, **f**, Magnified views of the white-boxed regions in **a** and **e**, respectively. **c**, **d**, **g**, **h**, Box plots of mean intensity ratios for each algorithm and its post-processed version at the WF PSF scale (**c**, **g**) and SIM PSF scale (**d**, **h**). Although these deep learning-based methods inherit the favorable intensity linearity of the Wiener-SIM reconstructions used as the training datasets or the inputs to be denoised, subsequent qHiFi-SIM post-processing can still moderately improve their intensity linearity, achieving an $R^2 > 0.995$ and a mean intensity ratio ≥ 0.987 at the SIM PSF scale. Scale bars: **a**, **e** 2 μm; **b, f** 1 μm.

**Note:** The low-SNR data in **a** show an anomalously larger grayscale value range than the high-SNR data in **e**, which may result from a larger normalization factor applied to the low-SNR data, rather than indicating higher SNR. This accounts for the higher maximum value of the intensity color bar in **a** compared to **e**.

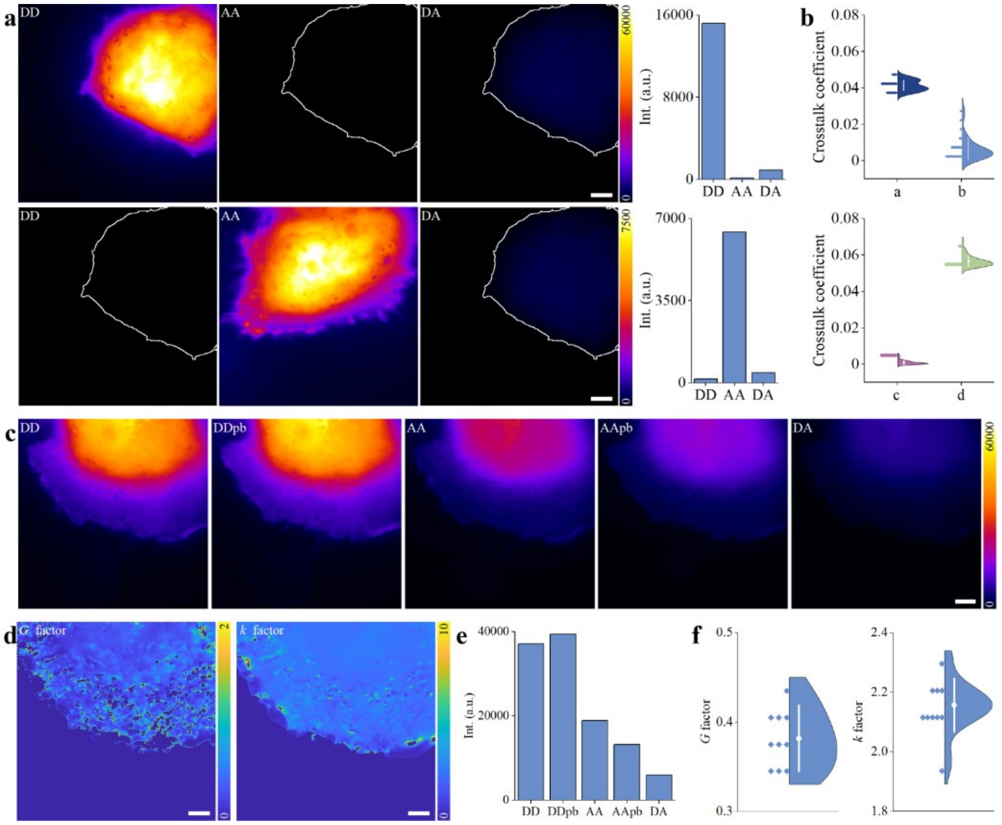


**Supplementary Fig. 29 | Experimental calibration of spectral crosstalk and *G*/*k* factors for quantitative FRET imaging. a**, Representative donor-only (top row) and acceptor-only (bottom row) control images acquired in the DD, AA, and DA channels; bar graphs on the right show the mean intensities of the corresponding channels. **b**, Distributions of the four spectral-crosstalk coefficients derived from the single-color controls. **c**, Representative donor–acceptor tandem construct images before and after partial acceptor photobleaching for mPb-G calibration. **d**, Representative pixel-wise *G*- and *k*-factor maps. **e**, Changes in DD and AA intensities used to determine the acceptor photobleaching fraction and donor dequenching. **f**, Distributions of the calibrated *G* and *k* factors from independent measurements. Scale bars: **a**, **c**, **d** 2 μm.

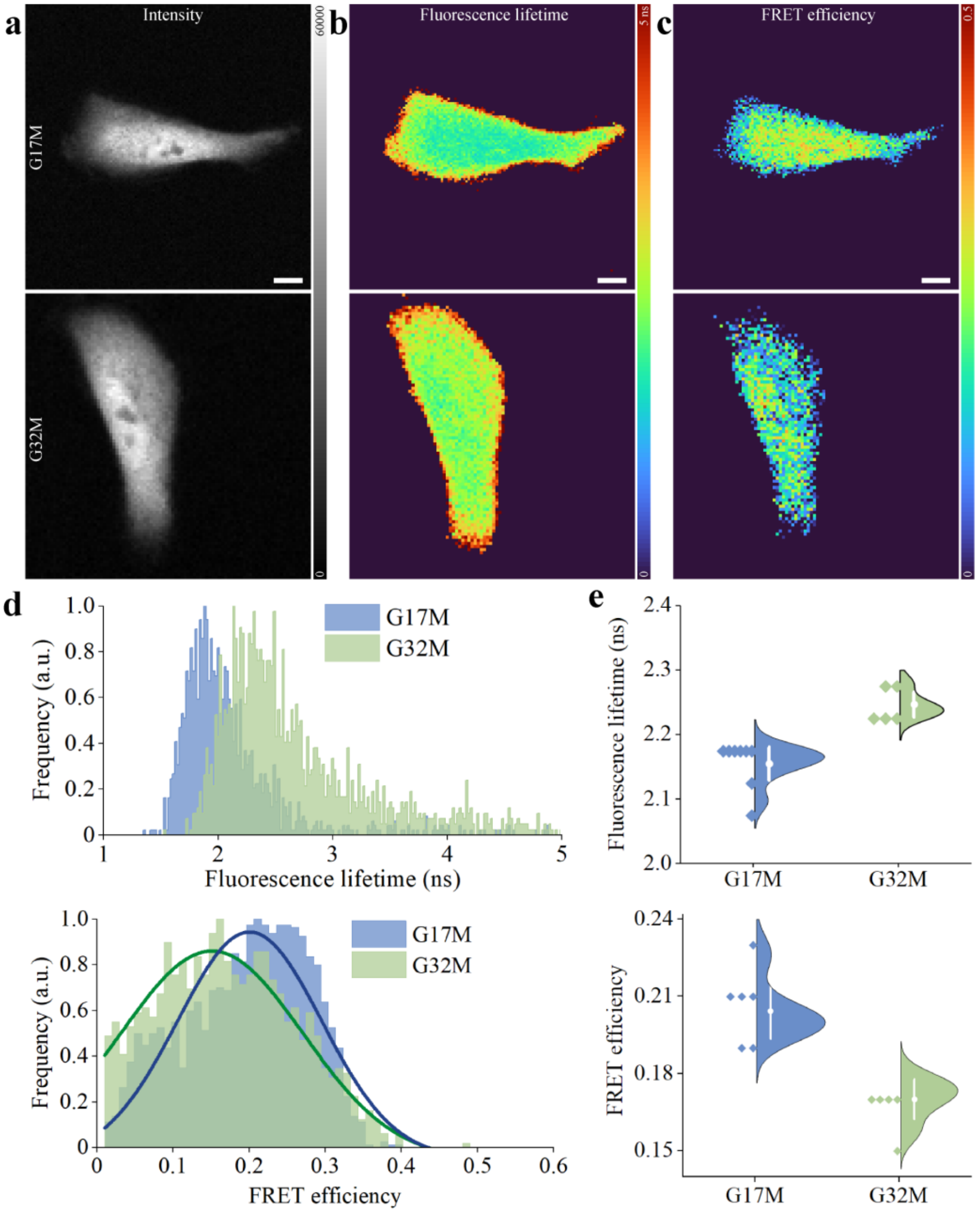


**Supplementary Fig. 30 | FLIM-FRET validation using reference FRET constructs in HeLa cells. a**, Representative donor-intensity images of G17M and G32M. **b**, **c**, Corresponding donor fluorescence-lifetime and FLIM-derived FRET-efficiency maps. **d**, Pixel-wise distributions of fluorescence lifetime and FRET efficiency for the representative cells. **e**, Cell-level distributions of mean donor lifetime and FLIM-derived FRET efficiency for G17M (*n* = 10) and G32M (*n* = 5). FLIM measurements were performed using a PicoQuant LSM Upgrade Kit coupled to a Zeiss LSM 780. Scale bars: **a–c** 8 μm.

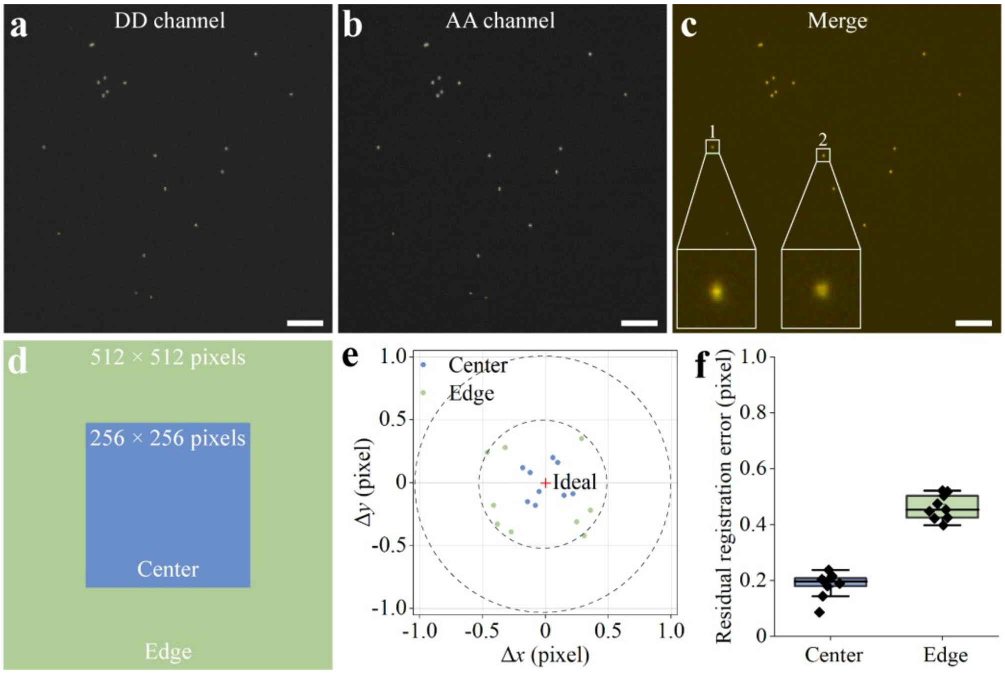


**Supplementary Fig. 31 | Validation of inter-channel registration accuracy. a–c**, Representative 100-nm TetraSpeck fluorescent beads (T7279, Thermo Fisher Scientific) acquired in the DD (**a**) and AA (**b**) channels, and their registered overlay (**c**). Insets 1 and 2 show magnified views of beads in the peripheral and central regions, respectively. **d**, Definition of the central (256 × 256 pixels) and peripheral regions used for field-dependent analysis. **e**, Residual lateral displacement $(\Delta x, \Delta y)$ of individual beads after affine registration. Blue and green dots denote beads in the central and peripheral regions, respectively; the red cross marks the ideal zero-displacement position. Dashed circles indicate radial errors of 0.5 and 1 pixel. **f**, Distribution of the residual registration error in the central and peripheral regions. All measured beads exhibited residual errors below one pixel. Scale bars: **a–c** 4 μm.

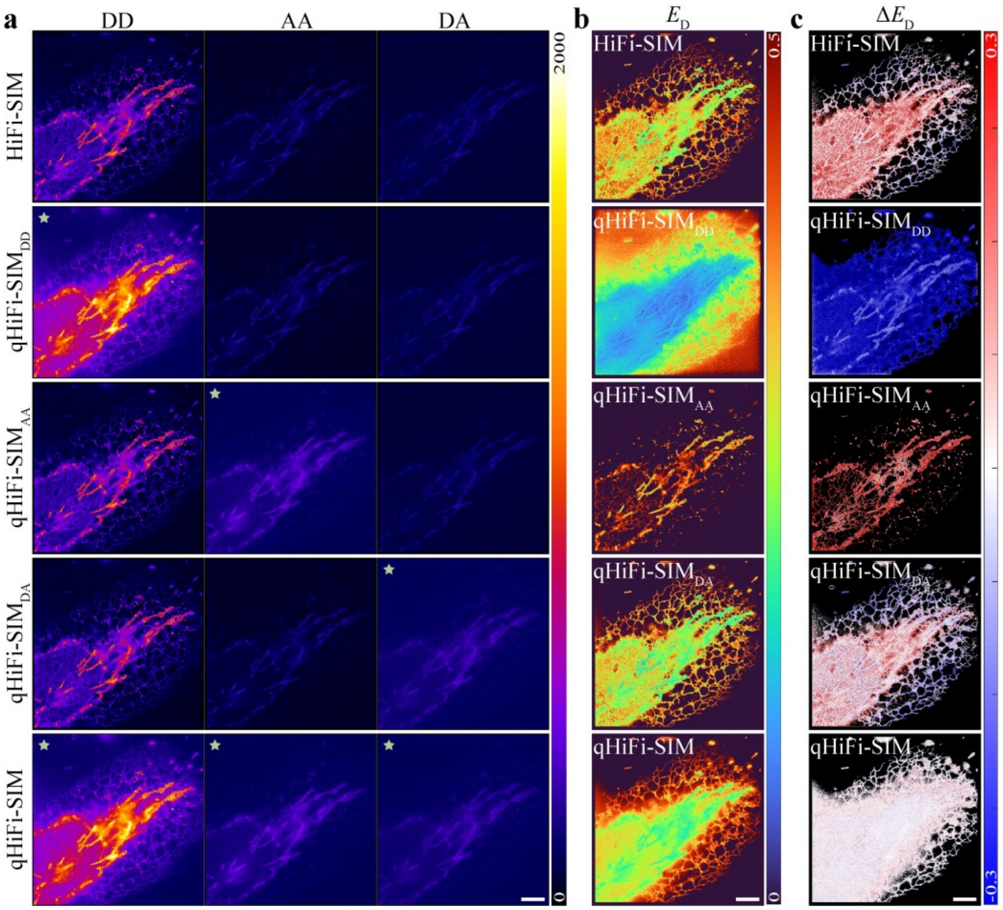


**Supplementary Fig. 32 | Channel-wise ablation of qHiFi-SIM processing for quantitative FRET.** The same FRET dataset as in **Fig. 5** was used. **a**, Representative HiFi-SIM and hybrid qHiFi-SIM images of DD, AA, and DA channels. In the hybrid combinations, qHiFi-SIM was applied individually to the DD, AA, or DA channel, or consistently to all three channels (bottom row). The stars indicate the channels processed with qHiFi-SIM. **b**, Corresponding donor-referenced FRET-efficiency ($E_\mathrm{D}$) maps calculated using the same fixed FRET calibration parameters. **c**, Pixel-wise deviations from the WF-derived $E_\mathrm{D}$ reference, defined as $\Delta E_\mathrm{D} = E_\mathrm{D}^\mathrm{method} - E_\mathrm{D}^\mathrm{WF}$. From top to bottom (HiFi-SIM, qHiFi-SIM$_\mathrm{DD}$, qHiFi-SIM$_\mathrm{AA}$, qHiFi-SIM$_\mathrm{DA}$, and qHiFi-SIM), the mean bias ± SD was 0.0436 ± 0.0623, −0.1552 ± 0.0379, 0.1726 ± 0.0613, 0.0130 ± 0.0605, and 0.0031 ± 0.0239, respectively. Scale bars: **a**–**c** 2 μm.

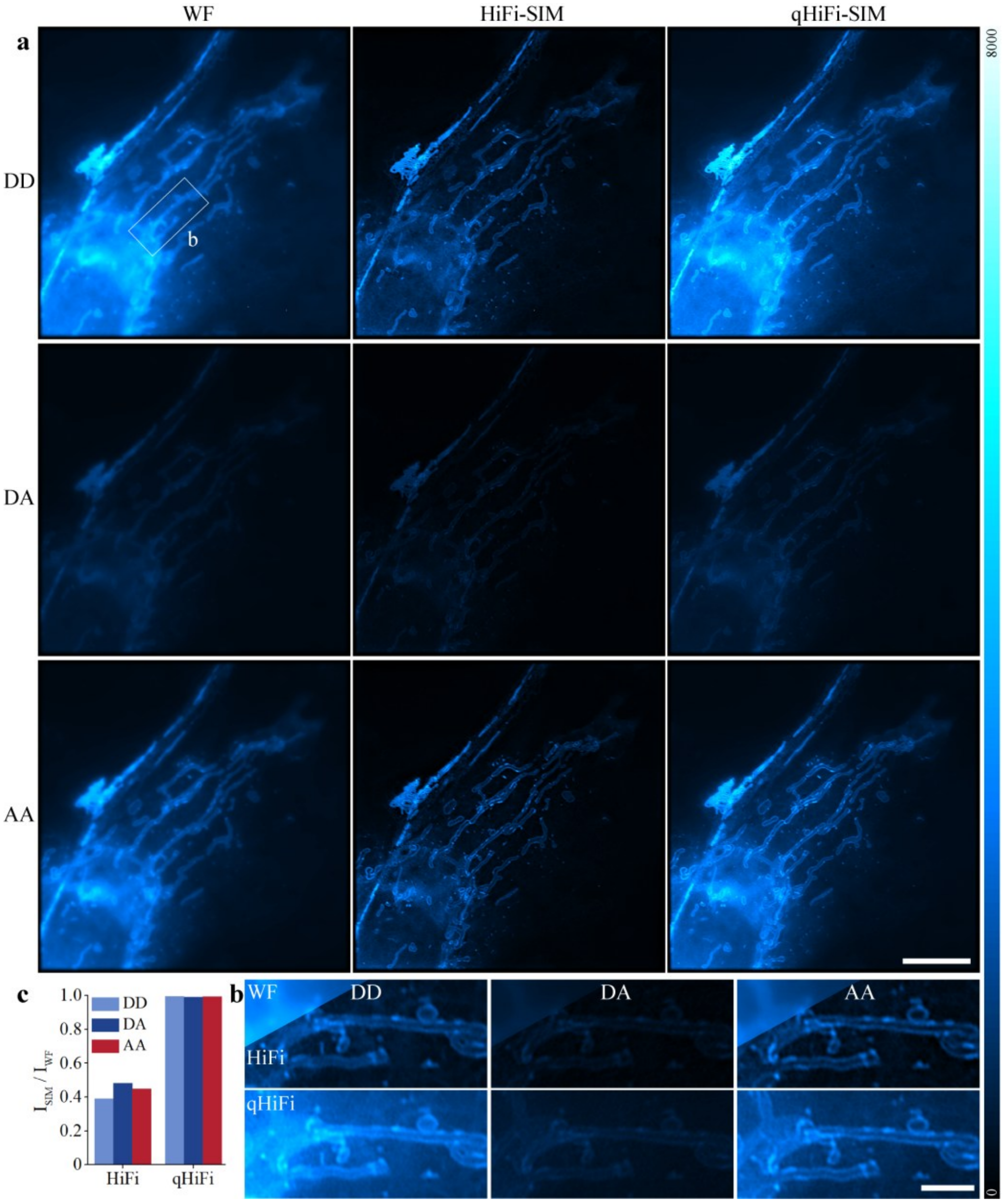


**Supplementary Fig. 33 | Large field-of-view SIM-FRET imaging data related to Fig. 6. a**, Data for the DD, DA, and AA channels are shown in rows (top to bottom). SR-SIM images were reconstructed using the HiFi-SIM and qHiFi-SIM algorithms. **b**, Magnified images of the white-box region in **a**. **c**, **d**, Mean intensity ratios of HiFi-SIM (with default OTF attenuation strength of 0.9), and qHiFi-SIM reconstructions relative to the corresponding WF images. HiFi-SIM reconstructions exhibited significant intensity deviations, yielding channel-specific mean intensity ratios of 0.39 (DD), 0.48 (DA), and 0.45 (AA). In contrast, qHiFi-SIM achieved excellent quantitative accuracy, with uniform mean intensity ratios of 0.99 across all channels. Scale bars: **a** 8 μm; **b** 2 μm.

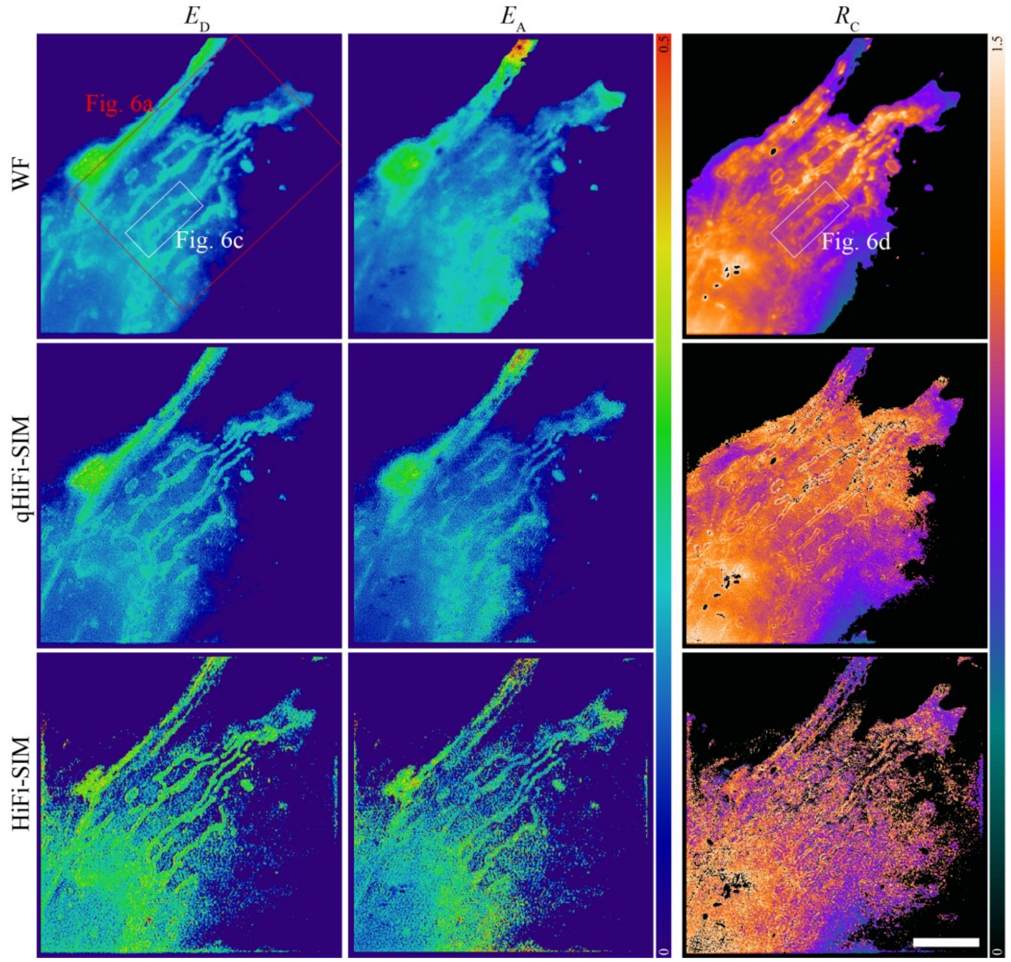


**Supplementary Fig. 34 | Quantitative analysis results of large field-of-view SIM-FRET imaging data related to Fig. 6.** Distributions of $E_D$, $E_A$, $R_C$ from FRET analysis are shown in rows (top to bottom), and corresponding images from WF, HiFi-SIM and qHiFi-SIM are displayed in columns (left to right). Scale bars: 8 μm.

| Sample No. | Resolution (nm) | |
|---|---|---|
| | **Base (qHiFi-SIM)** | **No SSFE** |
| 1 | 118.0 | 144.8 |
| 2 | 116.8 | 199.1 |
| 3 | 113.8 | 149.8 |
| 4 | 116.8 | 199.1 |
| 5 | 116.8 | 151.7 |
| 6 | 109.8 | 144.8 |
| 7 | 118.0 | 151.7 |
| 8 | 116.8 | 148.7 |
| 9 | 118.0 | 144.8 |
| **Mean ± Std** | **116.1 ± 2.7** | **159.4 ± 22.7** |

**Supplementary Table 1 | Ablation study of the SSFE branch: spatial resolution comparison.** Spatial resolution was estimated from the test dataset using decorrelation analysis. Base denotes the full qHiFi-SIM network with the SSFE branch; No SSFE indicates the network trained without the structural branch. Removal of the SSFE branch leads to a substantial degradation in spatial resolution, from 116.1 ± 2.7 nm (Base) to 159.4 ± 22.7 nm (No SSFE), accompanied by a marked increase in variability across independent test samples. These results demonstrate that the SSFE branch plays an essential role in preserving high-resolution structural information. Data are mean ± SD from $n$ = 9 independent test samples.

| Sample No. | SNR | |
|---|---|---|
| | **Base (qHiFi-SIM)** | **No WIFE** |
| 1 | 0.295 | 0.229 |
| 2 | 0.281 | 0.244 |
| 3 | 0.309 | 0.269 |
| 4 | 0.230 | 0.183 |
| 5 | 0.331 | 0.311 |
| 6 | 0.319 | 0.281 |
| 7 | 0.287 | 0.247 |
| 8 | 0.314 | 0.262 |
| 9 | 0.317 | 0.263 |
| **Mean ± Std** | **0.298 ± 0.030** | **0.254 ± 0.036** |

**Supplementary Table 2 | Ablation study of the WIFE branch: SNR comparison.** SNR was estimated from the test dataset using decorrelation analysis. Base denotes the full qHiFi-SIM network with the WIFE branch; No WIFE indicates the network trained without the intensity branch. Removal of the WIFE branch leads to a substantial reduction in SNR, from 0.298 ± 0.030 (Base) to 0.254 ± 0.036 (No WIFE), corresponding to a relative decrease of approximately 14.8%. These results demonstrate that the WIFE branch plays a critical role in preserving signal-to-noise ratio and intensity fidelity. Data are mean ± SD from $n$ = 9 independent test samples.

| Sample No. | Base (qHiFi-SIM) | | No WIFE | |
|---|---|---|---|---|
| | $R^2$ (WF PSF-scale) | $R^2$ (SIM PSF-scale) | $R^2$ (WF PSF-scale) | $R^2$ (SIM PSF-scale) |
| 1 | 0.9994 | 0.9989 | 0.9617 | 0.9600 |
| 2 | 0.9999 | 0.9989 | 0.9757 | 0.9704 |
| 3 | 0.9999 | 0.9998 | 0.9749 | 0.9753 |
| 4 | 0.9999 | 0.9999 | 0.9726 | 0.9723 |
| 5 | 0.9990 | 0.9990 | 0.9766 | 0.9777 |
| 6 | 0.9998 | 0.9997 | 0.9825 | 0.9818 |
| 7 | 0.9999 | 0.9998 | 0.9783 | 0.9800 |
| 8 | 0.9989 | 0.9987 | 0.9716 | 0.9690 |
| 9 | 0.9997 | 0.9998 | 0.9678 | 0.9691 |

**Supplementary Table 3 | Ablation study of the WIFE branch: $R^2$ comparison.** $R^2$ values were computed between the degraded WF image (SIM reconstruction convolved with the PSF) and the measured WF image, evaluated at both the WF PSF scale and the SIM PSF scale. $R^2$ values at each scale are derived from 500 randomly selected ROIs. Base denotes the full qHiFi-SIM network with the WIFE branch; No WIFE indicates the network trained without the intensity branch. The base model achieves consistently high $R^2$ values (> 0.998) across all samples and both PSF scales. In contrast, removal of the WIFE branch leads to a substantial reduction in $R^2$, with values dropping to approximately 0.960–0.983, indicating a marked loss of intensity fidelity. These results demonstrate that the WIFE branch is essential for maintaining quantitative intensity accuracy in qHiFi-SIM reconstructions. Data are from $n$ = 9 independent test samples.

| Sample No. | PSNR | | SSIM | | MSE | | NRMSE | |
|---|---|---|---|---|---|---|---|---|
| | **Base (qHiFi-SIM)** | **No WIFE** | **Base (qHiFi-SIM)** | **No WIFE** | **Base (qHiFi-SIM)** | **No WIFE** | **Base (qHiFi-SIM)** | **No WIFE** |
| 1 | 47.5927 | 30.5429 | 0.9953 | 0.9514 | 1155.4 | 58575.3 | 0.0042 | 0.0297 |
| 2 | 47.2499 | 28.1118 | 0.9936 | 0.9554 | 205.3 | 16837.0 | 0.0043 | 0.0393 |
| 3 | 48.2155 | 33.5767 | 0.9938 | 0.9603 | 281.1 | 8181.3 | 0.0039 | 0.0209 |
| 4 | 46.4268 | 25.1402 | 0.9935 | 0.9571 | 265.9 | 35759.0 | 0.0048 | 0.0553 |
| 5 | 47.7719 | 32.5748 | 0.9932 | 0.9458 | 463.2 | 15329.3 | 0.0041 | 0.0235 |
| 6 | 47.4636 | 33.5027 | 0.9936 | 0.9596 | 469.1 | 11678.1 | 0.0042 | 0.0211 |
| 7 | 46.0533 | 26.1269 | 0.9938 | 0.9475 | 641.0 | 63019.7 | 0.0050 | 0.0494 |
| 8 | 47.1666 | 29.1028 | 0.9900 | 0.9077 | 844.4 | 54065.6 | 0.0044 | 0.0351 |
| 9 | 45.9521 | 28.9121 | 0.9938 | 0.9293 | 742.2 | 37540.9 | 0.0050 | 0.0358 |

**Supplementary Table 4 | Ablation study of the WIFE branch: PSNR, SSIM, MSE, NRMSE comparison.** Quantitative image quality metrics (PSNR, SSIM, MSE, and NRMSE) were computed between the degraded WF image (SIM reconstruction convolved with the PSF) and the measured WF image on the test dataset. Base denotes the full qHiFi-SIM network with the WIFE branch; No WIFE indicates the network trained without the intensity branch. The base model consistently outperforms the No WIFE version across all metrics and samples. Specifically, PSNR decreases from 47.0 ± 0.8 dB (Base) to 29.8 ± 3.1 dB (No WIFE), SSIM drops from 0.993 ± 0.002 to 0.946 ± 0.017, MSE increases from 565.3 ± 350.3 to 33,665.5 ± 20,321.5, and NRMSE rises from 0.0044 ± 0.0004 to 0.0348 ± 0.0121. These results demonstrate that the WIFE branch is critical for maintaining reconstruction fidelity and suppressing reconstruction errors. Data are mean ± SD from $n$ = 9 independent test samples.

| Sample No. | Base (qHiFi-SIM) | | No Spatial Attention | |
|---|---|---|---|---|
| | Resolution (nm) | SNR | Resolution (nm) | SNR |
| 1 | 118.0 | 0.295 | 118.0 | 0.288 |
| 2 | 116.8 | 0.281 | 118.0 | 0.279 |
| 3 | 113.8 | 0.309 | 116.8 | 0.304 |
| 4 | 116.8 | 0.230 | 116.8 | 0.229 |
| 5 | 116.8 | 0.331 | 118.0 | 0.327 |
| 6 | 109.8 | 0.319 | 113.8 | 0.312 |
| 7 | 118.0 | 0.287 | 119.4 | 0.284 |
| 8 | 116.8 | 0.314 | 116.8 | 0.296 |
| 9 | 118.0 | 0.317 | 118.0 | 0.313 |

**Supplementary Table 5 | Ablation study of the Spatial Attention mechanism: spatial resolution and SNR comparison.** Spatial resolution and SNR were estimated from the test dataset using decorrelation analysis. Base denotes the full qHiFi-SIM network with the spatial attention mechanism; No spatial attention indicates the network trained without this module. The base model achieves slightly better resolution preservation (116.1 ± 2.9 nm) compared to the model without spatial attention (117.3 ± 1.6 nm), while SNR shows a more pronounced reduction upon removal of the module, decreasing from 0.298 ± 0.030 to 0.292 ± 0.031. These results suggest that the spatial attention module contributes to both spatial resolution maintenance and intensity fidelity, with a more notable effect on SNR preservation. Data are mean ± SD from $n$ = 9 independent test samples.

| Sample No. | SNR | |
|---|---|---|
| | Base (qHiFi-SIM) | No Identity |
| 1 | 0.295 | 0.292 |
| 2 | 0.281 | 0.279 |
| 3 | 0.309 | 0.306 |
| 4 | 0.230 | 0.227 |
| 5 | 0.331 | 0.329 |
| 6 | 0.319 | 0.315 |
| 7 | 0.287 | 0.285 |
| 8 | 0.314 | 0.306 |
| 9 | 0.317 | 0.314 |

**Supplementary Table 6 | Ablation study of the identity skip connection: SNR comparison.** SNR was estimated from the test dataset using decorrelation analysis. Base denotes the full qHiFi-SIM network with identity skip connections in the residual blocks; No identity indicates the network trained without these connections. Removal of the identity skip connection leads to a modest but consistent reduction in SNR across all test samples, from 0.298 ± 0.030 (Base) to 0.295 ± 0.031 (No identity). While this effect size is modest and falls within the range of measurement variance, the consistent directional shift across all test samples suggests that the skip connection primarily contributes to training stability and noise robustness. This is likely because it introduces the initial high-SNR SIM reconstruction as a structural prior, which facilitates stable feature propagation. Data are mean ± SD from $n = 9$ independent test samples.

| $\lambda_1, \lambda_2, \lambda_3$ | **$R^2$** (qWF vs WF) | | **PSNR** | **SSIM** | **MSE** | **NRMSE** | Notes |
|---|---|---|---|---|---|---|---|
| | WF PSF-scale | SIM PSF-scale | | | | | |
| 0.1, 0.9, 0.9 | 0.9533 | 0.9395 | 18.4728 | 0.6803 | 394200.2 | 0.1192 | |
| 0.8, 0.9, 0.1 | 0.9925 | 0.9887 | 23.8421 | 0.9347 | 114495.1 | 0.0643 | |
| 0.8, 0.1, 0.9 | 0.9575 | 0.9454 | 17.5978 | 0.5745 | 482189.5 | 0.1319 | |
| 0.5, 0.5, 0.5 | 0.9630 | 0.9570 | 19.5112 | 0.7448 | 310365.7 | 0.1058 | |
| **0.5, 0.1, 0.1** | **0.9999** | **0.9999** | **47.0851** | **0.9909** | **542.6** | **0.0044** | **Optimal** |
| 0.5, 0.1, 0 | 0.9999 | 0.9999 | 49.9245 | 0.9948 | 282.2 | 0.0032 | Severe artifacts |
| 0.5, 0, 0 | 0.9999 | 0.9998 | 51.1108 | 0.9955 | 214.7 | 0.0028 | Severe artifacts |

**Supplementary Table 7 | Performance metrics of qHiFi-SIM under different combinations of loss weights ($\lambda_1$, $\lambda_2$, $\lambda_3$).** See **Supplementary Note 3** for the roles and setting guidelines of these weights.

| Noise level | PSNR | | SSIM | | MSE | | NRMSE | | Notes |
|---|---|---|---|---|---|---|---|---|---|
| | Before correction | After correction | Before correction | After correction | Before correction | After correction | Before correction | After correction | |
| Level_01 | 22.3825 | 22.3020 | 0.7375 | **0.8909** | 257.2 | 262.1 | 0.0760 | **0.0767** | Severe artifacts |
| Level_02 | 24.6674 | **41.5617** | 0.7912 | **0.9642** | 1671.9 | **34.2** | 0.0584 | **0.0084** | |
| Level_03 | 24.9994 | **42.9062** | 0.8038 | **0.9747** | 3508.3 | **56.8** | 0.0562 | **0.0072** | |
| Level_04 | 24.9311 | **43.2736** | 0.8061 | **0.9797** | 6420.7 | **94.0** | 0.0567 | **0.0069** | |
| Level_05 | 25.0828 | **44.3171** | 0.8079 | **0.9827** | 9074.5 | **108.2** | 0.0557 | **0.0061** | |
| Level_08 | 25.4596 | **45.5449** | 0.8141 | **0.9883** | 21546.9 | **211.3** | 0.0533 | **0.0053** | |

**Supplementary Table 8 | Quantitative comparison of image quality metrics between the degraded WF and measured WF before and after qHiFi-SIM correction at different noise levels.**

| Background level | PSNR | | SSIM | | MSE | | NRMSE | |
|---|---|---|---|---|---|---|---|---|
| | Before correction | After correction | Before correction | After correction | Before correction | After correction | Before correction | After correction |
| Level_1 | 24.1131 | **45.6041** | 0.8296 | **0.9953** | 448596.0 | **3182.4** | 0.0623 | **0.0052** |
| Level_2 | 22.8801 | **45.3552** | 0.8172 | **0.9965** | 749969.5 | **4241.6** | 0.0718 | **0.0054** |
| Level_3 | 22.5098 | **44.7926** | 0.8101 | **0.9969** | 1003830.3 | **5934.4** | 0.0749 | **0.0058** |

**Supplementary Table 9 | Quantitative comparison of image quality metrics between degraded WF and measured WF before and after qHiFi-SIM correction under different background levels.**

| Split | BioSR group IDs | Number of samples |
|---|---|---|
| Training | Cell_001, Cell_002, Cell_006, Cell_007, Cell_009, Cell_010, Cell_012, Cell_013, Cell_014, Cell_015, Cell_018, Cell_019, Cell_020, Cell_022, Cell_023, Cell_024 (two sub-images: Cell_024_1, Cell_024_2), Cell_025, Cell_026, Cell_027, Cell_028, Cell_029, Cell_030, Cell_031, Cell_032, Cell_033, Cell_034, Cell_035, Cell_036, Cell_037, Cell_038, Cell_039, Cell_040, Cell_041, Cell_042 | 35 |
| Validation | Cell_043, Cell_044, Cell_046, Cell_047, Cell_048, Cell_049, Cell_050, Cell_051 | 8 |
| Testing | Cell_003, Cell_004, Cell_005, Cell_008, Cell_011, Cell_016, Cell_017, Cell_021, Cell_045 | 9 |

**Supplementary Table 10 | Exact sample identifiers for the training, validation, and test splits.** The training set comprises 34 original BioSR groups; Cell_024 contributes two sub-images, yielding 35 training samples in total. The validation set contains 8 samples, and the test set contains 9 samples. The exact split can be reproduced directly from the table.

| System | $\lambda_{em}$ (nm) | NA | SIM pixel size (nm) | $r_{WF}$ (nm) | WF ROI size (pixels) | $r_{SIM}$ (nm) | SIM ROI size (pixels) |
|---|---|---|---|---|---|---|---|
| BioSR F-actin (training data) | 525 | 1.41 | 31.3 | 227.1 | **17 × 17** | 113.6 | **9 × 9** |
| Polar-SIM (Airy Technologies) | 525 | 1.49 | 32.5 | 214.9 | **15 × 15** | 107.5 | **9 × 9** |
| Polar-SIM (Airy Technologies) | 605 | 1.49 | 32.5 | 247.7 | **17 × 17** | 123.8 | **9 × 9** |
| Home-built SIM-FRET | 600 | 1.49 | 36.1 | 245.6 | **15 × 15** | 122.8 | **9 × 9** |
| GE Healthcare (DeltaVision OMX V4) | 525 | 1.40 | 39.75 | 228.8 | **13 × 13** | 114.4 | **7 × 7** |
| GE Healthcare (DeltaVision OMX SR) | 527 | 1.42 | 39.3 | 226.4 | **13 × 13** | 113.2 | **7 × 7** |
| 3I-SIM | 610 | 1.49 | 32.5 | 249.7 | **17 × 17** | 124.9 | **9 × 9** |

**Supplementary Table 11 | PSF-scale ROI sizes for mean intensity ratio analysis across imaging systems.**
